# Regional specialization, polyploidy, and seminal fluid transcripts in the Drosophila female reproductive tract

**DOI:** 10.1101/2023.10.05.561141

**Authors:** Rachel C. Thayer, Elizabeth S. Polston, Jixiang Xu, David J. Begun

**Affiliations:** Department of Evolution and Ecology, University of California, Davis, CA, USA

**Keywords:** spermatheca, seminal receptacle, gene expression, marker gene, seminal fluid protein, uterus, oviduct, parovaria, accessory gland, secretion, endoreplication

## Abstract

Internal fertilization requires the choreographed interaction of female cells and molecules with seminal fluid and sperm. In many animals, including insects, the female reproductive tract is physically subdivided into sections that carry out specialized functions. For example, females of many species have specialized organs for sperm storage. *Drosophila melanogaster* is a premier model system for investigating many aspects of animal reproduction. Nevertheless, in contrast to males, much of the basic biology of the *D. melanogaster* female reproductive tract remains poorly understood or completely unknown. Here we use single-cell RNA-seq data and in situ hybridization to reveal a rich and previously unknown female reproductive tract cell diversity, including widespread variation in ploidy levels. We find that many so-called seminal fluid protein genes appear to be transcribed in specialized cells of the female reproductive tract, motivating a re-evaluation of the functional and evolutionary biology of this major class of proteins.

## INTRODUCTION

The *Drosophila* female reproductive tract is morphologically and functionally complex. Below the ovaries, the lower reproductive tract (FRT) is composed of 5 morphologically distinct organs (Fig. 1A-B) with various critical functions. The uterus (UT) is a muscular organ that changes conformation to transit eggs and allow sperm to access the storage organs (Mattei et al. 2015; Adams and Wolfner 2007). To maneuver sperm and eggs, the uterine wall has functionally distinctive regions. These include the oviduct valve flap (OVF), which is closed prior to mating, and which can block access to the oviduct and sperm storage organs, and the specialized vaginal intima (SVI), a region in the lower uterine wall. The uterus is also the site of fertilization and mating plug formation, and is the first tissue contacted by male products during copulation. The oviduct (OV) transfers eggs into the uterus in response to octopaminergic neural signaling (Rubinstein and Wolfner 2013). Sperm from separate males compete for occupancy of the seminal receptacle (SR), which stores the sperm set primarily used for fertilization (Manier et al. 2010). The seminal receptacle is a long, coiled, tubular organ and has two morphologically distinct regions: a narrow proximal section and a contrastingly thicker distal section, which also differ in the arrangement of microvilli on the inner surface, the density of secretory vacuoles, and the postures adopted by occupying sperm within each section (Y. Heifetz and Rivlin 2010; Mayhew and Merritt 2013). The paired spermathecae (ST) are sclerotized, glandular capsules that also store sperm. The spermathecae are capped by secretory cells that are required for fertilization and ovulation (Schnakenberg, Matias, and Siegal 2011; Sun and Spradling 2013), as are the secretions of the bilaterally paired parovaria (PV, (Sun and Spradling 2012), also known as the female accessory glands. A reproductive-associated fat body (FB) is found agglomerated over the 4 glands (Fig. 2C).

**Figure 1:**
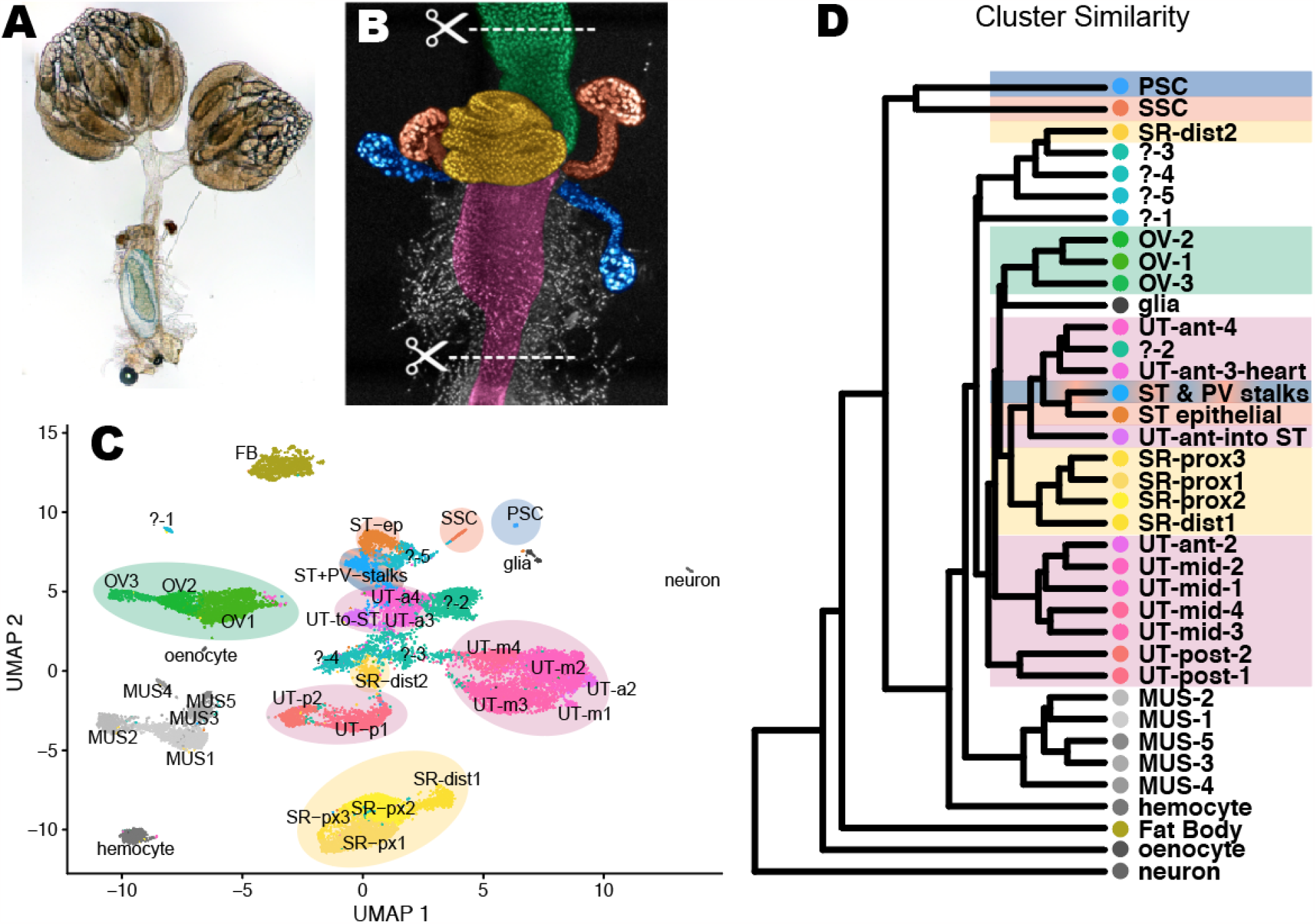
Organs and cell types in the somatic female reproductive tract. *A)* The complete *Drosophila* female reproductive tract, including the ovaries and a descended egg held in the uterus. *B)* Micrograph of the sequenced region of the lower FRT, with a DAPI stain and overlay colors showing organs: green = oviduct (OV), yellow = seminal receptacle (SR), orange = spermathecae (ST), blue = parovaria (PV, also called female accessory glands), and magenta = uterus (UT). *C)* Annotated UMAP plot of all sequenced nuclei. ‘SSC’ = spermathecal secretory cells. ‘PSC’ = parovaria secretory cells. ‘MUS’ = muscle. *D)* Cluster similarity, based on gene expression principal components analysis.

**Figure 2:**
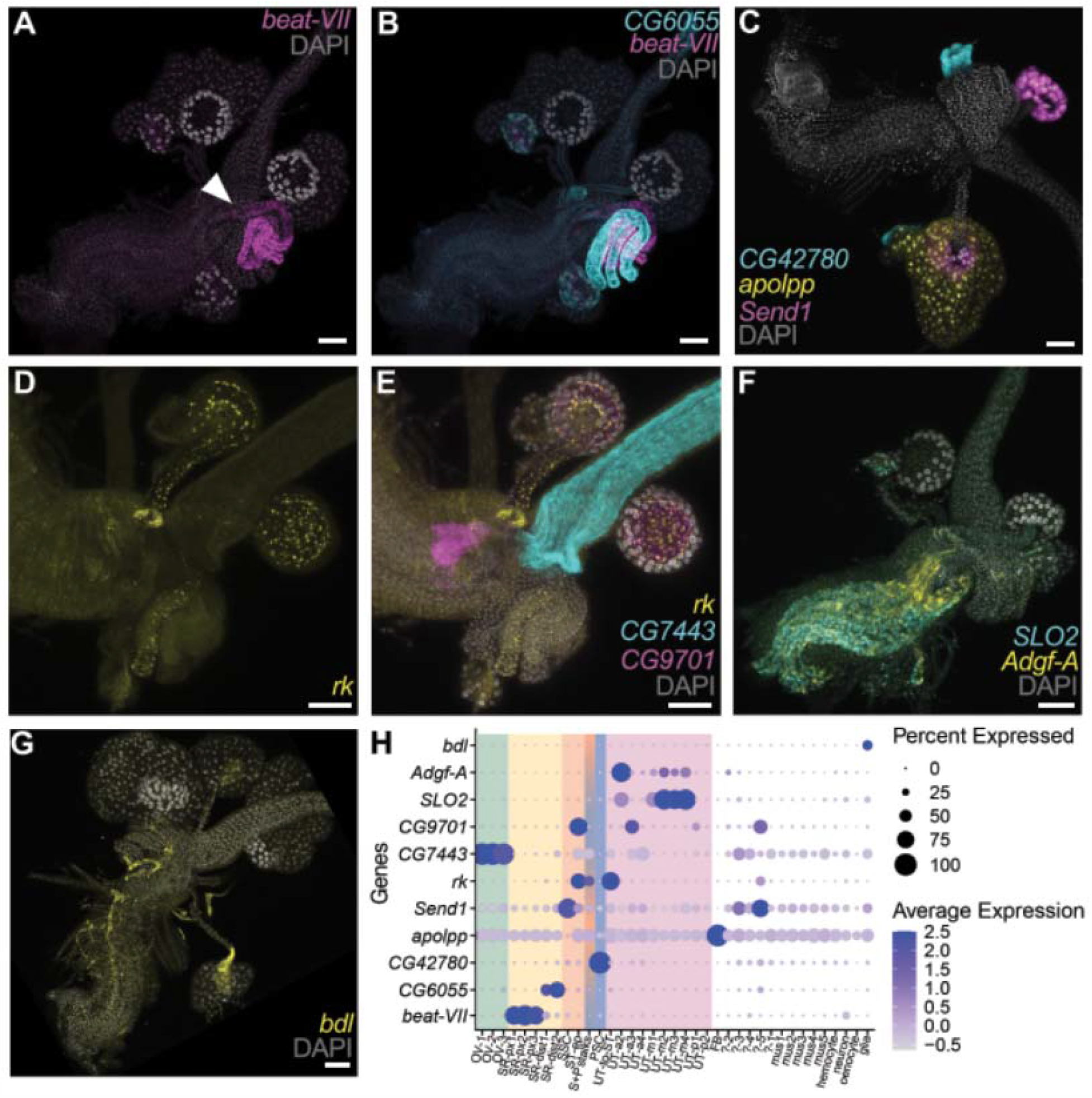
Cluster annotation with *in situ* hybridization chain reactions. *A-G)* Lower reproductive tracts from non-mated females processed for *in situ* hybridization chain reaction, with a nuclear stain (DAPI, grey). *A-B)* The proximal seminal receptacle is *beat-VII* positive (magenta), while the distal SR is *CG6055* positive (cyan). White arrowhead indicates the point at which the seminal receptacle joins the uterus. *C) CG42780* (cyan) is a marker gene for parovaria secretory cells. *Send1* expression (magenta) confirms the spermathecal secretory cell annotation. *Apolipophorin* (yellow) is a marker gene for the reproductive-associated fat body, which was dissected away from one lateral side of the tissue. *D-E)* A ring of cells in the uterine wall that encircle the entry point to the spermatheca and the stalks of the spermathecae and parovaria are all *rk* positive (yellow). *CG9701* expression (magenta) marks the oviduct valve flap. *In situ* probes against *rk* and *CG9701* transcripts co-localize in the spermathecal epithelial cells. *CG7443* (cyan) is an oviduct marker gene. *F) SLO2* (cyan) and *Adgf-A* (yellow) are co-expressed in much of the uterus, with relatively stronger *Adgf-2* expression in a subset of the anterior uterus versus higher *SLO2* expression through the middle uterus. *G) borderless* (yellow) is a marker gene for glia. *H)* Expression across clusters for the marker genes targeted with *in situ* hybridization chain reaction. Scale bar = 50 micrometers.

In addition to its core functions managing sperm and eggs, the FRT is the site of high-stakes interactions with male products, specifically seminal fluid proteins (SFPs). SFPs are proteins that males transfer to females during copulation; there are >290 in *D. melanogaster* (Sepil et al. 2019). Upon recognizing the receipt of male products, the FRT becomes the initiation site for a cascade of systemic post-mating responses: increased activity and reduced sleep (Isaac et al. 2010), altered neural gene expression (Yael Heifetz et al. 2014), reduced receptivity to remating (Chen et al. 1988), increased aggression (Bath et al. 2017), improved learning (Scheunemann et al. 2019), reduced immune capacity despite systemic upregulation of immunity genes (Fedorka et al. 2007), and even reduced lifespan (Chapman et al. 1995). Taken together, it is clear that events in the lower FRT, including signal transduction in response to mating, can have profound, systemic consequences for female biology.

Despite model organism status, only six gene expression surveys of the female *D. melanogaster* somatic reproductive tract are reported (Prokupek et al. 2009; Leader et al. 2018; Mack et al. 2006; McDonough-Goldstein, Borziak, et al. 2021; Allen and Spradling 2008; Robinson et al. 2013). Of these, four used old methods that are incomplete or biased and five of these studies either aggregated all somatic reproductive tissues or included only the spermatheca. Only one unbiased transcriptomic analysis of each lower reproductive tract tissue exists (McDonough-Goldstein, Borziak, et al. 2021). Recent single-cell transcriptome investigation of D. melanogaster by the Fly Cell Atlas consortium excluded the somatic female reproductive tract (Li et al. 2022), leaving gene expression at cellular resolution wholly unexplored in these tissues. This state of affairs stands in contrast to the level of research investment in male somatic reproductive tissue–the accessory glands–which have been characterized in every generation of the Fly Atlas project (Chintapalli, Wang, and Dow 2007; Leader et al. 2018) and in many independent transcriptomic investigations. Given the similarities between fly and mammal reproductive tracts, such as their highly secretory nature and shared enrichments for functional classes of expressed products (e.g. serine proteases, (Muytjens, Yu, and Diamandis 2018; Lawniczak and Begun 2007; Kelleher and Pennington 2009), the lack of FRT data contributes to the general research shortfall in female subjects in the biomedical literature (Beery and Zucker 2011; Woitowich, Beery, and Woodruff 2020; Orr et al. 2020). In direct consequence, marker genes for key female organs and cell types are generally unknown and functional tools such as GAL4 drivers are lacking, leaving outstanding unanswered questions about female reproduction. To address this gap in the literature, we characterized gene expression and cell type diversity in the *Drosophila* female reproductive tract, using single-nucleus RNA-sequencing and fluorescent *in situ* hybridization chain reaction.

## METHODS

### Fly strains and husbandry

We sampled unmated, 2-3 day old female flies from 30 isofemale lines that were established from gravid females collected from Fairfield, Maine (September, 2011) and Panama City, Panama (January 2012) (Zhao et al. 2015). Flies were housed on standard yeast-cornmeal-agar food at 25° C on a 12-hour light:dark cycle.

### Nuclei isolation and sequencing

Whole lower reproductive tracts were dissected in chilled Schneider’s insect medium. Five reproductive tracts per each of 15 isofemale lines (i.e. 15 genotypes; 75 animals) from Maine were pooled and processed together. The same sampling strategy was followed separately with 15 genotypes from Panama, resulting in two sequencing libraries that together included 150 flies from 30 genotypes. Nuclei were isolated using a modified protocol from (Martelotto n.d.; Majane, Cridland, and Begun 2022). Briefly, tissues were dounced in lysis buffer, passed through a sterile filter, and centrifuged. The clarified preparation was stained with 10 μg/ml of DAPI prior to FACS analysis and selection by flow cytometry technical personnel using the Beckman Coulter “Astrios” cell sorter. The Astrios was pre-configured for rapid, chilled processing of the nuclei preparations using the 70 _μ_m nozzle at 60 psi fluid pressure. The total cellular preparation was assessed for laser light scatter (forward vs. side angle scatter) and gated based on laser scatter to exclude debris and aggregates.

Putative single nuclei were assessed for DAPI intensity following illumination with a 405 nm laser and detection in a 450/50 restricted photodetector (Supp figures x). The majority of the sample (59%) exhibited little or no fluorescence (DAPI negative, median fluorescent intensity [MFI] = 39) or low fluorescence (DAPI low, 33.7%, MFI = 571). The DAPI bright positive nuclei (fluorescent intensity > 5000), which comprised ∼4.8% of the total cellular preparation, were collected for subsequent genomic analysis. A total of at least 100,000 DAPI+ nuclei were collected per sample. Barcoded 3’ single nucleus libraries were prepared with the Chromium Next GEM Single Cell 3’ kit v3.1 (10X Genomics, Pleasanton, California) according to the manufacturer recommendations. Library quality was assessed with a Bioanalyzer 2100 (Agilent, Santa Clara, CA). The libraries were sequenced on a NovaSeq 6000 sequencer (Illumina, San Diego, CA) with paired-end 150 bp reads. The sequencing generated approximately 12,671 / 13,554 reads per cell and 128 / 220 million reads per library for Maine and Panama respectively. The Maine and Panama libraries recovered 10,073 and 16,207 nuclei, respectively.

### RNAseq data preparation

Alignment, barcode removal, and UMI counting were done in CellRanger (6.1.2) with the “count” command. Reads were aligned to version 6.41 of the *D. melanogaster* genome (FlyBase, downloaded August 9, 2021), with an index built using CellRanger “mkref.” To the FlyBase *D. melanogaster* 6.41 GTF file, we added *de novo* gene annotations from (Cridland et al. 2022) and de novo and newly assembled gene annotation models from (Lombardo et al. 2023). Because the cellranger pipeline does not accept annotations with unknown strandedness, we edited the annotations for all single exon de novo genes to be on the ‘+’ strand. We then used CellRanger “mkgtf” to remove non-polyA transcripts that overlapped with protein-coding gene models.

### RNAseq data reduction

We used R 4.1.2 to curate and analyze the dataset. SoupX v1.5.2 (Young and Behjati 2020) was used with default parameters to remove cell-free mRNA contamination. Using Seurat v4.3.0 (Satija et al. 2015), we applied filters to remove genes detected in fewer than 3 nuclei, nuclei with >30,000 RNAs, nuclei with >275 genes, and nuclei with >2% mitochondrial gene expression. Because we recovered more nuclei than the 10,000 maximum that 10x Genomics recommends per sequencing lane, crowding and doublets were likely–an especially devilish problem given that little information that could be used to guide clustering was previously available (e.g. how many cell types are present; validated marker genes). To remove doublets, following preliminary clustering in Seurat, we used doubletfinder (McGinnis, Murrow, and Gartner 2019), a high-performing tool (Xi and Li 2021), according to default settings and assuming a 7-8% percent doublet rate in the ME and PAN libraries, respectively, per the 10x Genomics user guide. However, doubletfinder heavily penalized polyploid cell types, to such an extent that accepting all of doubletfinder’s designations would result in complete loss of the PSC and SSC clusters. Therefore, we did not remove putative doublets from the two clusters corresponding to the polyploid spermathecal and parovaria secretory cells. The polyploid fat cell cluster was also heavily penalized (about 18% of cells flagged by doubletfinder). Though this aggressive pruning may also include false positive doublet identifications, it was not so severe that the cluster, nor its signatures of polyploidy including high average RNA counts, were lost. To retain a conservative set of high-confidence singlets, we removed all putative doublets from the fat body cluster.

### RNAseq clustering and data analysis

Prior to clustering, singlets from the ME and PAN libraries were combined using “merge” in Seurat. Data were normalized and scaled using the following functions in Seurat: “NormalizeData”, “FindVariableFeatures” set to 12,300 features, and “ScaleData” with the argument to regress out variation in the percentage of mitochondrial reads. Nuclei were clustered using “FindNeighbors,” “FindClusters”, and “FindUMAP” with principal components 1 to 75. We used “clustree” (Zappia and Oshlack 2018) to evaluate cluster stability across resolution settings from 0.5 to 4 before proceeding with a resolution of 1.5. Cluster similarity was evaluated using the Seurat function “BuildClusterTree” and principal components 1 to 75. Genes were called as ‘expressed’ using two different thresholds. For analyses focused on determining the typical attributes of a cluster (GO enrichment analysis), we called a gene ‘expressed’ if it had >0 logCPM in >50% of cells per cluster. This strict threshold conservatively includes genes whose expression we can be confident is typical for the cluster. However we observed that, perhaps owing to the gene dropout challenges inherent to single cell sequencing, this threshold was too strict to detect some externally validated expressed genes. Antibody staining (Rezával et al. 2012) indicates that all FRT neurons express *ppk*, while *fru* and *dsx* expression can be used to identify subsets of FRT neurons. In our dataset, the neuron cluster robustly expresses both *fru* and *dsx*, but *ppk* is only detectable in 2/34 cells (just under 6% of cells). Therefore, for analyses focused on gene discovery (e.g. to report the total set of expressed genes / SFPs), we applied an empirically-calibrated threshold wherein a gene is considered ‘expressed’ if it is detected at least as strongly as was *ppk* in the neuron cluster; specifically, >1 logCPM in >6% of cells per cluster.

### HCR in situ hybridization

Lower reproductive tracts were dissected from flies with matched genotypes, ages, and mating status to the sequenced specimens. Tissues were fixed with 3.2% paraformaldehyde for 40 minutes, methanol dehydrated, and stored at -20**°**C until the hybridization reaction. Prospective gene targets were identified using the Seurat “FindAllMarkers” and “FindMarkers” functions, and prioritized using the Seurat visualization tools “VlnPlot” and “DotPlot” (Supplement) along with practical considerations–transcript length and an absence of overlapping gene annotations. Given the large number of putative cell types to be annotated in a thick, whole-mount tissue with melanized and chitinous regions prone to autofluorescence, we used a high-throughput approach with multiplexed probes and a split-initiator scheme that minimizes background fluorescence (Choi et al. 2018). Probes were designed using previously published software (Kuehn et al. 2022) implemented with (https://github.com/rwnull/HCRProbeMakerCL) with the following settings: 0 basepair 5’ offset, 5 maximum A/T homopolymer length, 4 maximum C/G homopolymer length, between 35-55% CG bases in probe, BLAST against v 6.41 of the D. melanogaster reference genome, and a maximum of 40 probe pairs. For short genes that otherwise would not accommodate enough total probe pairs (*CG6055, CG7443, CG17108, CG17239*, and *red*) we relaxed the limits to 31-61% CG probe content. We used transcript sequences from the Drosophila genome v. 6.41 and substituted in SNP alleles that occur at greater than 50% frequencies in the east coast source populations per (Svetec et al. 2016; Reinhardt et al. 2014). Probes were preferentially designed against only the CDS unless CDS length was too short. Probes were ordered from Integrated DNA Technologies, Inc., (Coralville, Iowa, USA) in three-gene oligo pools, where the probe sequences for three genes were designed to each use different initiation sequences and were ordered pooled in a single tube. Probe sequences are listed in (Supp Table 1). Successful probe sets had from 8-40 probe pairs per gene. We used three amplifier hairpin designs from Molecular Instruments Inc., (Los Angeles, CA, USA), specifically the B1 initiation sequence conjugated to AlexaFluor 647, B2= AlexaFluor488, and B4 =AlexaFluor 546. Amplifier hairpin and initiation sequences were developed by (Choi, Beck, and Pierce 2014). Hybridization reactions followed the (Bruce et al. 2021) protocol, using 3x probe concentrations, no sonication step, a 200 uL reaction volume for hairpin amplification, and 20+ hour incubation times for the hybridization and amplification steps. For small genes with ten or fewer probe pairs (*CG7443, CG43112, EbpIII*), we doubled the concentration of the probe within the oligo pool during probe design, resulting in an effective 6x probe concentration. In other words, we optimized the protocol for bright signal in a high-level whole-tissue image, to be able to detect rare or disperse cell types for which we had no a priori information regarding which region to focus on while imaging. Negative controls received the same treatment, including the application of hairpins, except that no probes were applied. We also used a second negative control strategy, using previously validated probe sets against two genes, *wg* and *snail*, which were not expressed in our single nuclei dataset (Supplement, ‘Pool B’ section).

### Confocal microscopy and image processing

Samples were mounted in 50% glycerol / PBS and imaged on a Zeiss Airyscan 980 in confocal mode, using the 355, 488, 561, and 639 nm lasers. Consistent with our objective to locate cell types, rather than to precisely quantify signal, we optimized laser power and gain settings to detect signal for each sample, allowing variation in excitation between gene sets and replicates. Consequently, relative brightness may be compared within but not among images in our dataset. We imaged negative controls with both maximal and typical (mean and median) settings for laser power and gain (see full presentation of negative controls in Supplement). Images were processed in ImageJ v2.3.0 as follows: for each laser channel, Z-series were projected using the maximum intensity setting, and the ‘Maximum’ slider in the ‘Brightness & Contrast’ settings was adjusted to set the brightest pixels at saturation, while viewing the image with the ‘HiLo’ lookup table. As with gain and laser power, brightness was optimized separately for each sample and color channel, with maximal and typical adjustments applied to the respective negative controls. All brightness adjustments were uniformly applied to the entire image. Color channels were imported as layers to Adobe Photoshop 2015.5.2 and merged using the ‘screen’ overlay setting.

## RESULTS

### Nuclei clustering and annotation identify novel cell types

To characterize gene expression among cell types in the Drosophila female somatic reproductive tract (FRT, Fig. 1B), we generated two single-nucleus sequencing libraries from unmated females. To ensure that our results, including putative cell type designations with their marker genes and transcriptomic profiles, would be robust to potential genotypic variation, we sampled from 30 wild-derived isofemale line genotypes from the North American east coast, evenly represented in these libraries (Methods). Sequencing generated approximately 128/220 million reads per library, with an average of 12,671/13,554 reads per nucleus among the 10,073 /16,207 recovered nuclei per library. To discover female reproductive tract cell types we clustered nuclei by gene expression principal components of variation, resulting in 37 putative cell types (Fig. 1C-D).

We then set out to first determine which of our inferred clusters were likely to represent previously described cell types in the species. Using the known marker *Send 1* for spermathecal secretory cells (SSC, Fig. 2C (Schnakenberg, Matias, and Siegal 2011) and known markers *fru, dsx*, and *ppk* for neurons (Rezával et al. 2012) we assigned clusters to these cell identities. We then investigated which of our clusters might correspond to cell types previously identified in Fly Cell Atlas (Li et al. 2022). To do so, we extracted marker genes for each cluster using Seurat’s “findallmarkers” function and then investigated their expression in Fly Cell Atlas, followed by reciprocally checking top marker genes of potentially matching Fly Cell Atlas clusters in our dataset. The following matched sets of marker genes and cell types in Fly Cell Atlas allowed us to annotate multiple cell-type clusters in our dataset: oenocytes, *FASN2, FASN3, LpR1*; hemocytes, *Hml, Ppn, Nimc1*; muscle, *bt, sls, up, Octalpha2R, CG44422, CG45076*; fat body, *CG13315, CG4716, Ubx, apolpp*; sensory neuron annotation, *para, Rdl*, and *futsch*. Another cluster, “?-1” (Fig. 1C), shared some marker genes (*CG8012, CG5162, wat*) with adult tracheal cells, but its top two marker genes (*Antp–*see Supplement, and *CG15353*) were instead only expressed in an unannotated cell type in the Fly Cell Atlas tracheal dissection dataset. Within Fly Cell Atlas, this latter, unannotated cell type was distinguished from adult tracheal cells by the expression of *wgn, pk*, and *gol*, which are unexpressed in Fly Cell Atlas tracheal cells and in our cluster. While some type of tracheal identity is most likely for this cluster, we have taken the conservative approach of leaving it unannotated in our figures. Finally, we determined whether for the unidentified clusters, our top 4 marker genes corresponded to gene expression inferred from individual organ dissection of the FRT (McDonough-Goldstein, Borziak, et al. 2021). These comparisons allowed us to identify 3 uterine clusters, 1 seminal receptacle cluster, 1 parovaria cluster, and 3 oviduct clusters. Additionally, cross-checking confirmed that our top spermathecal secretory cell marker genes were most highly expressed in the ST bulk tissue transcriptome.

For the remaining clusters, our marker genes did not all exhibit their highest expression in bulk organ-level FRT transcriptomes (McDonough-Goldstein, Borziak, et al. 2021). Thus, in total, based on prior literature we identified 5 cell types, with suggestive or tissue-level annotations for an additional 14 clusters (including “?-1”). The remaining 18 clusters are unlikely to correspond to cell types with known molecular identities in this model species.

To further investigate the unknown clusters, we performed fluorescent *in situ* hybridization chain reaction using subsets of cluster marker genes (Fig. 2, Supp). By targeting combinations of marker genes, we annotated 13 additional clusters (Fig. 1 C-D) and discovered extensive novel cell type heterogeneity within tissues and striking spatial regionalization. For example, previous research described two morphologically distinct regions in the seminal receptacle: a narrow proximal section and a contrastingly thicker distal section, which also differ in the arrangement of microvilli on the inner surface, the density of secretory vacuoles, and the postures adopted by occupying sperm within each section (Y. Heifetz and Rivlin 2010). We find molecular markers that are specific to the seminal receptacle and distinguish these two sections: the proximal SR is *beat-VII* positive, while the distal section is marked by *CG6055* (Fig. 2A-B). We also find molecular markers and cell types that map onto spatially discrete and thus, likely functionally specialized regions in the uterus, including a previously undescribed region in the uterine wall encircling the entrance to the spermathecal ducts, marked by *cyp313a3* and *rk* (UT-into-ST, Fig. 2D-E and Supplement, pool H section). *CG9701* marks the cells of the oviduct valve flap (Adams and Wolfner 2007), a portion of the inner, anterior uterine wall that changes conformation to allow access to the oviduct and storage organs (UT-a3, Fig. 2E). *obst-E* marks the UT-OVF + UT-into-ST clusters and adds a third expression domain that corresponds to UT-a4 cells, which is located along the portion of the anterior uterine wall that becomes the papillate elevation region after mating (Supplement, pool H section, (Adams and Wolfner 2007).

Next, we sought to use our cell-level data to significantly augment organ-level expression data from the FRT (McDonough-Goldstein, Borziak, et al. 2021) (Leader et al. 2018)) and distinguish between transcripts produced in reproductive cells from those produced by connective tissues and other unintentional ‘by-catch’ in bulk tissue transcriptome sequencing. For example, (McDonough-Goldstein, Borziak, et al. 2021) reported a significant enrichment of expressed receptor genes unique to the seminal receptacle, with 10 SR receptor genes detected – an exciting prospect, since females are thought to respond to the receipt of seminal fluid proteins, yet only one SFP receptor has been identified (*Sex peptide receptor*, (Yapici et al. 2008)). In our data, five of these putative SR receptor genes are unexpressed in any SR cluster, and 4 are instead expressed by hemocytes. Moreover, our top hemocyte marker genes are most strongly expressed in SR transcriptomes from the (McDonough-Goldstein, Borziak, et al. 2021) bulk-tissue analysis. Together, these observations suggest that hemocytes may be disproportionately captured alongside the SR in bulk tissue dissections, and that the observed enrichment for receptor genes in the bulk-tissue SR transcriptome may be artefactual. To identify a high-confidence set of female reproductive tract genes we report ∼5960 genes expressed in our 22 validated reproductive (e.g. non-muscle, -hemocyte, -oenocyte, -glia) clusters (Supplement Table 2). Additionally, our findings greatly expand the number of female reproductive tract marker genes (Supplement Table 3), which can facilitate future resource development and functional analyses. This list includes a much needed, perfectly specific marker gene for parovaria secretory cells (*CG42780*, Fig. 2C,H), as well as marker genes for spermathecal epithelial cells, the glandular ducts, and a variety of morphologically specialized regions throughout the seminal receptacle and uterus. Indeed, many FRT marker genes identified in our analysis are not currently associated with any functional annotation, demonstrating the great potential value of these data.

### Polyploidy is pervasive

While the spermathecal secretory cells have previously been recognized as polyploid (Mayhew and Merritt 2013; Almeida Machado Costa et al. 2022), we find that polyploidy is common in the reproductive tract, occurring in at least 5 of the 6 major reproductive tissues, and spanning four different ploidy levels. In aggregate, our data suggest that more than half of the FRT cells are polyploid. Four lines of evidence support this conclusion. First, we noticed disparate nuclei sizes upon visual inspection of dissociated nuclei; this was also noticeable in confocal micrographs stained with DAPI. Second, quantitative evidence of polyploidy was observed while FACS sorting the nuclear preparation prior to sequencing, evidenced by discrete strata of DAPI fluorescent intensity (Fig. 3A). DAPI bright positive nuclei exhibited at least four levels of increasing intensity, corresponding to 4 quantized levels of DNA abundance per nucleus, with ∼29% of the nuclei exhibiting a median fluorescent intensity (MFI) of 7,061, 20% with MFI of 11,548, 25% with MFI of 19,754 and a lesser population of 4% of total nuclei at an MFI of 40,112 intensity units (Supplement). Notably, median fluorescence intensity approximately doubles with each step. Further evidence comes from highly variable total RNA counts per cell type (Fig. 3B), where SSC, PSC, and fat cells in particular stand out as having very high total RNA counts per cell; certain oviduct and uterine cells are also elevated. Finally, based on expressed genes per cell type, 14 clusters had significant Gene Ontology term enrichments for ‘polytene chromosome’, ‘polytene puff’, and/or ‘polytene band’ (Fig. 3B), a chromosomal morphology associated with polyploidy in Drosophila. To further investigate this we checked the expression of genes that regulate or initiate endoreplication (Fig. 3C, (Almeida Machado Costa et al. 2022). These genes are associated with in-progress endoreplication and are not necessarily expected to mark cells that previously completed a limited round of endoreplication. When endoreplication is underway, overexpression of *Myc* can stimulate additional rounds of endocycling. Several signaling pathways can regulate endocycle in specific contexts (Almeida Machado Costa et al. 2022). Patterns of gene expression in our data suggest that the JNK pathway (terminal kinase *Bsk*), the Hippo pathway (terminal member *Yki*), and the MAPK pathway (activated by *KDM5*) may influence endocycling in the FRT. One recognized function of polyploidy is to achieve a higher transcriptional output, especially in secretory cells; in this context, the extensive polyploidization of the FRT is consistent with the volume of secretions that it produces.

**Figure 3:**
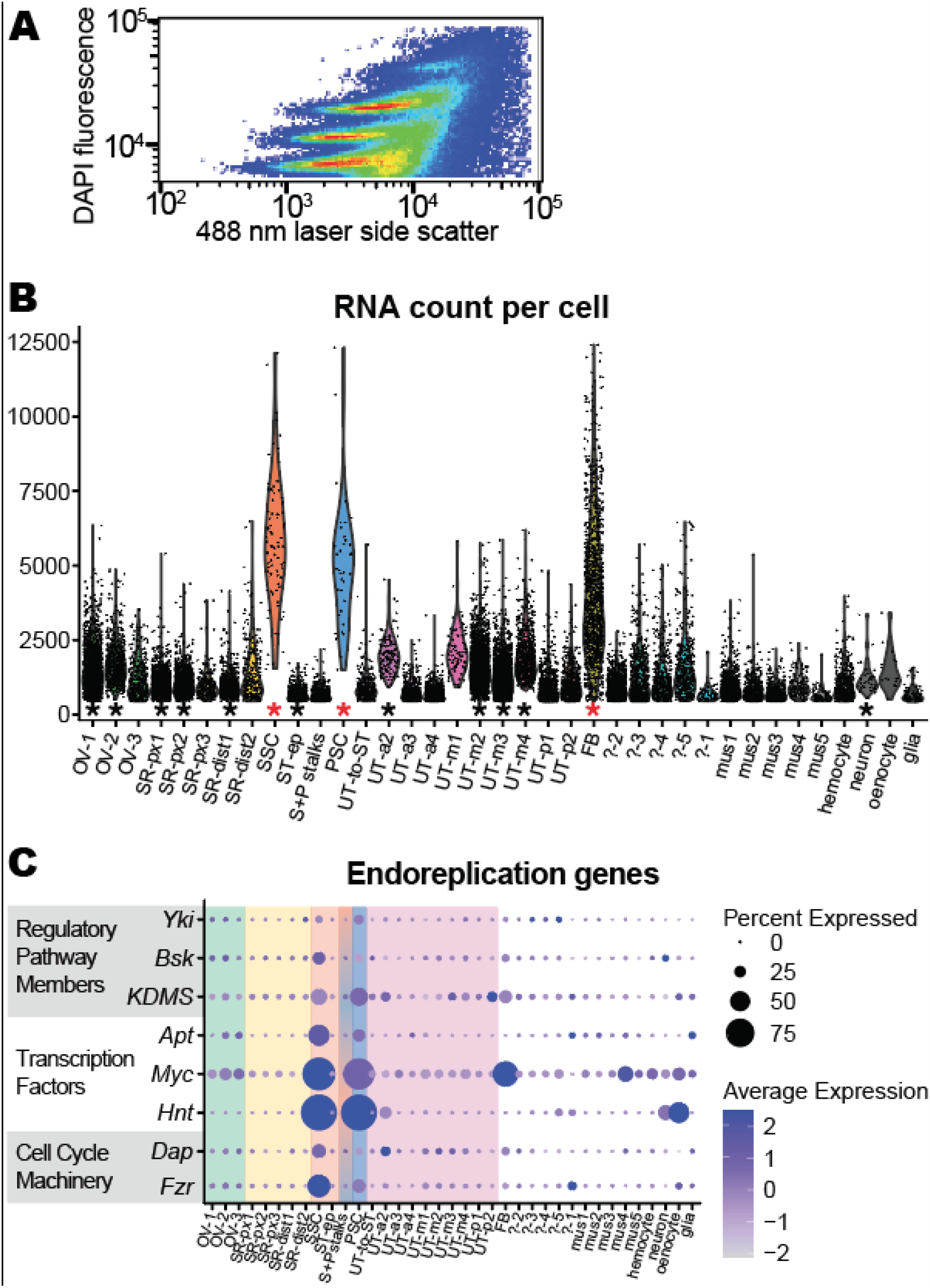
Polyploidy is common in the FRT. *A)* Nuclei segregate into strata during FACs sorting by side scatter (size) and DAPI fluorescent intensity, with nuclei frequency shown by heat map colors. *B)* RNA count per cell varies dramatically among clusters. Asterisks above the cluster ID along the x-axis indicate significant Gene Ontology enrichments for “polytene chromosomes,” a chromosomal morphology associated with polyploidy in Drosophila (red = adjusted p < 10^-4^, black = adjusted p < 0.01). *C)* Expression of genes associated with the endoreplication process (Almeida Machado Costa et al. 2022).

### Female reproductive cell types specialize in producing seminal fluid proteins

While seminal fluid protein gene (SFP) functions are typically interpreted with respect to their production and transfer by males and their influence on females, we find many SFP genes expressed in unmated female reproductive tracts. Importantly, certain FRT cell types are enriched for SFP transcription (Fig. 4C). For example, in cluster “SR-dist2”, a cell type in the distal seminal receptacle, 30.4% of mRNAs in the median nucleus are from SFP genes, and SFP transcripts comprise up to 65% of mRNAs in some nuclei. This result is driven by transcripts of 11 very highly expressed SFP genes, 22 with moderate expression, and an additional 15 detected (total 48 expressed SFP genes). Cluster “?-5” also appears specialized for SFP transcription, with a median of 11% of mRNAs aligning to SFP genes; this cluster expresses the greatest number of SFP genes relative to other FRT clusters (60 genes; 20% of the 292 *D. melanogaster* SFPs). Other clusters for which >5% of mRNAs align to SFPs in the median cell are the spermathecal secretory cells (9%), the ST epithelial cells (5%) and the remaining SR clusters (SR-prox 1-3 and -dist1, 5-8%). In total, 78 of the 292 *D. melanogaster* SFPs (>26% of SFPs) are expressed in one or more FRT cell types. SFPs are enriched in different cell types than other biologically relevant gene sets (Fig. 4), and their expression is not a simple correlate of high overall levels of transcription (Fig. 3B), mating responsivity (Fig. 4 A-B), nor secretion (Fig. 4 D).

**Figure 4:**
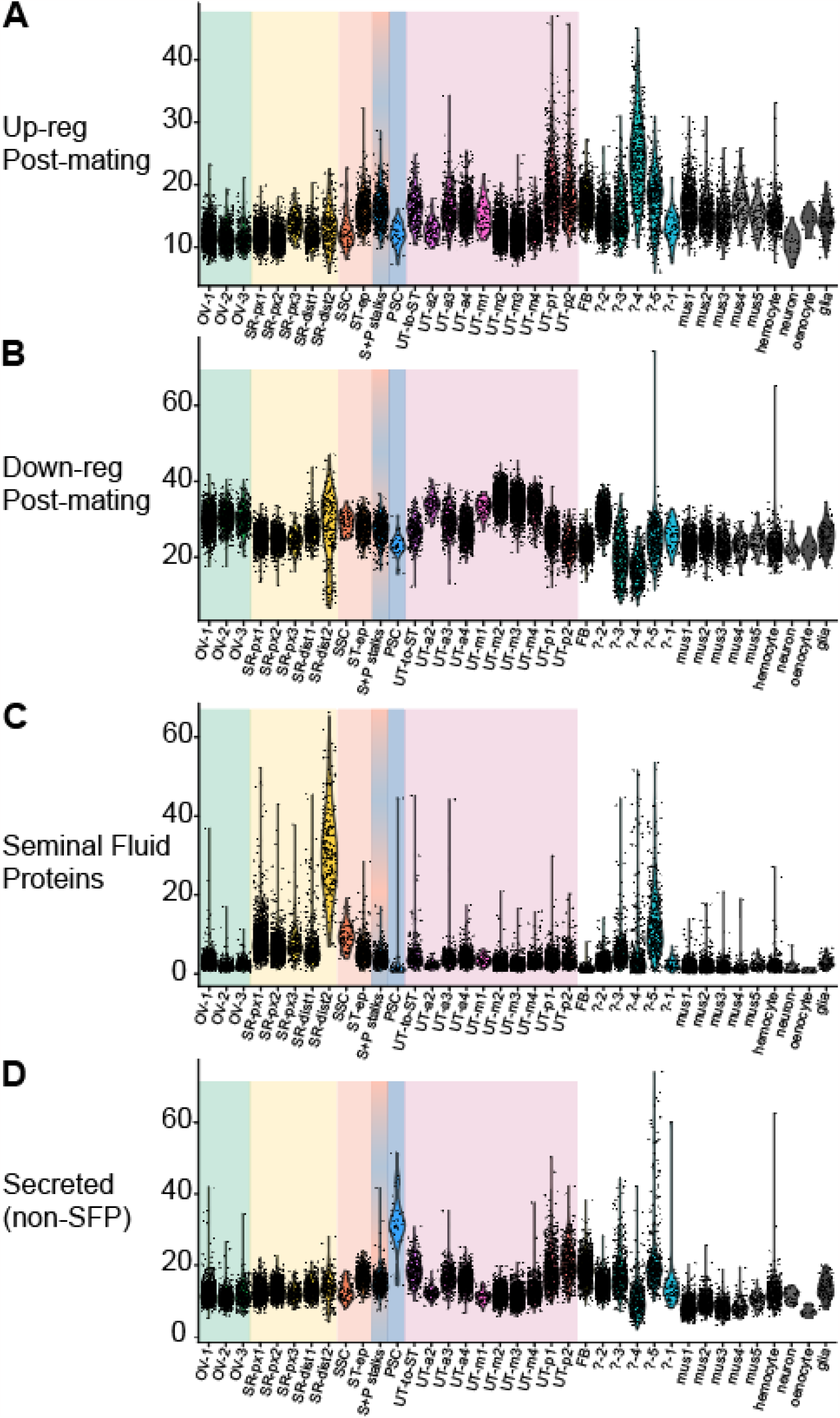
The expression of biologically relevant gene sets varies among cell types in the FRT. *A-D)* Each point is one cell; each violin is the distribution of cell values for one cell type cluster. The y-axis gives the percent of RNA read counts per cell that align to genes belonging to the relevant functional category. Background colors correspond to organ-level identity of the clusters, as given in Fig. 1B. *A)* Expression of genes that are up-regulated in response to mating in (McDonough-Goldstein, Borziak, et al. 2021). *B)* Expression of genes that are down-regulated in response to mating. *C)* Expression of genes which produce seminal fluid proteins (SFPs, as identified in (Wigby et al. 2020). The seminal receptacle, in particular, vigorously transcribes seminal fluid protein genes. *D)* Transcription of genes that have a secretion signal peptide detected by the SignalP algorithm (Almagro Armenteros et al. 2019), excluding seminal fluid genes.

To further investigate the prevalence of FRT-expressed SFPs, we looked at previously published FRT transcriptomes (McDonough-Goldstein, Borziak, et al. 2021; Cridland and Begun 2023). These two studies complement our work by supplying gene expression profiles for the mated-state FRT (McDonough-Goldstein, Borziak, et al. 2021) and by the use of contrasting female and male genotypes to distinguish male-contributed RNAs from endogenous female RNAs in the FRT after mating (Cridland and Begun 2023). Taking evidence from these studies together with our results, at least 127 of the 292 known SFP genes–43% of SFPs–are expressed by the female reproductive tract. Intriguingly, this set of female-expressed SFPs includes several individually notable SFPs, specifically, SFPs involved in sperm storage (*est-6, Acp62F, Pde1c*); in binding Sex Peptide to sperm and, ergo, the maintenance of the long-term post-mating response (*lectin-46Ca, lectin-46Cb*); in influencing female remating receptivity (*CG10433*, and nonsignificant trend *CG32833*); and *Dup99B*, which is a predicted duplicate of *Sex Peptide* that can weakly elicit aspects of the systemic post-mating response (Saudan et al. 2002). If we further include suggestive evidence of gene expression (i.e. detection in female carcass (McDonough-Goldstein, Borziak, et al. 2021); detection in mated female FRTs in study designs that cannot exclude the possibility the RNAs were transferred by males), there may be >160 SFPs produced by females, i.e., the majority of *melanogaster* SFPs. The functions of most SFPs are unknown; these data suggest that investigating their function in females is necessary to attain a realistic view of their biological roles.

## DISCUSSION

The work presented here has several major implications for our understanding of female insect reproduction. We provide transcriptomes, functional gene set enrichments, and marker genes for 22 reproductive cell types, all but one of which (SSC) was previously only morphologically defined (ST cap epithelial cells, PSC, and morphologically described cell types within SR and UT) or entirely unknown (cell type surrounding the entry to the ST, “UT-into-ST”). Many of these new marker genes are unnamed genes for which no functional information was previously available, so their status as marker genes in the FRT adds substantial biological insight.

SFP genes have canonically been primarily thought of as coding for male-expressed proteins. Indeed, recent analyses have established criteria for SFP identification that include evidence of transfer from males to females during copulation (Wigby et al. 2020). Whereas several SFPs play a role in initiating female post-mating responses (Liu et al. 2014; Isaac et al. 2010; Chen et al. 1988; Chapman et al. 1995; Pilpel et al. 2008; Avila and Wolfner 2009; Rubinstein and Wolfner 2013), the past 30 years of literature have often explored the rhetorical framing that males use SFPs to control, manipulate, or harm females (Chapman et al. 1995; Rice 1996; Wigby and Chapman 2004; Hollis et al. 2019; Gioti et al. 2012). From this perspective, SFPs are an important substrate of interlocus sexual conflict against which females must continually evolve defensive countermeasures, which has been hypothesized to explain observations that SFPs evolve rapidly (Haerty et al. 2007; Patlar et al. 2021). Our finding that females transcribe >40% of all SFP genes, and that SFPs are enriched in specialized female cell types, suggests this simple view is incorrect. For perspective, we can tally the SFPs asserted to influence post-mating female biology (Wigby et al. 2020). We liberally include non-significant trending results (*CG32833*, (Sirot et al. 2014)); suggestive evidence from an association study (*msopa*, (Fiumera, Dumont, and Clark 2007)) that was not supported by gene knockdown (Patlar and Civetta 2022); conclusions based on whole-animal manipulations for which results could conceivably be mediated by male expression in non-reproductive tissues (*CG10433*, (Liu et al. 2014)); genes whose documented effect opposes male interests (i.e. accelerating female remating receptivity, *est-6 (Gilbert, Richmond, and Sheehan 1981)*; and genes that we now recognize are also expressed by the FRT (*CG10433, est-6, Acp62F, Dup99B, Pde1c, lectin-46Ca, lectin-46Cb, CG32833*), to reach a final tally of only 24 genes (8% of SFPs). Importantly, these constitute a far smaller sample of SFPs than the female-expressed subset. Thus, the premise that the majority of SFPs have strictly male functions directed at manipulating the female post-mating response in favor of male interests, should be re-evaluated (e.g. (Hopkins and Perry 2022). As for how typical it is that SFPs function in facilitating post-mating responses, >26% are found expressed in females that have never mated, but it remains possible that the proteins are secreted, trafficked to their proper destination, or activated only following mating, consistent with (Sanchez-Lopez et al. 2022). Investigating SFP function in females is necessary to attain a realistic view of their biological roles, and expression patterns in females furnish helpful clues–it is striking that SFPs are enriched in both sperm storage organs, consistent with possible roles in moving, activating, or nourishing sperm.

While it is well appreciated that lower reproductive tract functions include interacting with male products, the FRT’s potential interactions with eggs, beyond physically moving them from ovulation through oviposition, are poorly understood. In many animals, eggs are still under construction post-ovulation, with major contributions to egg development occurring in the somatic reproductive tract. For example, in birds, biomineralization of the eggshell is accomplished by the uterus (Gautron et al. 2021). In many insects, the parovaria (i.e. female accessory glands) produce additional substances applied to the egg’s exterior, such as adhesives or silk to anchor the egg after oviposition; or, particularly among aquatic insects, gelatinous substances that hold a clutch of eggs together (Lancaster and Downes 2013). These additional coatings can also enhance desiccation resistance and deter predation or parasitism by their sticky or tough textures. In *Drosophila*, the eggshell is produced pre-ovulation by follicle cells, including the deposition of three major layers and enzymatic treatments that covalently crosslink the eggshell proteins. Prior work has identified proteins that constitute the eggshell via mass spectrometry and gel electrophoresis (Fakhouri et al. 2006; Waring and Mahowald 1979). However, owing to technical limitations on solubilizing or separating proteins after cross-linking, these studies focused on pre-ovulation eggs, meaning that if subsequent contributions or modifications occur post-ovulation in the lower reproductive tract, those effects would go undetected. The FRT expresses more than 1000 genes with secretion signal peptides, resulting in a secretion-rich environment in its lumen, including many enzymes, antifungal proteins, and proteins of unknown function (McDonough-Goldstein, Whittington, et al. 2021; McDonough-Goldstein, Pitnick, and Dorus 2022). It would be surprising if the egg were wholly unaffected by being washed through this protein milieu. Certain FRT cell types that may be especially relevant to post-ovulation egg development, specifically the parovaria secretory cells and the two distinct cell types in the posterior uterus, show enriched transcription of non-SFP secreted genes (Fig. 4D). One can easily imagine the posterior uterine cells could apply a final protective coating or lubricant during oviposition. Interactions between FRT secretions and the egg are a promising direction for future work.

## ACKNOWLEDGEMENTS

For FACS implementation, we thank Bridget McLaughlin and Jonathan Van of the UC Davis Flow Cytometry Shared Resource Laboratory, which is supported by a UC Davis Comprehensive Cancer Center Support Grant (NCI P30CA093373). Library preparation and sequencing was carried out by Hong Qiu of the DNA Technologies and Expression Analysis Core at the UC Davis Genome Center, supported by NIH Shared Instrumentation Grant 1S10OD010786-01. Confocal images were acquired at the MCB Light Microscopy Imaging Facility at UC Davis, which is supported by NIH grant 1S10OD026702-01, with guidance from Nipam Patel and Thomas Wilkop. We thank Nipam Patel for sharing pre-stained embryonic samples, used in optimizing the imaging protocol, and Alondra Sandoval for contributing to fly dissection. This work was supported by the National Institutes of Health via grant number R35 GM134930 to D.J.B. and a Ruth Kirschstein fellowship (5F32GM146419-02) to R.C.T.

## DECLARATION OF INTERESTS

The authors declare no competing interests.

## Notes

### Competing Interest Statement

The authors have declared no competing interest.

