## Supplemental Table 1 for "Regional specialization, polyploidy, and seminal fluid transcripts in the Drosophila female reproductive tract"

### In situ probe sequences
#### Adgf-A

| Pair | Initiator | Spacer | Hybridzation | Hybridzation | Spacer | Initiator |
|---|---|---|---|---|---|---|
| 1 | CCTCAACCTACCTCCAAC | AA | ACATCGGCGATGAACTGGTCCCATT | GATCCCCGGTTGGAGGAACGATCAT | AT | TCTCACCATATTCGCTTC |
| 2 | CCTCAACCTACCTCCAAC | AA | GCGTCTCCCGTCAGAGAACTATATT | TGCCACTTTTCCAGCGCCTCGAACT | AT | TCTCACCATATTCGCTTC |
| 3 | CCTCAACCTACCTCCAAC | AA | GCAAGTCGGAGTGCTGCGAGGCGAT | AATTTAGGGCCAACTTCTTCAGCAG | AT | TCTCACCATATTCGCTTC |
| 4 | CCTCAACCTACCTCCAAC | AA | CAGGGGAGTGGCTTTCCAGAAGGAG | TAAGAAGGCGATATAGAAGTCATGC | AT | TCTCACCATATTCGCTTC |
| 5 | CCTCAACCTACCTCCAAC | AA | CCATCGGCGAAGAAGTGAGAGCACG | TCATCCGAGGATATAACCACTGGAT | AT | TCTCACCATATTCGCTTC |
| 6 | CCTCAACCTACCTCCAAC | AA | GCACTTGATTCGAAATCGGATTCAC | GGTTGCGGAAGTCAGAGACCAGCTG | AT | TCTCACCATATTCGCTTC |
| 7 | CCTCAACCTACCTCCAAC | AA | GTCCAGCACCACGGGATGTTTGACC | AATGGCCACATTCAGCTTTTTCAGC | AT | TCTCACCATATTCGCTTC |
| 8 | CCTCAACCTACCTCCAAC | AA | CCCAGCAAAATGGCATCGATCAGGT | CCGAATCCATGACCAATACGCTTTG | AT | TCTCACCATATTCGCTTC |
| 9 | CCTCAACCTACCTCCAAC | AA | TTTCTCCGGCGTGGAAGTAGAAATC | CATCCACAGTGGAACCGAACCAGTT | AT | TCTCACCATATTCGCTTC |
| 10 | CCTCAACCTACCTCCAAC | AA | GACAAAGTCCCTCAAAGGACGTCCC | GTCATCGGGGATAGATAATAATTCG | AT | TCTCACCATATTCGCTTC |
| 11 | CCTCAACCTACCTCCAAC | AA | GCTACAAATTCCGGGTACTTTTCCT | TCCTCCTGGCCCACCAAATCGAAAC | AT | TCTCACCATATTCGCTTC |
| 12 | CCTCAACCTACCTCCAAC | AA | TTACGCCCTCTGCATTCGTATAACG | TTTGTTTAAGGGTTTGGATATAGCC | AT | TCTCACCATATTCGCTTC |
| 13 | CCTCAACCTACCTCCAAC | AA | GAAATCTGGATGAGCCTCCTTGAAC | AGGAGCATAGATCATCCTTGAACCA | AT | TCTCACCATATTCGCTTC |
| 14 | CCTCAACCTACCTCCAAC | AA | GTATCCATAGGAGTAAACTCCGTTC | TCCAAGGTTTCCACATAAATGCGCA | AT | TCTCACCATATTCGCTTC |
| 15 | CCTCAACCTACCTCCAAC | AA | ATCGGAATTCCAAGTATAGAACACC | CCATGTCATATAGCGTGGGAACAAC | AT | TCTCACCATATTCGCTTC |
| 16 | CCTCAACCTACCTCCAAC | AA | AGATTGAAAATGGACATGAACGTGG | GGGGCGTACATCACCAGACCGTCGA | AT | TCTCACCATATTCGCTTC |
| 17 | CCTCAACCTACCTCCAAC | AA | TGGTTGGATACAGTGTGAGTCTCTC | AGGCGGCATTGTTGTCCTCAAACTT | AT | TCTCACCATATTCGCTTC |
| 18 | CCTCAACCTACCTCCAAC | AA | GTATTTGGCTCGCACATCGGAGAGC | CAGATAGTCATCGACCTTGTCAGCT | AT | TCTCACCATATTCGCTTC |
| 19 | CCTCAACCTACCTCCAAC | AA | GGCTTATCCTTGGAAAACCTTAGGG | CTCCAATCGCAGTCACTGAGGGCCT | AT | TCTCACCATATTCGCTTC |
| 20 | CCTCAACCTACCTCCAAC | AA | CAGAGATGCTCATAGTAGGTCAGGT | CTAAGATCTCCGTCCTGCTGGCAGG | AT | TCTCACCATATTCGCTTC |
| 21 | CCTCAACCTACCTCCAAC | AA | TTGGGCATCTTTTTCAGCAGCTTGA | GTATCGTGGGCATGAAGAACCGCGC | AT | TCTCACCATATTCGCTTC |
| 22 | CCTCAACCTACCTCCAAC | AA | CATCAAATATGTGCTGTGAGGGCTT | GATCGGTGTTCCGGATGCCGTCCAA | AT | TCTCACCATATTCGCTTC |
| 23 | CCTCAACCTACCTCCAAC | AA | AAACTCCTTTAATTTGGCCTTCATA | GAGGTGAGGGGTAACCAGGCCTTCG | AT | TCTCACCATATTCGCTTC |
| 24 | CCTCAACCTACCTCCAAC | AA | TTTAGATCCAGGTCATGGCCCAATG | GTTTCGTTAGCCTTAATTTCTCTGC | AT | TCTCACCATATTCGCTTC |
| 25 | CCTCAACCTACCTCCAAC | AA | TTCTCAACGTCTTGTAGGTATCCGG | GTGATTCCTCGTACCGGAAAAAGGC | AT | TCTCACCATATTCGCTTC |
| 26 | CCTCAACCTACCTCCAAC | AA | AACAGCTTTCTATCAACCTGGTTCT | ACGTGGGGCGTGCTGCCGTAGATCG | AT | TCTCACCATATTCGCTTC |
| 27 | CCTCAACCTACCTCCAAC | AA | CCCAAAGTGAGACAGGCGATAAGAT | GGTCCGAAGCTAAGACAGACGCACA | AT | TCTCACCATATTCGCTTC |
| 28 | CCTCAACCTACCTCCAAC | AA | ATTTCGTTAGGGTCCGATCGCGACT | GATGACTGGCGACATGATGAGCGGT | AT | TCTCACCATATTCGCTTC |
| 29 | CCTCAACCTACCTCCAAC | AA | GAGGAGCTCCGATTTTGGTTTCGAT | ACTCTGCGACTGACGTCTGCGGCGA | AT | TCTCACCATATTCGCTTC |

### apolpp (B1 initiator, pool A)

| Pair | Initiator | Spacer | Hybridzation | Hybridzation | Spacer | Initiator |
|---|---|---|---|---|---|---|
| 1 | GAGGAGGGCAGCAAACGG | AA | GATAAAACATGTAAGCCCAGTTTAG | AAAAGATTTTTCCTCACGAAGGGAT | TA | GAAGAGTCTTCCTTTACG |
| 2 | GAGGAGGGCAGCAAACGG | AA | CAAACACCCAGCCACCGTTATTAAG | CTTTTAATTTTTCAAAGTTTTGTGT | TA | GAAGAGTCTTCCTTTACG |
| 3 | GAGGAGGGCAGCAAACGG | AA | TCGCACACCGATAATGCTTTTTGCC | CCAATTCTTATATTCTAAAGAATCA | TA | GAAGAGTCTTCCTTTACG |
| 4 | GAGGAGGGCAGCAAACGG | AA | AAAACAATCACTCCGACTTGTACAT | AGAGCGGGATATCGTTTGGTTTCCT | TA | GAAGAGTCTTCCTTTACG |
| 5 | GAGGAGGGCAGCAAACGG | AA | GTAATAGCACCCACTTATTAATCTG | GTCCTGCGCACTTAATGCAACGCGG | TA | GAAGAGTCTTCCTTTACG |
| 6 | GAGGAGGGCAGCAAACGG | AA | TTATTGTCAAATTCAATTGCTGTGC | TCCTCTCTTTTGGAGCTTTTAAACT | TA | GAAGAGTCTTCCTTTACG |
| 7 | GAGGAGGGCAGCAAACGG | AA | ATATAAGTGAATAGTGTAAAATCGG | ACATACTATGCCTACTCCGTACTCA | TA | GAAGAGTCTTCCTTTACG |
| 8 | GAGGAGGGCAGCAAACGG | AA | CGCGAAATTCAGGCCGTCGAAAGTG | GAGAATATATTTGCAGTTGCCCGGA | TA | GAAGAGTCTTCCTTTACG |
| 9 | GAGGAGGGCAGCAAACGG | AA | AAATGGAACGAACAGCCTTTATTAA | CTGGATAAGTGCTTCTAATCGAGAC | TA | GAAGAGTCTTCCTTTACG |
| 10 | GAGGAGGGCAGCAAACGG | AA | TTACAATTTCTCCAATAGCTTTTCC | CTTCCTGAGCAGCGTCGTTTATAGG | TA | GAAGAGTCTTCCTTTACG |
| 11 | GAGGAGGGCAGCAAACGG | AA | ATAATCGGTGCAATTGATTGATCAG | TTTTTAAATAGGGTGCGTATGCCAT | TA | GAAGAGTCTTCCTTTACG |
| 12 | GAGGAGGGCAGCAAACGG | AA | TTTGCAGAATTAGGTTGAACCTCCG | ATGAAATCTTGACGCTTCAGTATAA | TA | GAAGAGTCTTCCTTTACG |
| 13 | GAGGAGGGCAGCAAACGG | AA | ACCATTTTGAGAACGGGAGTTGCGC | AACAGTTGTAATATATGCAATATTG | TA | GAAGAGTCTTCCTTTACG |
| 14 | GAGGAGGGCAGCAAACGG | AA | GCAGAAAAGCGGGCAACGTCGTCAG | CGTTCGGTGCCCGTTACATCTACAA | TA | GAAGAGTCTTCCTTTACG |
| 15 | GAGGAGGGCAGCAAACGG | AA | CTCAGGTCAACAATAATAAGTGTGG | TCAAACTCGTATGGATATCCAAAAT | TA | GAAGAGTCTTCCTTTACG |
| 16 | GAGGAGGGCAGCAAACGG | AA | GTTGCCGGGTAAGCTTAAAATTGCC | CTTCCCTATCAGTACCAAAATCAAA | TA | GAAGAGTCTTCCTTTACG |
| 17 | GAGGAGGGCAGCAAACGG | AA | TAAAACTCATCAATGCTGACGTATG | TTCTTCGAGCTCCAGTTGACAATTA | TA | GAAGAGTCTTCCTTTACG |
| 18 | GAGGAGGGCAGCAAACGG | AA | GAATCATCCAAAGGTTTAAAAGTGC | TTGAATAGTGCAGTGACTACATATG | TA | GAAGAGTCTTCCTTTACG |
| 19 | GAGGAGGGCAGCAAACGG | AA | TTTGAGCTTCAAAATCATAGCTGCC | ACTTTAATGGAGACTCCATATTGTT | TA | GAAGAGTCTTCCTTTACG |
| 20 | GAGGAGGGCAGCAAACGG | AA | GTGCTTGATTATTGTTGTAGTTTCC | CAATATTAGCCTGAAGCCCATATTC | TA | GAAGAGTCTTCCTTTACG |
| 21 | GAGGAGGGCAGCAAACGG | AA | GCCGTGTAGACTAAATTACTGTTTG | CTTATCTCCGACTTTTCTCCATTAA | TA | GAAGAGTCTTCCTTTACG |
| 22 | GAGGAGGGCAGCAAACGG | AA | AAGGTACCTTGTATTTCCCCATTTC | ACTCGACCATCGGGAGCGTTAAATA | TA | GAAGAGTCTTCCTTTACG |
| 23 | GAGGAGGGCAGCAAACGG | AA | CACCCTTTCCATTTTTGGCTACACG | TCAGAGTGTCAATACTGGCTGTAAA | TA | GAAGAGTCTTCCTTTACG |
| 24 | GAGGAGGGCAGCAAACGG | AA | ATGTCTTGGGCCGAGTAACGTTGTC | TGAACATTGCCTTGAATATTAATAT | TA | GAAGAGTCTTCCTTTACG |
| 25 | GAGGAGGGCAGCAAACGG | AA | TCCATTTTATTAACATCAAAATTCG | TGCTCACGCGAAATTTCCACATCGA | TA | GAAGAGTCTTCCTTTACG |
| 26 | GAGGAGGGCAGCAAACGG | AA | ATGAAGAAACCAGCCTCATACAGAC | ATATTTTCGGGAGAATGTTCATCAT | TA | GAAGAGTCTTCCTTTACG |
| 27 | GAGGAGGGCAGCAAACGG | AA | CATTCTTGTCGTTGTTTATTCCACC | GTTCGTATTGACCATATACAACCAA | TA | GAAGAGTCTTCCTTTACG |
| 28 | GAGGAGGGCAGCAAACGG | AA | AACCTCACTTAAGCTTGTTGAACTC | CTCTGCAACATTGCCGTTACCAGCT | TA | GAAGAGTCTTCCTTTACG |
| 29 | GAGGAGGGCAGCAAACGG | AA | AGTCCAGTGGATAGTACTTGTGCAT | GCAGAATGGGCTGACGAAACCACGC | TA | GAAGAGTCTTCCTTTACG |
| 30 | GAGGAGGGCAGCAAACGG | AA | AGGAATAAGTGGTGACTGGAAAATT | CTAGTTGGATAAGCGAGGTCTGTAT | TA | GAAGAGTCTTCCTTTACG |
| 31 | GAGGAGGGCAGCAAACGG | AA | TTCAATGAGTTTAACAATTTCGTCA | ACCAGCAGCAACACCATTATTGCCA | TA | GAAGAGTCTTCCTTTACG |
| 32 | GAGGAGGGCAGCAAACGG | AA | AATGTTTTTGTAACCCTAATGTTAG | AAACTGTATCGTCTGTAGTCAACGG | TA | GAAGAGTCTTCCTTTACG |
| 33 | GAGGAGGGCAGCAAACGG | AA | CTTAAGAAGTTCCAAGGATTTAGAT | ACGCAACTCTGAATCTTCGTTACGA | TA | GAAGAGTCTTCCTTTACG |
| 34 | GAGGAGGGCAGCAAACGG | AA | TGTACACAATGCGCTCTTCTTCCCG | TCTTTGCATTTCCTAGTCCCTTTAA | TA | GAAGAGTCTTCCTTTACG |
| 35 | GAGGAGGGCAGCAAACGG | AA | AGGGTCTCTTTGTCAACACTTTTCA | GGTAAGAGCTGAGAAGCCGCTTGAT | TA | GAAGAGTCTTCCTTTACG |
| 36 | GAGGAGGGCAGCAAACGG | AA | GTCTCACTATCAGATTGGCGTAGCA | TGTGGAAACGCAGCCAGTTCTAATA | TA | GAAGAGTCTTCCTTTACG |
| 37 | GAGGAGGGCAGCAAACGG | AA | TACCGCTTGGATTTTTCAGTTTTAA | AACCAGTTCCTGGTGAGTTTGCCTT | TA | GAAGAGTCTTCCTTTACG |
| 38 | GAGGAGGGCAGCAAACGG | AA | TGCTCGCGATGCGAACAGCTGTTTA | TTGCCGGACACCAAGCCACTGTTTA | TA | GAAGAGTCTTCCTTTACG |
| 39 | GAGGAGGGCAGCAAACGG | AA | AAAGAGTAGTCCAAGTCGCTGGAGT | AATGATACAACTGCGCGCTTAATAT | TA | GAAGAGTCTTCCTTTACG |
| 40 | GAGGAGGGCAGCAAACGG | AA | GCAGATCCGGACACTTTAAGACTTG | CCACAGTTTCCTTTCGCAAAAATTT | TA | GAAGAGTCTTCCTTTACG |

### apolpp (B4 initiator, pool E)

| Pair | Initiator | Spacer | Hybridzation | Hybridzation | Spacer | Initiator |
|---|---|---|---|---|---|---|
| 1 | CCTCAACCTACCTCCAAC | AA | GATAAAACATGTAAGCCCAGTTTAG | AAAAGATTTTTCCTCACGAAGGGAT | AT | TCTCACCATATTCGCTTC |
| 2 | CCTCAACCTACCTCCAAC | AA | CAAACACCCAGCCACCGTTATTAAG | CTTTTAATTTTTCAAAGTTTTGTGT | AT | TCTCACCATATTCGCTTC |
| 3 | CCTCAACCTACCTCCAAC | AA | TCGCACACCGATAATGCTTTTTGCC | CCAATTCTTATATTCTAAAGAATCA | AT | TCTCACCATATTCGCTTC |
| 4 | CCTCAACCTACCTCCAAC | AA | AAAACAATCACTCCGACTTGTACAT | AGAGCGGGATATCGTTTGGTTTCCT | AT | TCTCACCATATTCGCTTC |
| 5 | CCTCAACCTACCTCCAAC | AA | GTAATAGCACCCACTTATTAATCTG | GTCCTGCGCACTTAATGCAACGCGG | AT | TCTCACCATATTCGCTTC |
| 6 | CCTCAACCTACCTCCAAC | AA | TTATTGTCAAATTCAATTGCTGTGC | TCCTCTCTTTTGGAGCTTTTAAACT | AT | TCTCACCATATTCGCTTC |
| 7 | CCTCAACCTACCTCCAAC | AA | ATATAAGTGAATAGTGTAAAATCGG | ACATACTATGCCTACTCCGTACTCA | AT | TCTCACCATATTCGCTTC |
| 8 | CCTCAACCTACCTCCAAC | AA | CGCGAAATTCAGGCCGTCGAAAGTG | GAGAATATATTTGCAGTTGCCCGGA | AT | TCTCACCATATTCGCTTC |
| 9 | CCTCAACCTACCTCCAAC | AA | AAATGGAACGAACAGCCTTTATTAA | CTGGATAAGTGCTTCTAATCGAGAC | AT | TCTCACCATATTCGCTTC |
| 10 | CCTCAACCTACCTCCAAC | AA | TTACAATTTCTCCAATAGCTTTTCC | CTTCCTGAGCAGCGTCGTTTATAGG | AT | TCTCACCATATTCGCTTC |
| 11 | CCTCAACCTACCTCCAAC | AA | ATAATCGGTGCAATTGATTGATCAG | TTTTTAAATAGGGTGCGTATGCCAT | AT | TCTCACCATATTCGCTTC |
| 12 | CCTCAACCTACCTCCAAC | AA | TTTGCAGAATTAGGTTGAACCTCCG | ATGAAATCTTGACGCTTCAGTATAA | AT | TCTCACCATATTCGCTTC |
| 13 | CCTCAACCTACCTCCAAC | AA | ACCATTTTGAGAACGGGAGTTGCGC | AACAGTTGTAATATATGCAATATTG | AT | TCTCACCATATTCGCTTC |
| 14 | CCTCAACCTACCTCCAAC | AA | GCAGAAAAGCGGGCAACGTCGTCAG | CGTTCGGTGCCCGTTACATCTACAA | AT | TCTCACCATATTCGCTTC |
| 15 | CCTCAACCTACCTCCAAC | AA | CTCAGGTCAACAATAATAAGTGTGG | TCAAACTCGTATGGATATCCAAAAT | AT | TCTCACCATATTCGCTTC |
| 16 | CCTCAACCTACCTCCAAC | AA | GTTGCCGGGTAAGCTTAAAATTGCC | CTTCCCTATCAGTACCAAAATCAAA | AT | TCTCACCATATTCGCTTC |
| 17 | CCTCAACCTACCTCCAAC | AA | TAAAACTCATCAATGCTGACGTATG | TTCTTCGAGCTCCAGTTGACAATTA | AT | TCTCACCATATTCGCTTC |
| 18 | CCTCAACCTACCTCCAAC | AA | GAATCATCCAAAGGTTTAAAAGTGC | TTGAATAGTGCAGTGACTACATATG | AT | TCTCACCATATTCGCTTC |
| 19 | CCTCAACCTACCTCCAAC | AA | TTTGAGCTTCAAAATCATAGCTGCC | ACTTTAATGGAGACTCCATATTGTT | AT | TCTCACCATATTCGCTTC |
| 20 | CCTCAACCTACCTCCAAC | AA | GTGCTTGATTATTGTTGTAGTTTCC | CAATATTAGCCTGAAGCCCATATTC | AT | TCTCACCATATTCGCTTC |
| 21 | CCTCAACCTACCTCCAAC | AA | GCCGTGTAGACTAAATTACTGTTTG | CTTATCTCCGACTTTTCTCCATTAA | AT | TCTCACCATATTCGCTTC |
| 22 | CCTCAACCTACCTCCAAC | AA | AAGGTACCTTGTATTTCCCCATTTC | ACTCGACCATCGGGAGCGTTAAATA | AT | TCTCACCATATTCGCTTC |
| 23 | CCTCAACCTACCTCCAAC | AA | CACCCTTTCCATTTTTGGCTACACG | TCAGAGTGTCAATACTGGCTGTAAA | AT | TCTCACCATATTCGCTTC |
| 24 | CCTCAACCTACCTCCAAC | AA | ATGTCTTGGGCCGAGTAACGTTGTC | TGAACATTGCCTTGAATATTAATAT | AT | TCTCACCATATTCGCTTC |
| 25 | CCTCAACCTACCTCCAAC | AA | TCCATTTTATTAACATCAAAATTCG | TGCTCACGCGAAATTTCCACATCGA | AT | TCTCACCATATTCGCTTC |
| 26 | CCTCAACCTACCTCCAAC | AA | ATGAAGAAACCAGCCTCATACAGAC | ATATTTTCGGGAGAATGTTCATCAT | AT | TCTCACCATATTCGCTTC |
| 27 | CCTCAACCTACCTCCAAC | AA | CATTCTTGTCGTTGTTTATTCCACC | GTTCGTATTGACCATATACAACCAA | AT | TCTCACCATATTCGCTTC |
| 28 | CCTCAACCTACCTCCAAC | AA | AACCTCACTTAAGCTTGTTGAACTC | CTCTGCAACATTGCCGTTACCAGCT | AT | TCTCACCATATTCGCTTC |
| 29 | CCTCAACCTACCTCCAAC | AA | AGTCCAGTGGATAGTACTTGTGCAT | GCAGAATGGGCTGACGAAACCACGC | AT | TCTCACCATATTCGCTTC |
| 30 | CCTCAACCTACCTCCAAC | AA | AGGAATAAGTGGTGACTGGAAAATT | CTAGTTGGATAAGCGAGGTCTGTAT | AT | TCTCACCATATTCGCTTC |
| 31 | CCTCAACCTACCTCCAAC | AA | TTCAATGAGTTTAACAATTTCGTCA | ACCAGCAGCAACACCATTATTGCCA | AT | TCTCACCATATTCGCTTC |
| 32 | CCTCAACCTACCTCCAAC | AA | AATGTTTTTGTAACCCTAATGTTAG | AAACTGTATCGTCTGTAGTCAACGG | AT | TCTCACCATATTCGCTTC |
| 33 | CCTCAACCTACCTCCAAC | AA | CTTAAGAAGTTCCAAGGATTTAGAT | ACGCAACTCTGAATCTTCGTTACGA | AT | TCTCACCATATTCGCTTC |
| 34 | CCTCAACCTACCTCCAAC | AA | TGTACACAATGCGCTCTTCTTCCCG | TCTTTGCATTTCCTAGTCCCTTTAA | AT | TCTCACCATATTCGCTTC |
| 35 | CCTCAACCTACCTCCAAC | AA | AGGGTCTCTTTGTCAACACTTTTCA | GGTAAGAGCTGAGAAGCCGCTTGAT | AT | TCTCACCATATTCGCTTC |
| 36 | CCTCAACCTACCTCCAAC | AA | GTCTCACTATCAGATTGGCGTAGCA | TGTGGAAACGCAGCCAGTTCTAATA | AT | TCTCACCATATTCGCTTC |
| 37 | CCTCAACCTACCTCCAAC | AA | TACCGCTTGGATTTTTCAGTTTTAA | AACCAGTTCCTGGTGAGTTTGCCTT | AT | TCTCACCATATTCGCTTC |
| 38 | CCTCAACCTACCTCCAAC | AA | TGCTCGCGATGCGAACAGCTGTTTA | TTGCCGGACACCAAGCCACTGTTTA | AT | TCTCACCATATTCGCTTC |
| 39 | CCTCAACCTACCTCCAAC | AA | AAAGAGTAGTCCAAGTCGCTGGAGT | AATGATACAACTGCGCGCTTAATAT | AT | TCTCACCATATTCGCTTC |
| 40 | CCTCAACCTACCTCCAAC | AA | GCAGATCCGGACACTTTAAGACTTG | CCACAGTTTCCTTTCGCAAAAATTT | AT | TCTCACCATATTCGCTTC |

### beat VII

| Pair | Initiator | Spacer | Hybridzation | Hybridzation | Spacer | Initiator |
|---|---|---|---|---|---|---|
| 1 | GAGGAGGGCAGCAAACGG | AA | TCAGTGTAGTAGATTGGCCAAGCAG | GTATATTTAATATGCTGTATATCCC | TA | GAAGAGTCTTCCTTTACG |
| 2 | GAGGAGGGCAGCAAACGG | AA | CAGTGTACATATAAATATTCGTGGC | ACTTTACACTTAATTCTGTCTGCTG | TA | GAAGAGTCTTCCTTTACG |
| 3 | GAGGAGGGCAGCAAACGG | AA | TTTATGTTCGATAAACAATGGGAAC | TCCACGTGATTTGTGCATGCGATTA | TA | GAAGAGTCTTCCTTTACG |
| 4 | GAGGAGGGCAGCAAACGG | AA | CCGGACAAAATGTCTTTGTCTGGCC | TGATAAAGGTAGATGTGACTCTGAT | TA | GAAGAGTCTTCCTTTACG |
| 5 | GAGGAGGGCAGCAAACGG | AA | TATATGACTTGGAGAAGCCTCAACA | TGGCAGCCGAGTCAATATCCACTTA | TA | GAAGAGTCTTCCTTTACG |
| 6 | GAGGAGGGCAGCAAACGG | AA | GGGATCCGAAAATGTTACAACATTT | CTCGAGAGGCTGGGCTCTCATTGTA | TA | GAAGAGTCTTCCTTTACG |
| 7 | GAGGAGGGCAGCAAACGG | AA | CGATAGAGGTTGCATAATTAGCGAC | ACATAGATACACATACATAGATATG | TA | GAAGAGTCTTCCTTTACG |
| 8 | GAGGAGGGCAGCAAACGG | AA | AAGACAACACGCAGTTTGCATATTT | TTGCTGAGACAACAGACGACTGAAT | TA | GAAGAGTCTTCCTTTACG |
| 9 | GAGGAGGGCAGCAAACGG | AA | GTGATGCGTTGATAGGGTGTTTGCA | CTGCGGATGTGTTGGCGAGTGTAGA | TA | GAAGAGTCTTCCTTTACG |
| 10 | GAGGAGGGCAGCAAACGG | AA | CATAGGAAGACGTCAATACCTGCAT | TTTTTCTGTTAATAATAGACAGCGT | TA | GAAGAGTCTTCCTTTACG |
| 11 | GAGGAGGGCAGCAAACGG | AA | ACTCACAATGAGGACAACCCTTGGA | ACTGGCCAAGAAACTACATGTACGT | TA | GAAGAGTCTTCCTTTACG |
| 12 | GAGGAGGGCAGCAAACGG | AA | TGTTTGGTTTTGCATTTCGCCGTCG | GCAGGTTTAGTTGTTATACTTAATT | TA | GAAGAGTCTTCCTTTACG |
| 13 | GAGGAGGGCAGCAAACGG | AA | CATAGTTACATTGTTATGCTACGTG | TTCCGTGGTTGCTACCATCTTGCTT | TA | GAAGAGTCTTCCTTTACG |
| 14 | GAGGAGGGCAGCAAACGG | AA | TTGTTAAGCAATGCGACTAAGCCGC | TACACACATGAAATGCATACGTACT | TA | GAAGAGTCTTCCTTTACG |
| 15 | GAGGAGGGCAGCAAACGG | AA | ATGAAAATAAATAAATAAGCGTCGG | CAACGAACCCAAAATGAAGGCTCCA | TA | GAAGAGTCTTCCTTTACG |
| 16 | GAGGAGGGCAGCAAACGG | AA | TTCCTTCGGGTTCTGCGTGTTATTT | CGCAGATTGCGCCTCTTGCGGCTCC | TA | GAAGAGTCTTCCTTTACG |
| 17 | GAGGAGGGCAGCAAACGG | AA | ACGGTACTGCTGGCTCCTAAACTCA | GGTACTTTGGTACGGGGCATTCGGG | TA | GAAGAGTCTTCCTTTACG |
| 18 | GAGGAGGGCAGCAAACGG | AA | CACTAAGATCATGCATTTTAGGTAC | TGTAATTCGATTTGGTATGCACTAC | TA | GAAGAGTCTTCCTTTACG |
| 19 | GAGGAGGGCAGCAAACGG | AA | TCTCCCAAGCATCTGACAAGGTTGA | TTTTTGGTGCGGGATGGGGCTGTGC | TA | GAAGAGTCTTCCTTTACG |
| 20 | GAGGAGGGCAGCAAACGG | AA | TCCCGAAACTTATGTTCAATTAGCT | TTTGTTCTGAGTTGCAGTCGCCTCC | TA | GAAGAGTCTTCCTTTACG |
| 21 | GAGGAGGGCAGCAAACGG | AA | AATTACAAAAATTTCTCTTTGGCAT | ACCCCATTTCCCGTCCGACTGCAAC | TA | GAAGAGTCTTCCTTTACG |
| 22 | GAGGAGGGCAGCAAACGG | AA | GGCGAGCACAAATGCTAAAAATGTT | TGTGAACTATTTAAATTGGCCGACC | TA | GAAGAGTCTTCCTTTACG |
| 23 | GAGGAGGGCAGCAAACGG | AA | TGCTGCTTTTGTCCTTGATGGGATG | TTTTTCCGATGCTGCTGCTGCTTTT | TA | GAAGAGTCTTCCTTTACG |
| 24 | GAGGAGGGCAGCAAACGG | AA | TGCATTCCCCATTTGGTGCCATCTA | GTGGTCTTCTCCTCTCCTCCGTTCG | TA | GAAGAGTCTTCCTTTACG |
| 25 | GAGGAGGGCAGCAAACGG | AA | GGGATGCCCAGCTCCTTGTTCATGT | AGCTGGTTCATGGGTCCAGCGTTTG | TA | GAAGAGTCTTCCTTTACG |
| 26 | GAGGAGGGCAGCAAACGG | AA | CCATGTTCTTCTTGTTGCTGTCGTA | TCTCCATCTGCTGCTGCTTCTTCAG | TA | GAAGAGTCTTCCTTTACG |
| 27 | GAGGAGGGCAGCAAACGG | AA | GCCAGCGATCCAAACGGATTGTTGT | TGGAACTCGCCCACGTCATGGTAGT | TA | GAAGAGTCTTCCTTTACG |
| 28 | GAGGAGGGCAGCAAACGG | AA | CTTTTGTCGTAGCGATCCATGAATT | GCTCGTGCTCCTGCTGGCCATCGTA | TA | GAAGAGTCTTCCTTTACG |
| 29 | GAGGAGGGCAGCAAACGG | AA | ACTGGTTGAAGTACTGCTGCTGGAA | CAACGGGGATGTGATACTGCTGATG | TA | GAAGAGTCTTCCTTTACG |
| 30 | GAGGAGGGCAGCAAACGG | AA | GCACGTACCTGTCCTCCACCTTTTT | CATTGAAGGAGAACACGTGGTGGGT | TA | GAAGAGTCTTCCTTTACG |
| 31 | GAGGAGGGCAGCAAACGG | AA | GCGTTCGCTTGCGAAACTTAATGGT | CGAAAAGTTTCTCGCCCACCGCGAA | TA | GAAGAGTCTTCCTTTACG |
| 32 | GAGGAGGGCAGCAAACGG | AA | CATCGGCGCTGGCCTTGGTAAAAAT | TCTGGGGCAAGAACACGCTCATCAG | TA | GAAGAGTCTTCCTTTACG |
| 33 | GAGGAGGGCAGCAAACGG | AA | TGGCCGGATAGATCGAACTGGACAT | GTGTCCGTGGAGACCTCGCAGTAGA | TA | GAAGAGTCTTCCTTTACG |
| 34 | GAGGAGGGCAGCAAACGG | AA | ATTTCGGCTCCCTCGATGGTTGAGT | TGCTGCTCGTTGGAATTGTCCCAGT | TA | GAAGAGTCTTCCTTTACG |
| 35 | GAGGAGGGCAGCAAACGG | AA | ATTCAAAGAACTTGCCCTTGTCAAC | GGAATGGCGGATTTCGTCCGTTTAT | TA | GAAGAGTCTTCCTTTACG |

### borderless

| Pair | Initiator | Spacer | Hybridzation | Hybridzation | Spacer | Initiator |
|---|---|---|---|---|---|---|
| 1 | CCTCAACCTACCTCCAAC | AA | AGTTTATGTTACAGATAGCAGCTAG | CGGTTAGTTTAGCGTTACAATTAGT | AT | TCTCACCATATTCGCTTC |
| 2 | CCTCAACCTACCTCCAAC | AA | TGGTGCACGGTGAACAACCCACCCA | TATATTAAAAGTCAGATTCCCACGG | AT | TCTCACCATATTCGCTTC |
| 3 | CCTCAACCTACCTCCAAC | AA | GGCCAACATGGTTGCTGTCAAATCG | CCTCATCTATCGCCTCTTAGTTTCA | AT | TCTCACCATATTCGCTTC |
| 4 | CCTCAACCTACCTCCAAC | AA | TTAGGCAGCTCCGCTTCACGTATAA | CGGAGCTATCCTTGGCGAACAGATG | AT | TCTCACCATATTCGCTTC |
| 5 | CCTCAACCTACCTCCAAC | AA | TTGAGGTGGCGCAGCTGGTAGTAGT | ACCTGAACCTTGAACAGTGTGTCCT | AT | TCTCACCATATTCGCTTC |
| 6 | CCTCAACCTACCTCCAAC | AA | TGGGCATCAAAGTCATCCGCTCTAA | ACCTGACCCAGATCGTGTTGCAGCT | AT | TCTCACCATATTCGCTTC |
| 7 | CCTCAACCTACCTCCAAC | AA | TTGCTGAACATGCCGTCTCCGTACT | GAGGGTAGCGTTTGAAAGCGGAACT | AT | TCTCACCATATTCGCTTC |
| 8 | CCTCAACCTACCTCCAAC | AA | CCTTTCCTGGCTGAAGGTGCTGCAC | CCTGGCTGAGCACCATAAATTCGTA | AT | TCTCACCATATTCGCTTC |
| 9 | CCTCAACCTACCTCCAAC | AA | AGCGTTCGCCATTCGGGTGCCTCCA | TCCATCACCTTTTTATCCAACACGC | AT | TCTCACCATATTCGCTTC |
| 10 | CCTCAACCTACCTCCAAC | AA | TGGGACGAAGGTATCCTGGCTGCCA | GTCGATACCAAACAGTGTACTCCAA | AT | TCTCACCATATTCGCTTC |
| 11 | CCTCAACCTACCTCCAAC | AA | AGAACAGGCTGCCGTTCATCTTGTA | CGGCATGGTTCTCGTCCACTTTGGC | AT | TCTCACCATATTCGCTTC |
| 12 | CCTCAACCTACCTCCAAC | AA | CGTCCTTCTCCCAGCGCAAATTCTT | GCACATTGTACGAGTCGAAGAGCAG | AT | TCTCACCATATTCGCTTC |
| 13 | CCTCAACCTACCTCCAAC | AA | GTAAATCACCTTGGCCTTATATTGT | GTACGGCAGGAAAACCTCCGGCGGG | AT | TCTCACCATATTCGCTTC |
| 14 | CCTCAACCTACCTCCAAC | AA | TGCCATCGGGACCCATGTAGAATCT | TCATCATGGTGGGATCGATGCTGAG | AT | TCTCACCATATTCGCTTC |
| 15 | CCTCAACCTACCTCCAAC | AA | GGATGCTTCATCACGCAGTGAAAGA | TTGTACCAGGAGGCCTGCGAGTTCT | AT | TCTCACCATATTCGCTTC |
| 16 | CCTCAACCTACCTCCAAC | AA | TGGGAAAGCTAACTTGGCAGTGGTA | CATTGTTGCGAACCGAGGGCGAGCG | AT | TCTCACCATATTCGCTTC |
| 17 | CCTCAACCTACCTCCAAC | AA | ACCAGGTGCAATCGGCCATTGAATA | GCGCGTCCAAATTCCGGATGGTTCT | AT | TCTCACCATATTCGCTTC |
| 18 | CCTCAACCTACCTCCAAC | AA | AGGTGAAGATTTTCTTGTTATCCTT | CACTGGTGGAGGTCTCCTGCTCGTA | AT | TCTCACCATATTCGCTTC |
| 19 | CCTCAACCTACCTCCAAC | AA | CAGGATATGTTCATTAAGAGAATGA | CGCCTGTGATCGCGGGCAGCGCTCG | AT | TCTCACCATATTCGCTTC |
| 20 | CCTCAACCTACCTCCAAC | AA | TGCGTTTCGCTGGCATTTCGATGTT | CGGATCCGGATTGCCGGAATGTTCT | AT | TCTCACCATATTCGCTTC |
| 21 | CCTCAACCTACCTCCAAC | AA | TCTGCACTTGTTGCACTCAGTCACA | TTTGATGTTTTCTTGCGTTTCTCAC | AT | TCTCACCATATTCGCTTC |
| 22 | CCTCAACCTACCTCCAAC | AA | GTAGGCCAGTATGCACAAAATCAAA | GCGCGGTGTCTTGCATAATGTTTCC | AT | TCTCACCATATTCGCTTC |
| 23 | CCTCAACCTACCTCCAAC | AA | CTCGTTTGCGGAATTCGGAATGTAT | TCAATTGGGGCCCTCGACGAGGAGC | AT | TCTCACCATATTCGCTTC |
| 24 | CCTCAACCTACCTCCAAC | AA | TTACGATATCTTTGTGCGCAGTTCC | AGAGGCGCACGAAACCCGATCCGTT | AT | TCTCACCATATTCGCTTC |
| 25 | CCTCAACCTACCTCCAAC | AA | GCGGCAGCTATGTTGTGGCGATCTT | CAGGTATTGTGTGGCATAGTATATG | AT | TCTCACCATATTCGCTTC |

### calcineurin A1

| Pair | Initiator | Spacer | Hybridzation | Hybridzation | Spacer | Initiator |
|---|---|---|---|---|---|---|
| 1 | CCTCGTAAATCCTCATCA | AA | GCACATTACTGGTGGTGGCCGATTT | AGAACTTCTTGGCGGTGAATCCTGC | AA | ATCATCCAGTAAACCGCC |
| 2 | CCTCGTAAATCCTCATCA | AA | TGCGGTATCATTACTGCTGGTGTTG | AGTCGTCCTGCTCGTTTTGGTTACG | AA | ATCATCCAGTAAACCGCC |
| 3 | CCTCGTAAATCCTCATCA | AA | GTGTTGCTGTTATTGTTGTTGTTGT | ATGTCCTTTGTCGTCGTGGTGCTGC | AA | ATCATCCAGTAAACCGCC |
| 4 | CCTCGTAAATCCTCATCA | AA | ATCTTGCCAATGGCCCGAATCTTAT | TCCCTGAGGATGGAGAAGACGCGCG | AA | ATCATCCAGTAAACCGCC |
| 5 | CCTCGTAAATCCTCATCA | AA | TGGAAGCTGGCTTACTGGGCGTATT | GAATTATCTCTTTGCGCAGCGCCGA | AA | ATCATCCAGTAAACCGCC |
| 6 | CCTCGTAAATCCTCATCA | AA | CGGCACGAGCACAATTTTCTTGCGT | GTTATTGTTATTATTGCTCGCGTTC | AA | ATCATCCAGTAAACCGCC |
| 7 | CCTCGTAAATCCTCATCA | AA | TCGTCTGAGCAAATGTTCAGGATGT | TCGTCATCGGGTCCCGCCACCAGTT | AA | ATCATCCAGTAAACCGCC |
| 8 | CCTCGTAAATCCTCATCA | AA | GACCATGTGAAGACGTCCATGAAGT | GTTACCTTTTCGCCCACGAAGGGCA | AA | ATCATCCAGTAAACCGCC |
| 9 | CCTCGTAAATCCTCATCA | AA | TGAACTGGCGGATATTCATCACGTT | GCAGCCAGTAGGGATGTGGCGAGCA | AA | ATCATCCAGTAAACCGCC |
| 10 | CCTCGTAAATCCTCATCA | AA | GTTGTAAACATCCAGATAATTGGGG | CTCGTACTTGAGCACAGCTGCCTTG | AA | ATCATCCAGTAAACCGCC |
| 11 | CCTCGTAAATCCTCATCA | AA | CCCGTAACCTGGTTCTTCCGATACA | GAGAATATGGTGATCAGCGAGGGGA | AA | ATCATCCAGTAAACCGCC |
| 12 | CCTCGTAAATCCTCATCA | AA | TGCAGGAACTCACAGCAGGCCGAGT | CTGACTATCGACAGCAGGTTGTTCT | AA | ATCATCCAGTAAACCGCC |
| 13 | CCTCGTAAATCCTCATCA | AA | TTGGTCTTCTCGTTGCCAAAGTCCT | CGCACCGAGTTGTGCGAGAAGAACT | AA | ATCATCCAGTAAACCGCC |
| 14 | CCTCGTAAATCCTCATCA | AA | GTCTTGATGTCGTCGAGTGTGAAAA | GCCGGTGGTTCCCTGAAGCGATTCA | AA | ATCATCCAGTAAACCGCC |
| 15 | CCTCGTAAATCCTCATCA | AA | GCGTCGTAAATACTTTCCGAATACT | GGCAGGCAGTCGAACGCCTCCATGC | AA | ATCATCCAGTAAACCGCC |
| 16 | CCTCGTAAATCCTCATCA | AA | ACTCTGTGAGATGCCTGCACTCGTG | TGATGCATTCCTGCTTGAAGGTAAA | AA | ATCATCCAGTAAACCGCC |
| 17 | CCTCGTAAATCCTCATCA | AA | AGGTTATTTTCAGGGACCACAGGTA | CTCGCAGCAGGGAAAGTGTGGTGGG | AA | ATCATCCAGTAAACCGCC |
| 18 | CCTCGTAAATCCTCATCA | AA | GTCCACGTAGTCGCCTAGAAAAAGG | AACGCACTCAATACTGAAGTATCCC | AA | ATCATCCAGTAAACCGCC |
| 19 | CCTCGTAAATCCTCATCA | AA | AGAACTGCCCGTGGATGTCACCGCA | CCACTTCGAAGAGCTTTACGAGATC | AA | ATCATCCAGTAAACCGCC |
| 20 | CCTCGTAAATCCTCATCA | AA | GATCGTCGTATACCTCCGACATGGT | CGTCGAAGTTGGGTTTTCCGGTCTT | AA | ATCATCCAGTAAACCGCC |
| 21 | CCTCGTAAATCCTCATCA | AA | ATGCGCTCCCTGGTTTTTGTGTACT | GTGGGCGGAAGTGGAACATCATCGA | AA | ATCATCCAGTAAACCGCC |
| 22 | CCTCGTAAATCCTCATCA | AA | TCGGGGATTTCGAGGAATTCGACGA | TTTTCTGTTTGTTTCCACTGGCGGC | AA | ATCATCCAGTAAACCGCC |
| 23 | CCTCGTAAATCCTCATCA | AA | TTTTGATAGCGATGATTTCCTCGTC | CTTTGCCGATTTGTCTGAACTAGAA | AA | ATCATCCAGTAAACCGCC |

# CG6055

| Pair | Initiator | Spacer | Hybridzation | Hybridzation | Spacer | Initiator |
|---|---|---|---|---|---|---|
| 1 | CCTCGTAAATCCTCATCA | AA | CACAGAAATGTTATTACAAATTTTG | CAATTTTGTTTATGATGCGCGGATC | AA | ATCATCCAGTAAACCGCC |
| 2 | CCTCGTAAATCCTCATCA | AA | ACAAACGAATGTTACGCGTTTTGGA | ATAGAACTTTTGCTGCGAACTAACC | AA | ATCATCCAGTAAACCGCC |
| 3 | CCTCGTAAATCCTCATCA | AA | GTAAATTTTAATCATCTCCCTCAGG | ATGTTCCGTTTAAGGCTACATAAAT | AA | ATCATCCAGTAAACCGCC |
| 4 | CCTCGTAAATCCTCATCA | AA | TGGGGTTCCGGGAGCGGACGAAGTT | AGGCTAGTTAGAATTACAAGCGGAC | AA | ATCATCCAGTAAACCGCC |
| 5 | CCTCGTAAATCCTCATCA | AA | TGGCTTGATGTGATGGCAGGCCACG | TAGCTCATCGGAGTCCTCGCACACA | AA | ATCATCCAGTAAACCGCC |
| 6 | CCTCGTAAATCCTCATCA | AA | TTGTTCAGGATGGACAGGCAGCTCT | TGCCACTTGATGCCATCGTTGTAGA | AA | ATCATCCAGTAAACCGCC |
| 7 | CCTCGTAAATCCTCATCA | AA | TTGGTATCCGCCCGTTGAAGACCAA | GGCCTCCCTATTATCGGGCTGCGGC | AA | ATCATCCAGTAAACCGCC |
| 8 | CCTCGTAAATCCTCATCA | AA | TTGGGAGGCTGCAGATCGGGCCTGT | GATCCCGACCAGAACCAACCGTTCT | AA | ATCATCCAGTAAACCGCC |
| 9 | CCTCGTAAATCCTCATCA | AA | AGGTCCAAATGTATCGCACATTGCC | AGCCAGCGAAGTTGCACTTCCTGCC | AA | ATCATCCAGTAAACCGCC |
| 10 | CCTCGTAAATCCTCATCA | AA | CTCCTGCGGCGTCTCCAGGGAAACG | GGCAATCCTCTGCTTCACAAAGTCA | AA | ATCATCCAGTAAACCGCC |
| 11 | CCTCGTAAATCCTCATCA | AA | CTCGCATCCAGCCAATCCACCTCCA | TCCATGCAGTGCCTGCGGCAGATGT | AA | ATCATCCAGTAAACCGCC |
| 12 | CCTCGTAAATCCTCATCA | AA | TACGAGTGGGACACGCCCCTGGCAT | GTAGGAGCATGCTCCCAGCTGAAGA | AA | ATCATCCAGTAAACCGCC |
| 13 | CCTCGTAAATCCTCATCA | AA | CATCAGTGCACGCAGCACAGACATA | CGCCACGGTGGCACCAAAGGCCAGA | AA | ATCATCCAGTAAACCGCC |
| 14 | CCTCGTAAATCCTCATCA | AA | ACAAAGCGGGGCTCAAAACTCGATA | GGATAGCTTTAGATTGAAACGAACG | AA | ATCATCCAGTAAACCGCC |
| 15 | CCTCGTAAATCCTCATCA | AA | TGTTGTTGTGGGAACAACCGCACGC | GAACAGCACAGATATCCAAGCTTAA | AA | ATCATCCAGTAAACCGCC |

# CG67443

| Pair | Initiator | Spacer | Hybridzation | Hybridzation | Spacer | Initiator |
|---|---|---|---|---|---|---|
| 1 | CCTCGTAAATCCTCATCA | AA | TTATTTGAGCTGGGTTATTAGCCCA | ATGGCTAAAACGGAATGAACGTCTG | AA | ATCATCCAGTAAACCGCC |
| 2 | CCTCGTAAATCCTCATCA | AA | TTAATAGCTAGTGTTAGAGAGGTGT | GCAACATTATCGTATACCAAATCCC | AA | ATCATCCAGTAAACCGCC |
| 3 | CCTCGTAAATCCTCATCA | AA | AACCACCAGGGGCAGCTGTGGGTGG | AAGGTGTGGGTAAGGATAAGATAAT | AA | ATCATCCAGTAAACCGCC |
| 4 | CCTCGTAAATCCTCATCA | AA | GGCAGTTACTGCAGGTTGTGGCGAT | AGCTGTAGGTGGGGCAGCTGTGGTT | AA | ATCATCCAGTAAACCGCC |
| 5 | CCTCGTAAATCCTCATCA | AA | CCTGCGGAGCTACAGTGCTTCCATT | GTGAACTCCCTGGAGCTGCTCCACC | AA | ATCATCCAGTAAACCGCC |
| 6 | CCTCGTAAATCCTCATCA | AA | TACCGTGCAGTTACGACAAATCAAA | TGTGGCGTTTGGAGCAGCTTGAACA | AA | ATCATCCAGTAAACCGCC |
| 7 | CCTCGTAAATCCTCATCA | AA | CTATCATTGTCATAGAGTATATCCT | TTCTCATCGCCGGCATCATCACTAT | AA | ATCATCCAGTAAACCGCC |
| 8 | CCTCGTAAATCCTCATCA | AA | TCGTCTTGACAGAGGTTTCCTCCAA | GGATAGTCCACCAGTCGGTATCGCG | AA | ATCATCCAGTAAACCGCC |
| 9 | CCTCGTAAATCCTCATCA | AA | CATAAGGCAAAGGCCCACTGCGAAT | CCTAGCCTCAATGCTCAGAGAACTA | AA | ATCATCCAGTAAACCGCC |

# CG6356

| Pair | Initiator | Spacer | Hybridzation | Hybridzation | Spacer | Initiator |
|---|---|---|---|---|---|---|
| 1 | CCTCAACCTACCTCCAAC | AA | TAGCGAAAGTCAGTTGGAGGTACAG | GTGTTTCGGATTGTAAAACAATTTA | AT | TCTCACCATATTCGCTTC |
| 2 | CCTCAACCTACCTCCAAC | AA | TGGAACTTAACCTATTGCTGTCATC | AGTAGTTAATCAGTTTAGTCGTGGA | AT | TCTCACCATATTCGCTTC |
| 3 | CCTCAACCTACCTCCAAC | AA | TTCCCGTTTCCGGAAGTGTGAGAGT | TCTCCACTTTGTCGGACAGCTTACC | AT | TCTCACCATATTCGCTTC |
| 4 | CCTCAACCTACCTCCAAC | AA | GCAAATCGTTGTTGGAATGTACCAG | TAGGCCAGCTACTAGAACGCAGCAG | AT | TCTCACCATATTCGCTTC |
| 5 | CCTCAACCTACCTCCAAC | AA | GAGTTTCTTATCTGGGTGGGATACA | GCGAAGGTGGAGCAGGTTCCCAATG | AT | TCTCACCATATTCGCTTC |
| 6 | CCTCAACCTACCTCCAAC | AA | ACAGGGACGAGATAGGAGGGTATAT | CGACGGCCCGTAAAGCGCAGCATTA | AT | TCTCACCATATTCGCTTC |
| 7 | CCTCAACCTACCTCCAAC | AA | ATCTGGACACCGACAAATTGGCGGC | CAAGACCGGTGCCTATAATGTATAG | AT | TCTCACCATATTCGCTTC |
| 8 | CCTCAACCTACCTCCAAC | AA | AGTGACACAGAACATGAAATACGAG | GAGAGCCGTCACATAGTAACTCAGC | AT | TCTCACCATATTCGCTTC |
| 9 | CCTCAACCTACCTCCAAC | AA | GCATACAGCTCGATGATGGACTTCA | AAGATCATCCTGCGCACCTTGGAGT | AT | TCTCACCATATTCGCTTC |
| 10 | CCTCAACCTACCTCCAAC | AA | AGAATCCCTCTTCGGGCTCCAGCGA | GACCATTCTTGAGTTTCTGTCCAAA | AT | TCTCACCATATTCGCTTC |
| 11 | CCTCAACCTACCTCCAAC | AA | GCAACGGCGGGCTGAATTTGGTAAT | TTATCGCTGGCGACTTGAGATGGGG | AT | TCTCACCATATTCGCTTC |
| 12 | CCTCAACCTACCTCCAAC | AA | CGCGATCATATCTTTGGTTCGTGAT | TGGGCTGTCTGAAGAGAATCTTGTA | AT | TCTCACCATATTCGCTTC |
| 13 | CCTCAACCTACCTCCAAC | AA | TTCAGAATGAGCAGCAGCGAAGGAA | CGGGGTGACTCACTCATCAGGTAGC | AT | TCTCACCATATTCGCTTC |
| 14 | CCTCAACCTACCTCCAAC | AA | ACGTAGCCCACAATGGCCAGGATGA | AATTGCAGATGCCTCCACGGTTGGA | AT | TCTCACCATATTCGCTTC |
| 15 | CCTCAACCTACCTCCAAC | AA | TGGACATCCAGGAACGATGCTTCAA | CCACCGGATAGGACGCTGAGAAGGC | AT | TCTCACCATATTCGCTTC |
| 16 | CCTCAACCTACCTCCAAC | AA | TGGATAGAACAGACCAGAGGAGGCT | GATGTTCTCCACGATCATGGTAAAT | AT | TCTCACCATATTCGCTTC |
| 17 | CCTCAACCTACCTCCAAC | AA | GGTACAGAAGAAGCTGCCCACAATA | CAGGAAGAGGCTGTACCAGGGCGAG | AT | TCTCACCATATTCGCTTC |
| 18 | CCTCAACCTACCTCCAAC | AA | CAAATTTGTCGGCCAGGATGCCAAA | CGCCCAGGGTGAATGAGGTTTTGCG | AT | TCTCACCATATTCGCTTC |
| 19 | CCTCAACCTACCTCCAAC | AA | TTCAGGGCTGCCTCGTAGGACATCT | GAAATCTGCGTCAGATGGCCGGTAT | AT | TCTCACCATATTCGCTTC |
| 20 | CCTCAACCTACCTCCAAC | AA | TAGGATGACACATGGAATCCGTTCA | TCATCGGGTAGGTGGCCCAGCAGCG | AT | TCTCACCATATTCGCTTC |
| 21 | CCTCAACCTACCTCCAAC | AA | AATGCCCCGCCACTGCCAACGACTT | ACACACAGGAAGAAGATTGCCCACA | AT | TCTCACCATATTCGCTTC |
| 22 | CCTCAACCTACCTCCAAC | AA | TTTCCTCGCTGATCAGCTCGCGATT | GCTGCTTGGTGAGCAGATCCTTCAT | AT | TCTCACCATATTCGCTTC |
| 23 | CCTCAACCTACCTCCAAC | AA | GATGCTAGCTGCTACTTCTACTTCT | CTCCGGATTCTCCGTGGATTGAGTC | AT | TCTCACCATATTCGCTTC |
| 24 | CCTCAACCTACCTCCAAC | AA | CGTTTGTCGCAGACTCTGAAGACGA | TGGGCTATTCAAAATTCAAAATCTT | AT | TCTCACCATATTCGCTTC |
| 25 | CCTCAACCTACCTCCAAC | AA | AGGCAGACGACTGACCCCACGCAAA | TGATGATGATGTTGAGGATGTTGTA | AT | TCTCACCATATTCGCTTC |
| 26 | CCTCAACCTACCTCCAAC | AA | CCTGTGCCACTGAACTTGAATTCCT | AATGTTCAGATTGCTCGGGAGGTTC | AT | TCTCACCATATTCGCTTC |
| 27 | CCTCAACCTACCTCCAAC | AA | TCGCTGAAGTGCATTCACTATGGGC | GGGGCTTAGGAACCCAGTGTTGTAG | AT | TCTCACCATATTCGCTTC |
| 28 | CCTCAACCTACCTCCAAC | AA | ACTTTCGCCCTCGACTACAGGTCGA | GGTGGGAGATAAGAGATGTACACTT | AT | TCTCACCATATTCGCTTC |
| 29 | CCTCAACCTACCTCCAAC | AA | TGGGAGCGCTTCACTTTTGACGGAC | TTTTTCAATTGCCAATGCAACAGCA | AT | TCTCACCATATTCGCTTC |
| 30 | CCTCAACCTACCTCCAAC | AA | TTTACCAACGCCGCTTGTTGTTTGC | GACACGTTTTTCTTTCTCTGCTCTT | AT | TCTCACCATATTCGCTTC |
| 31 | CCTCAACCTACCTCCAAC | AA | TTTCACTGCCTTGTATCTATGGTGC | ATGATATATTTTCCGTTGGGATATT | AT | TCTCACCATATTCGCTTC |
| 32 | CCTCAACCTACCTCCAAC | AA | TCGCCGATCTTCGCTATCAACTACT | CGTGCAGTTGCGTTTGCCTGCCGGT | AT | TCTCACCATATTCGCTTC |

# CG9701

| Pair | Initiator | Spacer | Hybridzation | Hybridzation | Spacer | Initiator |
|---|---|---|---|---|---|---|
| 1 | GAGGAGGGCAGCAAACGG | AA | ACTGACAGCACCGCTAGCACCACTT | CAGCAGTAGAATGCCCATCAGACTC | TA | GAAGAGTCTTCCTTTACG |
| 2 | GAGGAGGGCAGCAAACGG | AA | TCTGTAGCTCCAATCGATGGTGTTT | CAGCTGCTGCTCCTCATCCAGCTTG | TA | GAAGAGTCTTCCTTTACG |
| 3 | GAGGAGGGCAGCAAACGG | AA | CTGGTGCGTTGCGGTGAGTTGAAGT | AATACTCTCGCCGAGATCTTGGGCG | TA | GAAGAGTCTTCCTTTACG |
| 4 | GAGGAGGGCAGCAAACGG | AA | AGCCTGCCTTCCACTCATAGCTGTC | CATGATAGAGACCGAATTTCTCCGA | TA | GAAGAGTCTTCCTTTACG |
| 5 | GAGGAGGGCAGCAAACGG | AA | TTCGCTCCATCCTCCATGGCATCCA | AGACTCCAGGCGATGTAGCCACTGA | TA | GAAGAGTCTTCCTTTACG |
| 6 | GAGGAGGGCAGCAAACGG | AA | GCATAGTCCTCCAGACCACCGCGAT | AGATAGAGATTGTAGTAGTCCACGC | TA | GAAGAGTCTTCCTTTACG |
| 7 | GAGGAGGGCAGCAAACGG | AA | CGGGCGCATTGTACTCCCGGTGTAT | TGACTCCGTTCTCCGTAACGATTAT | TA | GAAGAGTCTTCCTTTACG |
| 8 | GAGGAGGGCAGCAAACGG | AA | ATAGACCTTGAGCCAAACGGATCCG | CATCAACAGATTGTACATTCCCTTG | TA | GAAGAGTCTTCCTTTACG |
| 9 | GAGGAGGGCAGCAAACGG | AA | CGTGATTAAAGGATGGAACTGGAAA | CCTCCTGGCTCTCGACCACGCCCAT | TA | GAAGAGTCTTCCTTTACG |
| 10 | GAGGAGGGCAGCAAACGG | AA | GACCAGGTTGCTGGTATAGGAATTG | GCCGGTATTATTGTGACCATTGGAG | TA | GAAGAGTCTTCCTTTACG |
| 11 | GAGGAGGGCAGCAAACGG | AA | ATGAATCTCCTCAGTGGTGAACTCA | GAAGAAGTCGGAGGTGCCGCGTATG | TA | GAAGAGTCTTCCTTTACG |
| 12 | GAGGAGGGCAGCAAACGG | AA | TCAAGTTCCGAATGCGCTCGATCAT | ATCGTGCTCCGAAACCCTGCTCCTT | TA | GAAGAGTCTTCCTTTACG |
| 13 | GAGGAGGGCAGCAAACGG | AA | GATGGGATGACCGAACCAGCCTACA | CTTGGGATAGTTACCGTGCTTGGAA | TA | GAAGAGTCTTCCTTTACG |
| 14 | GAGGAGGGCAGCAAACGG | AA | AGGTAGGCGGGAATGCCTGGATAGT | TGAGCCTTCAGCAGATTGTGACCAC | TA | GAAGAGTCTTCCTTTACG |
| 15 | GAGGAGGGCAGCAAACGG | AA | AACCGTGCTCACACACATGCCACGG | ACGAAGGTGCCATATAATCCACTCC | TA | GAAGAGTCTTCCTTTACG |
| 16 | GAGGAGGGCAGCAAACGG | AA | ATCTCCGTACATTTCCAGCACCAGG | ATTGACGGTGGTCCAGATCTTCACC | TA | GAAGAGTCTTCCTTTACG |
| 17 | GAGGAGGGCAGCAAACGG | AA | TCCGGATTGGTCCAGCCACCCAGCT | GCATAGTCCTTGAAAAGCGGAATTA | TA | GAAGAGTCTTCCTTTACG |
| 18 | GAGGAGGGCAGCAAACGG | AA | AGATCGTCACCATCGGAGTGATGTT | GCAACTTCTGCGGCAATTCCCAGTG | TA | GAAGAGTCTTCCTTTACG |
| 19 | GAGGAGGGCAGCAAACGG | AA | ATAGTACTTAATGCCGGCTGTACTG | CCGCAGCAGTTCGTCGATCAGGTTG | TA | GAAGAGTCTTCCTTTACG |
| 20 | GAGGAGGGCAGCAAACGG | AA | CGTGGCCAGGACAGGGAGAAGCGAT | TGGTTCATATAGCCACCGGGCATAA | TA | GAAGAGTCTTCCTTTACG |
| 21 | GAGGAGGGCAGCAAACGG | AA | GCACATCGCGCTTCCACTGATGATA | TGCCCACGTGCAGCTCCTTGACCAT | TA | GAAGAGTCTTCCTTTACG |
| 22 | GAGGAGGGCAGCAAACGG | AA | ATCGACTATTTTCTCGGGATGAGTG | GTCCGCCGAGACATCGCCATTGGAT | TA | GAAGAGTCTTCCTTTACG |
| 23 | GAGGAGGGCAGCAAACGG | AA | TTCGATTTGGTATGAAGATGATCCA | CCCCTTGTCATCCGCGTTCCAGCCG | TA | GAAGAGTCTTCCTTTACG |
| 24 | GAGGAGGGCAGCAAACGG | AA | ACATGAGGCAGTCAGCACAAAAAGA | ACGGGTTTGGCTCACCGGACTGCCA | TA | GAAGAGTCTTCCTTTACG |

## CG10513

| Pair | Initiator | Spacer | Hybridzation | Hybridzation | Spacer | Initiator |
|---|---|---|---|---|---|---|
| 1 | GAGGAGGGCAGCAAACGG | AA | CCATTCAGTCAATTACGTCTAGTAG | TCAAATACTCGTAGTATATGATGGC | TA | GAAGAGTCTTCCTTTACG |
| 2 | GAGGAGGGCAGCAAACGG | AA | TGAGATTGTCCTGCAACCTCTTGTT | TCCGATCGAAGCGTGGCAGCTCGCG | TA | GAAGAGTCTTCCTTTACG |
| 3 | GAGGAGGGCAGCAAACGG | AA | TCGTTCGTCATCCTTCATCAAAGCA | GTACAATACCTTCCGGAAGTTACGG | TA | GAAGAGTCTTCCTTTACG |
| 4 | GAGGAGGGCAGCAAACGG | AA | GTCAATATGGCCTGGCAGACCAGAG | AAGTCGGCATCGGCGTTTTGGTCAT | TA | GAAGAGTCTTCCTTTACG |
| 5 | GAGGAGGGCAGCAAACGG | AA | CGAGCTGCAGGACGAACTGCCTCAA | CTGTGACAGCAAAGAATCTGCCCCT | TA | GAAGAGTCTTCCTTTACG |
| 6 | GAGGAGGGCAGCAAACGG | AA | CTTAAGCGTTTCCACGAGCACCGTG | GGGAATGTAGCCACCAAAGTTGAGA | TA | GAAGAGTCTTCCTTTACG |
| 7 | GAGGAGGGCAGCAAACGG | AA | TCGTAGCGAATATCGACCTGCACCG | TAATACTGAAACAGGGCATCCTGCT | TA | GAAGAGTCTTCCTTTACG |
| 8 | GAGGAGGGCAGCAAACGG | AA | CCGGCGAGGACCAACTGCAGAATTG | TGTTGAAGAAGTAGTGCAGATCCAC | TA | GAAGAGTCTTCCTTTACG |
| 9 | GAGGAGGGCAGCAAACGG | AA | CTTATTTTCGCCGTAGCGCAACATC | GTCGATGAGAGTCATGTCCAGTGGC | TA | GAAGAGTCTTCCTTTACG |
| 10 | GAGGAGGGCAGCAAACGG | AA | AATGTATTAAAATCTCCCGGTTGGG | TTGTTCACCCAGTAGTCACCGTGCA | TA | GAAGAGTCTTCCTTTACG |
| 11 | GAGGAGGGCAGCAAACGG | AA | CACGCTCCTGCAGCTTCTTCAATTT | CGTAAACTCTTGTGGAGTACTCCAT | TA | GAAGAGTCTTCCTTTACG |
| 12 | GAGGAGGGCAGCAAACGG | AA | GCAAAGGCCTGGGTGTGGCGATTGA | ACGCCAACCGTATTCACAAAGAAGG | TA | GAAGAGTCTTCCTTTACG |
| 13 | GAGGAGGGCAGCAAACGG | AA | CGGGTTGACGTTCGTTCAGAACAGC | TGCCGTGATCGAACTTGGTCAGCAA | TA | GAAGAGTCTTCCTTTACG |
| 14 | GAGGAGGGCAGCAAACGG | AA | ACCAGACGATCGGCCAGGACATATT | AAGCGGGTGTGCTCCAGATCGAAAC | TA | GAAGAGTCTTCCTTTACG |
| 15 | GAGGAGGGCAGCAAACGG | AA | CCTCGTGCTCGTAGTCCACATGCAA | TCACCGCAAGGTCCTCAAAGATGAT | TA | GAAGAGTCTTCCTTTACG |
| 16 | GAGGAGGGCAGCAAACGG | AA | AGTCTTCTCGATGAGCGAGCTCAAT | CTTGGCGAAAACCTTTTCCGGCTGG | TA | GAAGAGTCTTCCTTTACG |
| 17 | GAGGAGGGCAGCAAACGG | AA | TCCGTGGTGCTAACCTGGTACTGGG | GGCAGGATCTTCTCGTACATACGCA | TA | GAAGAGTCTTCCTTTACG |
| 18 | GAGGAGGGCAGCAAACGG | AA | CATAGGTGGTCTTCACAATATAGTA | ATATTCCAGAGGCGAAGGCATCGTT | TA | GAAGAGTCTTCCTTTACG |
| 19 | GAGGAGGGCAGCAAACGG | AA | CAAGAACAAAATCCGCACGCGTGTC | AGTCTCGGGGCTTTGGCGCCACTT | TA | GAAGAGTCTTCCTTTACG |
| 20 | GAGGAGGGCAGCAAACGG | AA | GTGGCGGGCTTGATCACCAAATCGG | ACACTAGCATAATTGTCACCCTTGG | TA | GAAGAGTCTTCCTTTACG |
| 21 | GAGGAGGGCAGCAAACGG | AA | CCCGGAGCAGTCGCTCCAAGTAGGT | TCCTCAGTCCCGGATCATTCTTCAA | TA | GAAGAGTCTTCCTTTACG |
| 22 | GAGGAGGGCAGCAAACGG | AA | TGCCCTGCTCCACCATGATGATGCT | CCGGATGAAATTCGGTAGCGTCTTC | TA | GAAGAGTCTTCCTTTACG |

## CG14257

| Pair | Initiator | Spacer | Hybridzation | Hybridzation | Spacer | Initiator |
|---|---|---|---|---|---|---|
| 1 | GAGGAGGGCAGCAAACGG | AA | AATTAATGCCAACTGAAGTTTCTGG | ATACTTAAACGTAAGATGGGGAAGT | TA | GAAGAGTCTTCCTTTACG |
| 2 | GAGGAGGGCAGCAAACGG | AA | TAGGCAGGAACGGTTTGACTAACTC | GGGGAGTGACAGCAGATAGATTCGG | TA | GAAGAGTCTTCCTTTACG |
| 3 | GAGGAGGGCAGCAAACGG | AA | TGGGCGTGGTCCACAGCTTCCTTAT | TGGAGATGGACGGTGCTGTTAACGA | TA | GAAGAGTCTTCCTTTACG |
| 4 | GAGGAGGGCAGCAAACGG | AA | ATAGGCCGATGGCAGATAGTTGCTT | CCTCTACTGATGGTAGACATCTTGT | TA | GAAGAGTCTTCCTTTACG |
| 5 | GAGGAGGGCAGCAAACGG | AA | AGTTGATGTGCGCCGAGTTGATGAT | AGGAGTGACTAGTGGCGTAGGTGTG | TA | GAAGAGTCTTCCTTTACG |
| 6 | GAGGAGGGCAGCAAACGG | AA | ATGGTGTTCAGCACGGATTTCTTAA | CCGGTGATGGCCTTGACCGCCTGGA | TA | GAAGAGTCTTCCTTTACG |
| 7 | GAGGAGGGCAGCAAACGG | AA | CATGGACAACGATCTCGTGGGATTT | ACCAGGGATCGTAGTCCACAGAGTT | TA | GAAGAGTCTTCCTTTACG |
| 8 | GAGGAGGGCAGCAAACGG | AA | TGCACCTGGGACTCCTTTACACTCT | ACGTAGTCAAAGCTGCCGGACCGCT | TA | GAAGAGTCTTCCTTTACG |
| 9 | GAGGAGGGCAGCAAACGG | AA | CCAGGGAAAGGGACTTATCGCTAGC | CCGGTTCGTTGTCCAGATCTGTGTC | TA | GAAGAGTCTTCCTTTACG |
| 10 | GAGGAGGGCAGCAAACGG | AA | AAAGACTTAGGCTGAAGACCTTCAT | ACATCAGAGGGAGGGCGGCCACCAG | TA | GAAGAGTCTTCCTTTACG |
| 11 | GAGGAGGGCAGCAAACGG | AA | CTCAAATTGGGCCTTCGATTAGCTG | GTGCGCCTGTGTTTTTCGTGGGCTT | TA | GAAGAGTCTTCCTTTACG |
| 12 | GAGGAGGGCAGCAAACGG | AA | ACCGATGACGTCCTCGGGTTGTTTT | GGCAAATCGAGATCAAAATTCACTT | TA | GAAGAGTCTTCCTTTACG |
| 13 | GAGGAGGGCAGCAAACGG | AA | AACTTTCTTCACTCGGCCACTTGCA | TTTTTAACGCTGTCTCCTCTTCACA | TA | GAAGAGTCTTCCTTTACG |
| 14 | GAGGAGGGCAGCAAACGG | AA | ACTTCGATTTCGAACCGTTTCGCTT | GGCACTTTTGGCCGCCGCACTTTTT | TA | GAAGAGTCTTCCTTTACG |
| 15 | GAGGAGGGCAGCAAACGG | AA | AGCGGTTTGCAGCGCGCACACAAAA | GTTTGTTTTGTCATCGGCATTCGTC | TA | GAAGAGTCTTCCTTTACG |
| 16 | GAGGAGGGCAGCAAACGG | AA | AGTCTCCCGATCTGGGGATTTTGTA | CACTCTCCACTTTCTTTCACCGATT | TA | GAAGAGTCTTCCTTTACG |

## CG17108

| Pair | Initiator | Spacer | Hybridzation | Hybridzation | Spacer | Initiator |
|---|---|---|---|---|---|---|
| 1 | CCTCAACCTACCTCCAAC | AA | GTTTCGATGTAATACTTTAAGCCAC | GTTGAGTACAAAAACTATATTTGAT | AT | TCTCACCATATTCGCTTC |
| 2 | CCTCAACCTACCTCCAAC | AA | GTAATCCAGACTAGCCATGATAGCC | GAGAATTACAGGTATATTGGGCTTT | AT | TCTCACCATATTCGCTTC |
| 3 | CCTCAACCTACCTCCAAC | AA | ACCTAAGGAGACTGCAGGTCCAATT | GTAGCTACCACCACCGCCTGGTCCG | AT | TCTCACCATATTCGCTTC |
| 4 | CCTCAACCTACCTCCAAC | AA | CTCCATGGCCTTTGTGCTTGTGGCT | CGTATCCGCCTTTGAATCCGCCTCC | AT | TCTCACCATATTCGCTTC |
| 5 | CCTCAACCTACCTCCAAC | AA | ACCTCCACCAATAACTCCAACTGAA | TCCTCCGCCGTGACCTCCGCCGCCA | AT | TCTCACCATATTCGCTTC |
| 6 | CCTCAACCTACCTCCAAC | AA | CCGATTCCACCGCCTCCACCAGAAT | ATTCCTCCACCAAATCCAGGACCGC | AT | TCTCACCATATTCGCTTC |
| 7 | CCTCAACCTACCTCCAAC | AA | CCGATTCCGCTACCTCCACCAGAAT | ATTCCGCCACCAAATCCAGGGCCGC | AT | TCTCACCATATTCGCTTC |
| 8 | CCTCAACCTACCTCCAAC | AA | CTCCGGAATATCCACCTGATCCAAA | CACCTCCACCGCCGATTCCGCCACC | AT | TCTCACCATATTCGCTTC |
| 9 | CCTCAACCTACCTCCAAC | AA | TCCGAATCCGGGACCGCCACCGAAT | GATTCCTCCACCGGAATGACCACCT | AT | TCTCACCATATTCGCTTC |
| 10 | CCTCAACCTACCTCCAAC | AA | AGAGCCGAAACCGCCAGATCCAAGT | GCCGCCTCCAAGTCCTAAATCACCG | AT | TCTCACCATATTCGCTTC |
| 11 | CCTCAACCTACCTCCAAC | AA | CCTCCGCCGAGCTTACCGCCTCCAT | CCAGAACCGTGACCACCAGATCCAT | AT | TCTCACCATATTCGCTTC |
| 12 | CCTCAACCTACCTCCAAC | AA | TAACGCCACACAAACACTCAGCACT | TCCACCGCCGTAACCTCCGGCGGAA | AT | TCTCACCATATTCGCTTC |
| 13 | CCTCAACCTACCTCCAAC | AA | AGCTGACTAACTGGGCTCCCTGGAA | AACAGACGCATCGGAACCTTTGCCT | AT | TCTCACCATATTCGCTTC |

# CG17239

| Pair | Initiator | Spacer | Hybridzation | Hybridzation | Spacer | Initiator |
|---|---|---|---|---|---|---|
| 1 | GAGGAGGGCAGCAAACGG | AA | ATGGCTTAAATATGTTGTAAAATCC | GAAGTCCAGGAAAACAAGGATCTGC | TA | GAAGAGTCTTCCTTTACG |
| 2 | GAGGAGGGCAGCAAACGG | AA | TTCGATAGCGCCCAAAATCCAGGGC | ATTAATTTATTCAGATTTAGTTATT | TA | GAAGAGTCTTCCTTTACG |
| 3 | GAGGAGGGCAGCAAACGG | AA | GGATATTCGGGATGAGCGCACTCCT | AGCTCAGCCACGTTAGCGTAGACTC | TA | GAAGAGTCTTCCTTTACG |
| 4 | GAGGAGGGCAGCAAACGG | AA | TACCGGAGACCAGAGGACCTCCCGA | CAAAGGACACGATCCCAACGAGCTT | TA | GAAGAGTCTTCCTTTACG |
| 5 | GAGGAGGGCAGCAAACGG | AA | CCGCACAGATCATGTCCTTGGTGAT | CGGAGCAGGCATCCTTTCCTGGAGC | TA | GAAGAGTCTTCCTTTACG |
| 6 | GAGGAGGGCAGCAAACGG | AA | CTGGTCCACAATGTCCACCGAAGCG | TCTTCCGTACGACCTACGACACTGA | TA | GAAGAGTCTTCCTTTACG |
| 7 | GAGGAGGGCAGCAAACGG | AA | AAACCAATGGCTCCCCATCCAGAAA | AGTATGGACATGGGGTAATTCTTCT | TA | GAAGAGTCTTCCTTTACG |
| 8 | GAGGAGGGCAGCAAACGG | AA | TGACTGCACTTCCCAGCCGGAGCTT | GGGGAGTATCGGCCAAGGGAATGAC | TA | GAAGAGTCTTCCTTTACG |
| 9 | GAGGAGGGCAGCAAACGG | AA | GGACCAGGACTGGTCGTACTCCTCG | CTGAAGCCTCATCACGGCTATGTCG | TA | GAAGAGTCTTCCTTTACG |
| 10 | GAGGAGGGCAGCAAACGG | AA | TGACCACCGAAGAATGTGAACGATG | AACAGAACCCTGGATACCCTCACCA | TA | GAAGAGTCTTCCTTTACG |
| 11 | GAGGAGGGCAGCAAACGG | AA | TGGGCTGCCGTTATGACGATGTCTT | AACTCGGTCTCGCGGTCGGTGAGGC | TA | GAAGAGTCTTCCTTTACG |
| 12 | GAGGAGGGCAGCAAACGG | AA | GTCCACGCCTAAGAATGGAAGCCTG | TGTAGATGGCTGCACCGCAGTGAAA | TA | GAAGAGTCTTCCTTTACG |
| 13 | GAGGAGGGCAGCAAACGG | AA | GCCTCCCACGATTCGTTCCGGAATC | GGGAACTGATAGAATGGTTATCAAG | TA | GAAGAGTCTTCCTTTACG |
| 14 | GAGGAGGGCAGCAAACGG | AA | GCCAGGAAAATCCACTGCACAAACA | TTAGAGGAAACCACGGTGACGCTGA | TA | GAAGAGTCTTCCTTTACG |

# CG23700

| Pair | Initiator | Spacer | Hybridzation | Hybridzation | Spacer | Initiator |
|---|---|---|---|---|---|---|
| 1 | CCTCAACCTACCTCCAAC | AA | ACAATAGAGACAGTTACAGAGCGAC | CATTATAAGATACGAGTAATCTCAA | AT | TCTCACCATATTCGCTTC |
| 2 | CCTCAACCTACCTCCAAC | AA | GCGGGTACAATAGATTTACAAATGG | GGATTGTGGTTCACGATTATTTATT | AT | TCTCACCATATTCGCTTC |
| 3 | CCTCAACCTACCTCCAAC | AA | ATTTCCCTCGAATTTCTGCAATGGC | CTGGAAAACCTTTTGCTAAATACAA | AT | TCTCACCATATTCGCTTC |
| 4 | CCTCAACCTACCTCCAAC | AA | GGCAAACATAATGGCGAACTACGAG | CTTTTCTTACATTTATCTCCGATTT | AT | TCTCACCATATTCGCTTC |
| 5 | CCTCAACCTACCTCCAAC | AA | ATTTTTGCCATCATCGTCCAAATTC | CTTTTACATCCTTGCATTCTCATCA | AT | TCTCACCATATTCGCTTC |
| 6 | CCTCAACCTACCTCCAAC | AA | TGCTCAGGTCAAATTAGAGGAAAGG | TTTTTCCTTCGTCGTCCAGCGACAT | AT | TCTCACCATATTCGCTTC |
| 7 | CCTCAACCTACCTCCAAC | AA | ACCCATTTCATATTGAAAATTGGGG | TTTAGTTACGGGTCTCTTTAATTCT | AT | TCTCACCATATTCGCTTC |
| 8 | CCTCAACCTACCTCCAAC | AA | AAGACGCAAACTCGTTTCCGGTTCG | TTTCCTTCATTTTCCTGGAGTGCTA | AT | TCTCACCATATTCGCTTC |
| 9 | CCTCAACCTACCTCCAAC | AA | TGTACACTCTATGAGAAACAACAAC | GACGAGGACATATATCCTATGTTTA | AT | TCTCACCATATTCGCTTC |
| 10 | CCTCAACCTACCTCCAAC | AA | GACTTGAGAGTGCGGCGATCTCTTC | GAGACTTATATAAAAGAAAACTTCA | AT | TCTCACCATATTCGCTTC |
| 11 | CCTCAACCTACCTCCAAC | AA | TCATCGAATTGCACTACGTTACATG | CGAGACAGTTTTTCCAAATATTCGA | AT | TCTCACCATATTCGCTTC |
| 12 | CCTCAACCTACCTCCAAC | AA | ATTGGATTTCGGGCCTGGTCTACTC | GTAGTATTATTATTATTAATTGGGG | AT | TCTCACCATATTCGCTTC |
| 13 | CCTCAACCTACCTCCAAC | AA | CAGCCCAGTTCAAAAATAGTTGAAT | GCTGACCTGCTGGGCTGGGCAAGCA | AT | TCTCACCATATTCGCTTC |
| 14 | CCTCAACCTACCTCCAAC | AA | TAAATCGTTTTGCTCTTAGCCTAGA | CACATCGTTTTATTGCTGGTTTGGC | AT | TCTCACCATATTCGCTTC |
| 15 | CCTCAACCTACCTCCAAC | AA | ATCAATCAGGGTTAAATGGTAACTG | TCAATTTTGATTGCGCTGGTTGATT | AT | TCTCACCATATTCGCTTC |
| 16 | CCTCAACCTACCTCCAAC | AA | CACCTGAGGATTTCCACTTTCTAAT | AATGGAAAAACGAGCTAGTTCGAAA | AT | TCTCACCATATTCGCTTC |
| 17 | CCTCAACCTACCTCCAAC | AA | TTTCTTTCATGCGGTGGTTCACGAC | ATATTAGTTAGTTTTCATTTTGGTG | AT | TCTCACCATATTCGCTTC |
| 18 | CCTCAACCTACCTCCAAC | AA | CTTGAAATCGATTGATAATGGAACG | AACTGATTACATTGAGTTGCGTGTT | AT | TCTCACCATATTCGCTTC |
| 19 | CCTCAACCTACCTCCAAC | AA | TTTCGAGGATGGGTATCGAGTGTTG | TTTTTGACCACTTGGGAATTATAAT | AT | TCTCACCATATTCGCTTC |
| 20 | CCTCAACCTACCTCCAAC | AA | AACACTCCAATTAAATATTATTTGC | TTGGTTTCCTGTTGTCCCTTCGTCC | AT | TCTCACCATATTCGCTTC |
| 21 | CCTCAACCTACCTCCAAC | AA | TTTTTAAGTCGTAATTGTGGGCGAT | TGTCCTTGCATAGGGAGGATGTGTC | AT | TCTCACCATATTCGCTTC |
| 22 | CCTCAACCTACCTCCAAC | AA | ACATGGCACAGGTGTTAACCAAAAT | GCGACAGTTGTGGGCGTGGCACATG | AT | TCTCACCATATTCGCTTC |
| 23 | CCTCAACCTACCTCCAAC | AA | GCTTTATTTGTCCGCCTTTTCGCGT | TTTTTGGTTCTCTTTCACGTTCGTT | AT | TCTCACCATATTCGCTTC |
| 24 | CCTCAACCTACCTCCAAC | AA | TCTTGCCATCTCGCTCTTTGTTTAA | CCGCCCATTCTTTTCATCTTAAATT | AT | TCTCACCATATTCGCTTC |
| 25 | CCTCAACCTACCTCCAAC | AA | TGTTTGTTTGCTTTATGCGGCAGCC | TTTTTGTTTCATGCGGTTTTTGTGG | AT | TCTCACCATATTCGCTTC |
| 26 | CCTCAACCTACCTCCAAC | AA | AAATGTTAATAAAGGTCCAGTTGCA | TTTTTCCTTTGTCGGGTGGAAAGCA | AT | TCTCACCATATTCGCTTC |
| 27 | CCTCAACCTACCTCCAAC | AA | TGGTTGTTTGTATTGCCCAGTGTCC | CGAGTTGGTTCGGAGTTTTCAGGCA | AT | TCTCACCATATTCGCTTC |
| 28 | CCTCAACCTACCTCCAAC | AA | AATCCCCATCCAACCATCCAGCCAT | CGAATTCATCCGTCCATCTGTCGAA | AT | TCTCACCATATTCGCTTC |
| 29 | CCTCAACCTACCTCCAAC | AA | CTTATGTCTGCACGGGCACATTGCT | CTTGCTATCGTGCGTGTCCTTCAGC | AT | TCTCACCATATTCGCTTC |
| 30 | CCTCAACCTACCTCCAAC | AA | CACGCTGCCGCGGTGATCGGTTAAT | ATCCATGATGTCCAGCTGCAAGTTG | AT | TCTCACCATATTCGCTTC |
| 31 | CCTCAACCTACCTCCAAC | AA | GCGGCGATTGCGTCCGAAAAGCGTT | ACTGAAGGATACACCGGAAATGGAC | AT | TCTCACCATATTCGCTTC |
| 32 | CCTCAACCTACCTCCAAC | AA | GAATGCTTGCGCCGCTGTTGCATAT | CCCTCGGTAATTTGAAGCGATGGCA | AT | TCTCACCATATTCGCTTC |
| 33 | CCTCAACCTACCTCCAAC | AA | TTACTTCACCTTTGACTGCGGTGTG | TTTTTCTGGCTGCTTCAATTACTTT | AT | TCTCACCATATTCGCTTC |
| 34 | CCTCAACCTACCTCCAAC | AA | TCCTCCTCCGAATCGGAACGCGAAT | TCTGTGTCTCAAGCTTTCTCCGCCA | AT | TCTCACCATATTCGCTTC |
| 35 | CCTCAACCTACCTCCAAC | AA | TCACTTCGGCTCGGTCGTTTGTATT | ATTTAAATATTTGCCACACACACTC | AT | TCTCACCATATTCGCTTC |
| 36 | CCTCAACCTACCTCCAAC | AA | ATCTTTCGAAAATCGAAAACCGGGC | AATTATGTATGTTCAGCACTTTCGA | AT | TCTCACCATATTCGCTTC |
| 37 | CCTCAACCTACCTCCAAC | AA | GACCAGTGACCAGTGACTTGCGACT | GGCAGTAACTGCGTGACTGTGAACT | AT | TCTCACCATATTCGCTTC |

# CG42780

| Pair | Initiator | Spacer | Hybridzation | Hybridzation | Spacer | Initiator |
|---|---|---|---|---|---|---|
| 1 | CCTCGTAAATCCTCATCA | AA | GATACTTTTGGTCAATTCCCTTTGC | ATCATTATATTTTAGCACATAGCGG | AA | ATCATCCAGTAAACCGCC |
| 2 | CCTCGTAAATCCTCATCA | AA | ATAAACACGTTCGCCAATAACATTT | CTCTATGCCTGCCTTGGCGCATTCT | AA | ATCATCCAGTAAACCGCC |
| 3 | CCTCGTAAATCCTCATCA | AA | CGCCGGATTTCACCCAGATTATATG | TAAGCGTAGTTACAGGCCAGATTGT | AA | ATCATCCAGTAAACCGCC |
| 4 | CCTCGTAAATCCTCATCA | AA | TCTCATTGATCAAAACGGTCAAGTG | CTAAGCCACAACCAACAGCATTACT | AA | ATCATCCAGTAAACCGCC |
| 5 | CCTCGTAAATCCTCATCA | AA | TTAGGTGTCGTTTTCTTTAAGTCCT | AAAAAGTAAGTGACAAACTCTGGTG | AA | ATCATCCAGTAAACCGCC |
| 6 | CCTCGTAAATCCTCATCA | AA | ACTTTTGAACTGCCATCTCAAAGAG | CGGTAATATCCTTTTCCTCGCCTGC | AA | ATCATCCAGTAAACCGCC |
| 7 | CCTCGTAAATCCTCATCA | AA | TCCTAGCATGAGAAAATCCCATGGC | ACGTCATATTCATTTTAGTTTTCGT | AA | ATCATCCAGTAAACCGCC |
| 8 | CCTCGTAAATCCTCATCA | AA | TTCCGTATTGTGACACTCATCATGT | TAGATTCTGTCCAGATAGCCGAAAC | AA | ATCATCCAGTAAACCGCC |
| 9 | CCTCGTAAATCCTCATCA | AA | GCCAGCTTTTCCAGGTCCTTATTCC | ATAAGACAGGTTTTGGCATTTAAGA | AA | ATCATCCAGTAAACCGCC |
| 10 | CCTCGTAAATCCTCATCA | AA | ACTTAATTTTTGCTTTGCCACTAGC | CCATTTTGGCCATTTTACAGGCGGT | AA | ATCATCCAGTAAACCGCC |
| 11 | CCTCGTAAATCCTCATCA | AA | GATCAGAGCATTTTTATCCCCAAGA | TTTTTGTCTCACCAAGTTGTGGGCC | AA | ATCATCCAGTAAACCGCC |
| 12 | CCTCGTAAATCCTCATCA | AA | CAAGAGCTACCAAAGGATCCGTTTA | AGTTTAATCACGGTTGCATCTTTGG | AA | ATCATCCAGTAAACCGCC |
| 13 | CCTCGTAAATCCTCATCA | AA | TTTGTCGCTAAGTTTTCGAATGATC | CTGCTGCTCACATAACACATATTGA | AA | ATCATCCAGTAAACCGCC |
| 14 | CCTCGTAAATCCTCATCA | AA | GGTACAGAGATCCTGCCTGCAAAAA | TACACACGCTATGTGGGTGGTTCCT | AA | ATCATCCAGTAAACCGCC |
| 15 | CCTCGTAAATCCTCATCA | AA | GTACGAATATATTGAACGTAAAGAC | CTGCCAAACTATGGAGAACTAGTGA | AA | ATCATCCAGTAAACCGCC |

# CG43112

| Pair | Initiator | Spacer | Hybridzation | Hybridzation | Spacer | Initiator |
|---|---|---|---|---|---|---|
| 1 | CCTCGTAAATCCTCATCA | AA | TTAGAAAGTGCCCACTTTCATTTCC | ATATTTCATCCCGACAAATTTCTAC | AA | ATCATCCAGTAAACCGCC |
| 2 | CCTCGTAAATCCTCATCA | AA | GGTTTTATAGGGTTTCACCTGGTGG | TCAGTACTTTCCATTAACGACGTCA | AA | ATCATCCAGTAAACCGCC |
| 3 | CCTCGTAAATCCTCATCA | AA | CGCCCAAATAATACCTCCGTTTGAC | TACTCCTTTAGAGGGTATAATATAT | AA | ATCATCCAGTAAACCGCC |
| 4 | CCTCGTAAATCCTCATCA | AA | ATTTTAATGTTCTTTATTCACGCCC | GGCATAGTAAATGAGAACTCCGGTT | AA | ATCATCCAGTAAACCGCC |
| 5 | CCTCGTAAATCCTCATCA | AA | TGACCAGAGGCGTGGTCAGCGATGT | GTCACATGTCTCGCCAAGTGATTGC | AA | ATCATCCAGTAAACCGCC |
| 6 | CCTCGTAAATCCTCATCA | AA | CAAAATTCAAAGAAATCCCATTCTG | AATTGCTACAGGCCAAACACTTGCC | AA | ATCATCCAGTAAACCGCC |
| 7 | CCTCGTAAATCCTCATCA | AA | CATCGGGATAGTCCATGAATTCGAT | ATTCATTGCTATAGGCGGTAGCTCT | AA | ATCATCCAGTAAACCGCC |
| 8 | CCTCGTAAATCCTCATCA | AA | GGCGTAGGCAGAGACCAGACAAAGG | GTGATCGTTACAGTCGGTATGATCC | AA | ATCATCCAGTAAACCGCC |
| 9 | CCTCGTAAATCCTCATCA | AA | ATCTCGATTTCCGTTCAATTCGATA | CTGATGAGGACCTGGAGTGTTGGGC | AA | ATCATCCAGTAAACCGCC |
| 10 | CCTCGTAAATCCTCATCA | AA | TGACAATTTCGACTCAACGGCAGCT | TGTTAGTTCAAACTTAAGTGCAACT | AA | ATCATCCAGTAAACCGCC |

### cyp313a3

| Pair | Initiator | Spacer | Hybridzation | Hybridzation | Spacer | Initiator |
|---|---|---|---|---|---|---|
| 1 | CCTCAACCTACCTCCAAC | AA | AGTCCAGGTGACTGAGCTAGTTCCA | TTCTTTTAAGTTCGCCGATGAAATT | AT | TCTCACCATATTCGCTTC |
| 2 | CCTCAACCTACCTCCAAC | AA | CGTATCGAAAAGATGTGCTTACTTT | CAATGTTGTCAACGAACTCGAGATC | AT | TCTCACCATATTCGCTTC |
| 3 | CCTCAACCTACCTCCAAC | AA | GATGACATCAACCCATATTTCCAAC | CTAAGTATCTTAGACAAAGCAAGCT | AT | TCTCACCATATTCGCTTC |
| 4 | CCTCAACCTACCTCCAAC | AA | GAATGTATGCGTAAGGATGCCTATC | TACAATTCCGCCTTCCCTTCGAAAA | AT | TCTCACCATATTCGCTTC |
| 5 | CCTCAACCTACCTCCAAC | AA | GTTGAACGACGATGGATCGGTACCC | CACATTGTCGGGCAGAAAGTGATCT | AT | TCTCACCATATTCGCTTC |
| 6 | CCTCAACCTACCTCCAAC | AA | ATATCGATACCAATACCAACTCCTT | TGATCGCGGTTGCGATGGGTAGCAA | AT | TCTCACCATATTCGCTTC |
| 7 | CCTCAACCTACCTCCAAC | AA | AATCCCGTATTGTTTCCCTCGGTGT | GAATGACCACTCCAGAGGATAGACG | AT | TCTCACCATATTCGCTTC |
| 8 | CCTCAACCTACCTCCAAC | AA | GGTCACCTCAAAGTCACCGGCCACG | AAAAACCATCCTTTGCAAATCGTCG | AT | TCTCACCATATTCGCTTC |
| 9 | CCTCAACCTACCTCCAAC | AA | ATACACCGTGTCACCAGTTGTCTCA | GAACATGGCCAGCAGGATGAGTGCG | AT | TCTCACCATATTCGCTTC |
| 10 | CCTCAACCTACCTCCAAC | AA | TTGCACCTCTTCCCGGGACATTTCT | AGCGACAACGAAGGAACAGCACTCC | AT | TCTCACCATATTCGCTTC |
| 11 | CCTCAACCTACCTCCAAC | AA | ATGACAGAGGTAATTTCAGGCTGTG | TTCCGATGGAGTTCGATGGCCTTAT | AT | TCTCACCATATTCGCTTC |
| 12 | CCTCAACCTACCTCCAAC | AA | CATTTTGTTAACATTGGACTTGGCC | TAACTTTTTGTCAACAATCGTGCCT | AT | TCTCACCATATTCGCTTC |
| 13 | CCTCAACCTACCTCCAAC | AA | GTTGAAAATATCTTGTTGTGAGTGA | GCTTTCTGCGTTTCAAATCCGCCCA | AT | TCTCACCATATTCGCTTC |
| 14 | CCTCAACCTACCTCCAAC | AA | CTGAATCGTTCTTGAAGGACGTATC | TCATAAACGTCTCGTAACTTTTCAA | AT | TCTCACCATATTCGCTTC |
| 15 | CCTCAACCTACCTCCAAC | AA | AGTGGCTATTCTAAAAGACCATCTC | CTTTACATCAGTTCCCACTGTGGTT | AT | TCTCACCATATTCGCTTC |
| 16 | CCTCAACCTACCTCCAAC | AA | CAGGAAGGCAACTAAAGTTTTCGCC | TTTTTCACCCTGACCGACAAGTGAG | AT | TCTCACCATATTCGCTTC |
| 17 | CCTCAACCTACCTCCAAC | AA | AAAACATTTTGCTTGAATGCCGGAT | GAATTGAAGATGGGAAGAAAACTCA | AT | TCTCACCATATTCGCTTC |
| 18 | CCTCAACCTACCTCCAAC | AA | ATGCCTCTAATGATAATAATCCATC | AGTTCTTGCGACGATCAACCCATTT | AT | TCTCACCATATTCGCTTC |
| 19 | CCTCAACCTACCTCCAAC | AA | TATCGAGCTCCTATTTAGGCACTCT | TGTACAGCTTTTAACCGGCTTGGAA | AT | TCTCACCATATTCGCTTC |
| 20 | CCTCAACCTACCTCCAAC | AA | GGATCCCGAGTGATTACAAATGGCG | GACAAAAAGATTTCCTCGGCAATCT | AT | TCTCACCATATTCGCTTC |
| 21 | CCTCAACCTACCTCCAAC | AA | CGTAAATATCCATATACTTGGTTCG | GTCCCAACCACACCAAACATGTGGA | AT | TCTCACCATATTCGCTTC |
| 22 | CCTCAACCTACCTCCAAC | AA | ATACTCAAAGGCTATGCCCAGAATT | GCTCATTTTACGTTTATAGGTTATC | AT | TCTCACCATATTCGCTTC |
| 23 | CCTCAACCTACCTCCAAC | AA | TGCATCATATAGAGCCGGCGACGAC | AATCCCATTGGTCCTGGTATCTTAA | AT | TCTCACCATATTCGCTTC |
| 24 | CCTCAACCTACCTCCAAC | AA | CCACGGCTAAGAGCAGTTGAAACGT | ATAGGAAGTATATCCAGAAACACAC | AT | TCTCACCATATTCGCTTC |

### EbpIII

| Pair | Initiator | Spacer | Hybridzation | Hybridzation | Spacer | Initiator |
|---|---|---|---|---|---|---|
| 1 | CCTCGTAAATCCTCATCA | AA | TGATGATCAGATATAAACGTAGACC | TTCCGAGCACGATCGTAGAAATAAT | AA | ATCATCCAGTAAACCGCC |
| 2 | CCTCGTAAATCCTCATCA | AA | CGGAACCGGGATCCGCCTTACACCT | TTAACAGTGCATATAACCAAGTGTT | AA | ATCATCCAGTAAACCGCC |
| 3 | CCTCGTAAATCCTCATCA | AA | TTGGCCTGCAGCTGCTTCCACTCCT | TTGATGTAGATCTCCTCGGGATCGT | AA | ATCATCCAGTAAACCGCC |
| 4 | CCTCGTAAATCCTCATCA | AA | CCTTGTCGGTGTTCTGGCGCTGCTT | GCTTGTTTTCGATGATGTAGCGAAT | AA | ATCATCCAGTAAACCGCC |
| 5 | CCTCGTAAATCCTCATCA | AA | AGGCACTTGAAGTAGTTTCCGAAGA | TCCGGGGTGCACTTTCCGTTGTCCA | AA | ATCATCCAGTAAACCGCC |
| 6 | CCTCGTAAATCCTCATCA | AA | CGATGTTGTCGTACTTGGTGGTGTA | GGTCCGACTTCAGGATCTCGTCGAC | AA | ATCATCCAGTAAACCGCC |
| 7 | CCTCGTAAATCCTCATCA | AA | CAAGCGCAAGTATCATCTTCATCTT | CGGCGACCAGCACCAGTCCAAGGAC | AA | ATCATCCAGTAAACCGCC |
| 8 | CCTCGTAAATCCTCATCA | AA | GACTCGGAGCGAATAGATGTGAAGC | GTTTTAATGGGGCTCGACTTGTGGG | AA | ATCATCCAGTAAACCGCC |

### FASN2

| Pair | Initiator | Spacer | Hybridzation | Hybridzation | Spacer | Initiator |
|---|---|---|---|---|---|---|
| 1 | CCTCAACCTACCTCCAAC | AA | TACGTGTATTGGTAGGATTAGGGTA | ATTCTTAAACAATCAATGCGCGATT | AT | TCTCACCATATTCGCTTC |
| 2 | CCTCAACCTACCTCCAAC | AA | AAATCTCCTTGAGGCCATAGTCCTG | CCACATGCATGTCGATCTGGGCACT | AT | TCTCACCATATTCGCTTC |
| 3 | CCTCAACCTACCTCCAAC | AA | GCGTGTAACCAGCCAGAGTGTCTGC | CCTTCTTGATCACTTCTCTGGTTTG | AT | TCTCACCATATTCGCTTC |
| 4 | CCTCAACCTACCTCCAAC | AA | CTGGGGTAATACTTCCTTGGCGAAT | CTTTGACCGAAGGCGCACCATCACC | AT | TCTCACCATATTCGCTTC |
| 5 | CCTCAACCTACCTCCAAC | AA | TGGTCTTATCGGAGACACTCTTTAC | GCGAGTCCATGCCAAGATCGAAGAG | AT | TCTCACCATATTCGCTTC |
| 6 | CCTCAACCTACCTCCAAC | AA | GCTGTGCCCGGATAACCATATTCCT | GTGTCCCCAATTGCTCCCCATTGAA | AT | TCTCACCATATTCGCTTC |
| 7 | CCTCAACCTACCTCCAAC | AA | GCCAGTAGCTTCGGTGCACTTACCT | CGGGAGATCTTGTCCAAGTGTATGG | AT | TCTCACCATATTCGCTTC |
| 8 | CCTCAACCTACCTCCAAC | AA | ATCTGAACCTGAACTCCACGGTCGT | ACAGCGGTGGTCACATCGTTGGTAT | AT | TCTCACCATATTCGCTTC |
| 9 | CCTCAACCTACCTCCAAC | AA | TCTCCTGCTCTGCAAAGACTGTGGT | ACGCCATGAAACGGAACGCCTTCTC | AT | TCTCACCATATTCGCTTC |
| 10 | CCTCAACCTACCTCCAAC | AA | CTTGCAGCTTCTCCTCGGAGAGAGA | CATTCAAAGCCAGACAGCGAATGGA | AT | TCTCACCATATTCGCTTC |
| 11 | CCTCAACCTACCTCCAAC | AA | ATTTGGCCCCTGACCACCAAAGCGT | TGGATGAGGATCCTTTCACCCTCCT | AT | TCTCACCATATTCGCTTC |
| 12 | CCTCAACCTACCTCCAAC | AA | GCCAACATCACATCCCGGAAATTGA | AGAGCATCCACTCCCAGCTTACCGG | AT | TCTCACCATATTCGCTTC |
| 13 | CCTCAACCTACCTCCAAC | AA | TGGGGCACACCTTCATCCAGAATAA | GCGTAGAAGGGATCGTCCAGGGAGA | AT | TCTCACCATATTCGCTTC |
| 14 | CCTCAACCTACCTCCAAC | AA | GTTTATTGGTGAGGTGAAGTGTCAG | TTTTTAGCTCGTTGACCCACTTGAA | AT | TCTCACCATATTCGCTTC |
| 15 | CCTCAACCTACCTCCAAC | AA | CCTCCAGGAGCAGGAATCCCGAATC | CTATAATGTTATACCTGCTGGCTGC | AT | TCTCACCATATTCGCTTC |
| 16 | CCTCAACCTACCTCCAAC | AA | TGAGCTGCTCACCCAGTTCGGATTC | TGACTATCACCCGAATACCGTGCTC | AT | TCTCACCATATTCGCTTC |
| 17 | CCTCAACCTACCTCCAAC | AA | GCTGGGAATGAGGCCAGAATCTATA | TTTCCTGCTCCTCCCGCTGACCCAA | AT | TCTCACCATATTCGCTTC |
| 18 | CCTCAACCTACCTCCAAC | AA | TGCTTGAGAGGATCTATCAGTGCCC | TGTTGGTGCTTCTTTGCCAGCTCCA | AT | TCTCACCATATTCGCTTC |
| 19 | CCTCAACCTACCTCCAAC | AA | CCCTGGAAAACACCAGTGTAGTCGT | ACCGCCGAAATATCCGACTCCAAAA | AT | TCTCACCATATTCGCTTC |
| 20 | CCTCAACCTACCTCCAAC | AA | TGCCTTCACAGATTTCAAAGGAGCT | GGATCTTGCCCGATGTCACCAGGCT | AT | TCTCACCATATTCGCTTC |
| 21 | CCTCAACCTACCTCCAAC | AA | TCTGTGTGTGATCCCAGCCCACTAA | CGCCCTTGAACTTGGGGTAGTTCCA | AT | TCTCACCATATTCGCTTC |
| 22 | CCTCAACCTACCTCCAAC | AA | AGCTAATCTGCGGACTTAAGGATCG | CATGACCTCGCTGCATCAGACTCAC | AT | TCTCACCATATTCGCTTC |
| 23 | CCTCAACCTACCTCCAAC | AA | CCAGCCGATGCCAGGTGGTTTTTCT | GGGGATATGAGGTTGTTTATAAAGT | AT | TCTCACCATATTCGCTTC |
| 24 | CCTCAACCTACCTCCAAC | AA | GCATGGGAGCCGCTCCTTCAATGTA | TGATCAGTCGCTCCAGACTTCTACG | AT | TCTCACCATATTCGCTTC |
| 25 | CCTCAACCTACCTCCAAC | AA | CAATTGTCATCACTGTTGTGACACA | ATGGATGACGGTGGCCCGGAGACCG | AT | TCTCACCATATTCGCTTC |
| 26 | CCTCAACCTACCTCCAAC | AA | TATGTCCAGCACCCGGAGAAGATCT | CGAATGGCCAATGATTCCATCTGGC | AT | TCTCACCATATTCGCTTC |
| 27 | CCTCAACCTACCTCCAAC | AA | CTATAAGATCAAAGTCCACCTGGGC | TTCGCTCCGTGGAACGGGTAAGCAC | AT | TCTCACCATATTCGCTTC |
| 28 | CCTCAACCTACCTCCAAC | AA | ACGGATGCAGGGGAATAGGATGTGA | TGGTACCGAGAACAGCGTATCCCCT | AT | TCTCACCATATTCGCTTC |
| 29 | CCTCAACCTACCTCCAAC | AA | TGCGACCGGAACAGGTAACTAGCCT | TCAGCAACTGATTCACTGCATCCTC | AT | TCTCACCATATTCGCTTC |
| 30 | CCTCAACCTACCTCCAAC | AA | GGAAGATTGCGATCCACCACCTTAA | TTCAGACCCACAATGCCACCCTGCC | AT | TCTCACCATATTCGCTTC |
| 31 | CCTCAACCTACCTCCAAC | AA | AATCCGTCCGTATTGGTCCGAACAT | TCTGGGAAGGTTACGCCCTGCTCCT | AT | TCTCACCATATTCGCTTC |
| 32 | CCTCAACCTACCTCCAAC | AA | CCTTGAAGTCGAACGTGTAGGAGAT | ATGCGGTGTCCACGATAAAGCTGGG | AT | TCTCACCATATTCGCTTC |
| 33 | CCTCAACCTACCTCCAAC | AA | ACGCTAAAGAACTTGTCGTCGAACT | TCCATCAGCTCCGCCTGCTTCTGGT | AT | TCTCACCATATTCGCTTC |
| 34 | CCTCAACCTACCTCCAAC | AA | CCGTTGAGCAGGTTGTACTTGAACT | CGCGGCTCATCTGTGACCATGTCGA | AT | TCTCACCATATTCGCTTC |
| 35 | CCTCAACCTACCTCCAAC | AA | TTTTGGACGGCACCTTATTCACTTG | GTATGCCACTAATTCGGATCGTATC | AT | TCTCACCATATTCGCTTC |

### Jhbp13

| Pair | Initiator | Spacer | Hybridzation | Hybridzation | Spacer | Initiator |
|---|---|---|---|---|---|---|
| 1 | CCTCGTAAATCCTCATCA | AA | ACCGCAAATTGAGCTACTCCTTGAG | TTAACCATCATCATTCTATTAAATC | AA | ATCATCCAGTAAACCGCC |
| 2 | CCTCGTAAATCCTCATCA | AA | CTCGTTGAACATTCGCAACATAAGG | CTGGTCGATGGGAACTAGTTCGAAT | AA | ATCATCCAGTAAACCGCC |
| 3 | CCTCGTAAATCCTCATCA | AA | ATTATTCCGTAAATATCGCGCCAAT | TGCCAAAACTGTCGAGTCTCAGGCA | AA | ATCATCCAGTAAACCGCC |
| 4 | CCTCGTAAATCCTCATCA | AA | CCAGTTCAGGATCTGCAAATATTCC | GATTAGCCACATTCAGAGCGATTTT | AA | ATCATCCAGTAAACCGCC |
| 5 | CCTCGTAAATCCTCATCA | AA | TGGCTTTGAGTCGATATTCTTGAGG | TGCCTTGATAACCAAGTCCTTGATC | AA | ATCATCCAGTAAACCGCC |
| 6 | CCTCGTAAATCCTCATCA | AA | ACCTCACCATCGGTCACCGTTATGG | AAAAACCGATGACCATCCTTCTCGT | AA | ATCATCCAGTAAACCGCC |
| 7 | CCTCGTAAATCCTCATCA | AA | CACCCTTGGGCTTCAGTTGCAGCTG | CCACATCCAAGAGCGTTACGTTGAA | AA | ATCATCCAGTAAACCGCC |
| 8 | CCTCGTAAATCCTCATCA | AA | CACAATGTTCAACTTGGGCATTTGG | CACCTGCATGTCGGCCTTATAGCTG | AA | ATCATCCAGTAAACCGCC |
| 9 | CCTCGTAAATCCTCATCA | AA | TTGAGTTTGACGGACACCTCTTTGG | ACCAGACGGAGTTTCACCTTGTCGC | AA | ATCATCCAGTAAACCGCC |
| 10 | CCTCGTAAATCCTCATCA | AA | CATTGCGAACGGTGATGCGACCATT | GATTGGAGGAGAAGCCATAGATCTT | AA | ATCATCCAGTAAACCGCC |
| 11 | CCTCGTAAATCCTCATCA | AA | TCGGTTGAGGTGCAAAGGTTCGTAG | AGTGCCACTAGAGTATTGGAAGCTG | AA | ATCATCCAGTAAACCGCC |
| 12 | CCTCGTAAATCCTCATCA | AA | TTGAGGCGCGGAGTCAATGTGTTAA | TCGATTCTCAACTCTGGATTACCGT | AA | ATCATCCAGTAAACCGCC |
| 13 | CCTCGTAAATCCTCATCA | AA | CCTCTTCAGTCGAGCATTTTGGCAA | GCTGCTTCACGCACTCACCCAGCTG | AA | ATCATCCAGTAAACCGCC |
| 14 | CCTCGTAAATCCTCATCA | AA | TGCACTGCATCCCAAGGCCAGCAGG | AGCAAAGTAGTTTTTATCCGTGGGA | AA | ATCATCCAGTAAACCGCC |
| 15 | CCTCGTAAATCCTCATCA | AA | TGGGACTCGAATTCGACTTTCAACC | CATTCTACTACTAAACTATATCCTT | AA | ATCATCCAGTAAACCGCC |

### obst-E

| Pair | Initiator | Spacer | Hybridzation | Hybridzation | Spacer | Initiator |
|---|---|---|---|---|---|---|
| 1 | GAGGAGGGCAGCAAACGG | AA | AATTATTGCACAGGAAGTGTACGGG | TAAGCAGTTACATATCTACAGCATA | TA | GAAGAGTCTTCCTTTACG |
| 2 | GAGGAGGGCAGCAAACGG | AA | GGCACCTTGTTGTAGAAGTCGCACA | TTCTCCTTGAGCAGGGCATAGCACT | TA | GAAGAGTCTTCCTTTACG |
| 3 | GAGGAGGGCAGCAAACGG | AA | CGCACACGAAGTACTTCTTGCAGGT | AGTTGTACAGGCGCGGATGTCCGTT | TA | GAAGAGTCTTCCTTTACG |
| 4 | GAGGAGGGCAGCAAACGG | AA | TCGCACTGGTAGGTCTCCTCGTTGA | TTGCAACTTTCCACTAGATCGGGCC | TA | GAAGAGTCTTCCTTTACG |
| 5 | GAGGAGGGCAGCAAACGG | AA | CCATCCTGACACTCTGTGTAAGAGT | TCCGGACACAGCTTCTCCACGGGAG | TA | GAAGAGTCTTCCTTTACG |
| 6 | GAGGAGGGCAGCAAACGG | AA | CGGCGAGCCAAGAGGTTTTTCCTTT | AAATCGTCCATTGGGCGTGGGGCAT | TA | GAAGAGTCTTCCTTTACG |
| 7 | GAGGAGGGCAGCAAACGG | AA | AGTTTGCCAAATTCTTTAACTACAG | ATTTCGACGCGCAAAATGTCAACAA | TA | GAAGAGTCTTCCTTTACG |
| 8 | GAGGAGGGCAGCAAACGG | AA | TGGCATGTGCATGGGAAAAATCACA | AGAAAACTCGAGCAACCTATCGCCA | TA | GAAGAGTCTTCCTTTACG |
| 9 | GAGGAGGGCAGCAAACGG | AA | AGACGAGCAGCCTTGATTTGGGCAC | GGATTGACTTAGTTCTTCCTGGCGT | TA | GAAGAGTCTTCCTTTACG |
| 10 | GAGGAGGGCAGCAAACGG | AA | TACAGATGAAGTACACCTGGCAATT | AGCCAATGCGACGCGGACGTCCCTC | TA | GAAGAGTCTTCCTTTACG |
| 11 | GAGGAGGGCAGCAAACGG | AA | GCTGGCATCACCCATGCGGTAGTAT | GGCGCAGTTCATGAACTGTCCACAA | TA | GAAGAGTCTTCCTTTACG |
| 12 | GAGGAGGGCAGCAAACGG | AA | GATCCGGACACGGGAGCAGTTCCAT | TTCTTGCACTCAATGTAGGCATCGC | TA | GAAGAGTCTTCCTTTACG |
| 13 | GAGGAGGGCAGCAAACGG | AA | GCAACGCACAATAGAGCACTGATTA | GCGGCGGCAGCCATTGAGCCAAACA | TA | GAAGAGTCTTCCTTTACG |
| 14 | GAGGAGGGCAGCAAACGG | AA | AAGTTAAACTAAGGAAGAGCTACTG | GTTTTCGAATGGCAATGTGATCACT | TA | GAAGAGTCTTCCTTTACG |

### red

| Pair | Initiator | Spacer | Hybridzation | Hybridzation | Spacer | Initiator |
|---|---|---|---|---|---|---|
| 1 | GAGGAGGGCAGCAAACGG | AA | TGGCTCGATATGAGTGCAGTTCGAT | AAAAACATTCTCACGTACAACATGA | TA | GAAGAGTCTTCCTTTACG |
| 2 | GAGGAGGGCAGCAAACGG | AA | ATGCTTGTTTAACAATTGAGGTTAC | ATTGGTGTATATATTGTTTTGTGGT | TA | GAAGAGTCTTCCTTTACG |
| 3 | GAGGAGGGCAGCAAACGG | AA | GTTGGCCAATTTCGTGTTTTAGCGG | AAAATAAATTAAAATTACACGTCAA | TA | GAAGAGTCTTCCTTTACG |
| 4 | GAGGAGGGCAGCAAACGG | AA | TCTGGCGACTATATACAATTAACAT | GAGCAGTCCAGTTGGATTGTTGTGT | TA | GAAGAGTCTTCCTTTACG |
| 5 | GAGGAGGGCAGCAAACGG | AA | CATACAAAAAGTAAACATAATGGGG | TATACACAGTGTGATTAAAGCTAGA | TA | GAAGAGTCTTCCTTTACG |
| 6 | GAGGAGGGCAGCAAACGG | AA | TTCCGTATCTGTTCGATGCGGTTCA | AGGAGGGCGGACAGGAGGACAGAAG | TA | GAAGAGTCTTCCTTTACG |
| 7 | GAGGAGGGCAGCAAACGG | AA | TTATACAGGGCTGACACAAGATCAA | CGGATCTTGCAAGCAAATTGGCGGC | TA | GAAGAGTCTTCCTTTACG |
| 8 | GAGGAGGGCAGCAAACGG | AA | ACTACAGGTTTACATAGCGCAGCGC | GGGTTTACAGAACTGCCTTGTCATC | TA | GAAGAGTCTTCCTTTACG |
| 9 | GAGGAGGGCAGCAAACGG | AA | TTTTATCGTATAGGTAAGGTATCGC | TTAGGTTTTAGCTCAGTTACATTAA | TA | GAAGAGTCTTCCTTTACG |
| 10 | GAGGAGGGCAGCAAACGG | AA | ACATTTAGAGTGCGGCATTTCCGTG | TTGTCGATAGATATCATTATTTATA | TA | GAAGAGTCTTCCTTTACG |
| 11 | GAGGAGGGCAGCAAACGG | AA | CCAATGCGCGGCAATTAAAAACGAA | TGTAACATGCTGACCAAAGTCATCT | TA | GAAGAGTCTTCCTTTACG |
| 12 | GAGGAGGGCAGCAAACGG | AA | TAGATTGCGTCCCGCAGACGGACGC | TAAAATTAACTCTCTGCGTAATTAG | TA | GAAGAGTCTTCCTTTACG |
| 13 | GAGGAGGGCAGCAAACGG | AA | TCCAGCCGCTTCAGCGAATTCTGGA | AACTCAAAGAAGTCGTCCTGCTGCT | TA | GAAGAGTCTTCCTTTACG |
| 14 | GAGGAGGGCAGCAAACGG | AA | ACGGACGGTCGTTGTTGTTGGTCTT | CCGTCGAGCTGTGGCGCTGATACTG | TA | GAAGAGTCTTCCTTTACG |
| 15 | GAGGAGGGCAGCAAACGG | AA | GATGCAGATGTCCGCATCACTCTGA | CTGCGGGAAAACGTCGGCGGTGGCA | TA | GAAGAGTCTTCCTTTACG |
| 16 | GAGGAGGGCAGCAAACGG | AA | TATTTGCGCGACTCAGAAATGGTGT | CTGACCAGGTCCTTAGAGCGCTCCA | TA | GAAGAGTCTTCCTTTACG |
| 17 | GAGGAGGGCAGCAAACGG | AA | ATTTCCTGTTCTCCTCCTCGGGCGT | CTATCTTGTTGAGGAAGTCGTTCAT | TA | GAAGAGTCTTCCTTTACG |
| 18 | GAGGAGGGCAGCAAACGG | AA | CCGTGGCCGTGCATCATCGATGGAT | CTCTTGCTCTGCTTGCCGCTGCAAC | TA | GAAGAGTCTTCCTTTACG |
| 19 | GAGGAGGGCAGCAAACGG | AA | TCCCACTGCTGCTGCTGCTACTGCT | TGCCACTTGTGTTGCAGCTGCTGCT | TA | GAAGAGTCTTCCTTTACG |
| 20 | GAGGAGGGCAGCAAACGG | AA | CTTGTTGCTCTGGCTCAGATTATTG | GCTGATGTCGTTGCTCTTCATGAAG | TA | GAAGAGTCTTCCTTTACG |
| 21 | GAGGAGGGCAGCAAACGG | AA | GCACCTGGGGATAGTACGGTGAGTT | CGTCGAAGGCGGCTATCGAATCGCC | TA | GAAGAGTCTTCCTTTACG |
| 22 | GAGGAGGGCAGCAAACGG | AA | CAGGAACAGGCTGTCCGAGGCGAAG | CTCCACTGGAACGAGGAGAAACTGA | TA | GAAGAGTCTTCCTTTACG |
| 23 | GAGGAGGGCAGCAAACGG | AA | CAGCCGTACTTGAGGGCGATGCCCT | TTGGCCCGTCTTATTTGCTCCGTCG | TA | GAAGAGTCTTCCTTTACG |
| 24 | GAGGAGGGCAGCAAACGG | AA | GGATGAGGGTCTCATTGTTCCGGAG | GCGTGTCCGTCTTCTCGACGATGTG | TA | GAAGAGTCTTCCTTTACG |
| 25 | GAGGAGGGCAGCAAACGG | AA | CCCGAATCGATTGCTTCTCGTTGAA | CGTAGCGCTTGAGGGTGCGCCCGGA | TA | GAAGAGTCTTCCTTTACG |
| 26 | GAGGAGGGCAGCAAACGG | AA | GATCTCTTGAAACCGAACGCCGCTC | GAATTCCGAGTCATCCATGTCGGGG | TA | GAAGAGTCTTCCTTTACG |
| 27 | GAGGAGGGCAGCAAACGG | AA | AACTGTTACAGTTACTCCGCCGCAG | TGCAACGGTAACTGCAACTGCAGCT | TA | GAAGAGTCTTCCTTTACG |
| 28 | GAGGAGGGCAGCAAACGG | AA | CCATGGCAGGCCTTTCAAAACAGAG | CCCAAACTCACACACGCACGCACAC | TA | GAAGAGTCTTCCTTTACG |
| 29 | GAGGAGGGCAGCAAACGG | AA | GCCTGCCAACGAATTGCGCTAATAT | GGTGGCGTCTCTACCGCCCACTGCT | TA | GAAGAGTCTTCCTTTACG |
| 30 | GAGGAGGGCAGCAAACGG | AA | ATTTGCGATTTTTCGATCCAATTGT | CACACGCGAGTGCTCTGTTTTATGG | TA | GAAGAGTCTTCCTTTACG |
| 31 | GAGGAGGGCAGCAAACGG | AA | GTTTCCAATCCTTGTAGGCCTTCAC | TTTCTGTTTACTACGTCTGCGAATG | TA | GAAGAGTCTTCCTTTACG |

## rk

| Pair | Initiator | Spacer | Hybridzation | Hybridzation | Spacer | Initiator |
|---|---|---|---|---|---|---|
| 1 | CCTCAACCTACCTCCAAC | AA | TTGGCCGAGTCGCCATCTTCCATGA | ATGCCTAGAAAACTGGCTACTCCCG | AT | TCTCACCATATTCGCTTC |
| 2 | CCTCAACCTACCTCCAAC | AA | TGGATGATCTTGAGCTGCAGAAGTT | CTGAGGATGGTGGACCATGTCGCCA | AT | TCTCACCATATTCGCTTC |
| 3 | CCTCAACCTACCTCCAAC | AA | TCAATCTCTGCCACATGCTCGGTGT | TTCGGTTCTCCCCGCAGCAGAATGT | AT | TCTCACCATATTCGCTTC |
| 4 | CCTCAACCTACCTCCAAC | AA | TGTTTGCACAACGTCACACAGTCCT | CCTCCCACGACGCGGGATTCTTCGA | AT | TCTCACCATATTCGCTTC |
| 5 | CCTCAACCTACCTCCAAC | AA | CTGCTCCAGCGAAATCAGTTGCAGG | CAGCACAAATACCGTAAAGATCTTG | AT | TCTCACCATATTCGCTTC |
| 6 | CCTCAACCTACCTCCAAC | AA | TTCGCGAATCATTCGTGTTCCATGC | AGACCAACAGAGCCATTCGCTTGGC | AT | TCTCACCATATTCGCTTC |
| 7 | CCTCAACCTACCTCCAAC | AA | ATTACATAGGTCAAACTGGCCGGTC | GCGCATCCGTTGATGAACATCAGTG | AT | TCTCACCATATTCGCTTC |
| 8 | CCTCAACCTACCTCCAAC | AA | AAAAACCCATCCTACACTCATTATA | CAAAGGCATCAAAGCCATGATCAGG | AT | TCTCACCATATTCGCTTC |
| 9 | CCTCAACCTACCTCCAAC | AA | TGTAAACGGATAATTCTGAGCTGAG | TGCGCTCCAAAGTGATCACAGCCAA | AT | TCTCACCATATTCGCTTC |
| 10 | CCTCAACCTACCTCCAAC | AA | TCGGCCGCGGCCAGATTACAGACTA | ATACCGAGGTAGATGCCCATGAAGA | AT | TCTCACCATATTCGCTTC |
| 11 | CCTCAACCTACCTCCAAC | AA | CAGCGCAGAGTCCACCAATCGAAGA | GACAACAGAAAAACCACCCACACTC | AT | TCTCACCATATTCGCTTC |
| 12 | CCTCAACCTACCTCCAAC | AA | CTCCTCCATGTAGCTGGTGGCGATT | ACCACTCACATCGTGCTCCTCGAAG | AT | TCTCACCATATTCGCTTC |
| 13 | CCTCAACCTACCTCCAAC | AA | TGCTGAGATTATGCATCTGCGGCCA | GATCATGCATGGAAGCACCCAACTG | AT | TCTCACCATATTCGCTTC |
| 14 | CCTCAACCTACCTCCAAC | AA | GTGGGAGAAACGCGCAGCAGTGGTA | TTTTCTTCTGGGACGACATGGCAAC | AT | TCTCACCATATTCGCTTC |
| 15 | CCTCAACCTACCTCCAAC | AA | GGGAAAGGTGTCCGGTGGTGGAAAC | ATAGGAGAGTATAAGAGTTTGGATC | AT | TCTCACCATATTCGCTTC |
| 16 | CCTCAACCTACCTCCAAC | AA | AGGTGTAGCAAAGCGCGCAGTCCCG | CTAAGTTTTGGATTGTTGAAGGTCT | AT | TCTCACCATATTCGCTTC |
| 17 | CCTCAACCTACCTCCAAC | AA | CCAAATTGAGATCCTCGAGGGCTGT | CGGGCAACTCTGGAAAAATATTGTT | AT | TCTCACCATATTCGCTTC |
| 18 | CCTCAACCTACCTCCAAC | AA | CGAGATCTCGTTGCCCTCCAGGTCA | ACCGGAGAAGGCTTCCTTGTGTATG | AT | TCTCACCATATTCGCTTC |
| 19 | CCTCAACCTACCTCCAAC | AA | GCATCTTGTGGCAGTGCCTTGATTC | AGTTGGAGCTTCGGAATGCCCTGAA | AT | TCTCACCATATTCGCTTC |
| 20 | CCTCAACCTACCTCCAAC | AA | GCTTCAATCCATTGAAGGGCTTTCC | TGTAGGAGAGCAACAGATCGTTCAA | AT | TCTCACCATATTCGCTTC |
| 21 | CCTCAACCTACCTCCAAC | AA | ATCTAACAGCCTTAAGTCTCTGCAG | AATCTTTTCAATTTGATTGCTCGAT | AT | TCTCACCATATTCGCTTC |
| 22 | CCTCAACCTACCTCCAAC | AA | TTTAGTTCTAAACTCTTCAGTCTTG | TTTGGGATGCGTTTTAGGGAATTGG | AT | TCTCACCATATTCGCTTC |
| 23 | CCTCAACCTACCTCCAAC | AA | TGCCGGCACGATCCAGTTTCAGTAT | GGCATAGATTCGCTGGCACCTCCTG | AT | TCTCACCATATTCGCTTC |
| 24 | CCTCAACCTACCTCCAAC | AA | AGCGCAAAGTTCGCACATCCGATAA | CATGGCAGGCCTCAAGTTCGGGAAA | AT | TCTCACCATATTCGCTTC |
| 25 | CCTCAACCTACCTCCAAC | AA | TGGATTGGCCCTCATGTCCAGCATT | AAACGCCCCTGCCGAAATTGTGGAC | AT | TCTCACCATATTCGCTTC |
| 26 | CCTCAACCTACCTCCAAC | AA | CGCAGTCGATTGCTGGTGATGGAAA | CTGGGCAGTTCCGTGTCGTTTATCC | AT | TCTCACCATATTCGCTTC |
| 27 | CCTCAACCTACCTCCAAC | AA | CCTCGGGCAGGCTGCTTATTAAATT | CCTGCAGTTGCGACAATTTGCTGAG | AT | TCTCACCATATTCGCTTC |
| 28 | CCTCAACCTACCTCCAAC | AA | TAAATTTTTAAGAGCTCCCGCGGAG | GTCCAGCTCTAAAACTTTTAAGGCT | AT | TCTCACCATATTCGCTTC |
| 29 | CCTCAACCTACCTCCAAC | AA | AGCACGATCAAGTTTGGCATTCGCG | TTCATGATTTGATTTCGCTTCAGCA | AT | TCTCACCATATTCGCTTC |
| 30 | CCTCAACCTACCTCCAAC | AA | AATTGCTGCCCAAATTGAGTGCTTC | AGTCCTCGTCGTTTATTATTGTCAA | AT | TCTCACCATATTCGCTTC |
| 31 | CCTCAACCTACCTCCAAC | AA | GGAATGCGATAGAAGAGATTGCCCT | GTTCTGAGTCCGGCCAAGGCATTCG | AT | TCTCACCATATTCGCTTC |
| 32 | CCTCAACCTACCTCCAAC | AA | AGTTCTGCAGGCTCAATCGTTTCAA | ACTGCGGTGGCAAGGACTTGAGGCC | AT | TCTCACCATATTCGCTTC |
| 33 | CCTCAACCTACCTCCAAC | AA | CATATTAATGATGCTATTGTCGGAC | GGCCAGGCCATAGAATGCATTGGGA | AT | TCTCACCATATTCGCTTC |
| 34 | CCTCAACCTACCTCCAAC | AA | AAGTTGTTGAGGACGACCACCTCGT | AAAAATGAGTTCGCCTCCAATTTGG | AT | TCTCACCATATTCGCTTC |
| 35 | CCTCAACCTACCTCCAAC | AA | CACAAATGGAGTGCAGCGTTCGCTT | AGTGACAGTGCTTCTGCAACCACCG | AT | TCTCACCATATTCGCTTC |
| 36 | CCTCAACCTACCTCCAAC | AA | GGTATTCCTGCAAGGACTTCTCGTA | TTGAGGCGGTGGAGAGCTGACTGCC | AT | TCTCACCATATTCGCTTC |
| 37 | CCTCAACCTACCTCCAAC | AA | CCGTAGGAGTGACACTGGCTGATTT | ATGGCAGCATCACGTTCTCCCTGAT | AT | TCTCACCATATTCGCTTC |
| 38 | CCTCAACCTACCTCCAAC | AA | TTGTTGGACGGTTTGTGGATTTTGC | GGTTCTATTCACGGTAATTTGGCTA | AT | TCTCACCATATTCGCTTC |
| 39 | CCTCAACCTACCTCCAAC | AA | ACATATGGCTCTTGCTCCTGGCTAT | TGCTGCTGCTGCAGATGTTGCAGGT | AT | TCTCACCATATTCGCTTC |
| 40 | CCTCAACCTACCTCCAAC | AA | TTTCATCATGGCCCATTGCCGATGG | GCATATCCGGAGCGGGATTTTCTTT | AT | TCTCACCATATTCGCTTC |

#### Send1

| Pair | Initiator | Spacer | Hybridzation | Hybridzation | Spacer | Initiator |
|---|---|---|---|---|---|---|
| 1 | GAGGAGGGCAGCAAACGG | AA | TTGCTGCGAACGTTAAGTTGGAATG | AACTGGATTTATGAACAACAGTAAT | TA | GAAGAGTCTTCCTTTACG |
| 2 | GAGGAGGGCAGCAAACGG | AA | GAAAATGCAATACATCGATGGCACT | TAAATTACACATTTGTAGCGGGTGC | TA | GAAGAGTCTTCCTTTACG |
| 3 | GAGGAGGGCAGCAAACGG | AA | AGCGTACAGAGAAGATCCCAAGCAA | GATCCAGTTACGATAACGCGCCACA | TA | GAAGAGTCTTCCTTTACG |
| 4 | GAGGAGGGCAGCAAACGG | AA | ACTAGCTGGCCGTTGAAGACCAAGG | ATGTTTCCCGTTCTCGATGTGATCC | TA | GAAGAGTCTTCCTTTACG |
| 5 | GAGGAGGGCAGCAAACGG | AA | CATTCTTCCCGCGCAGATGTCCTCA | GGAATCTCCATAACAGCCGCCTTTT | TA | GAAGAGTCTTCCTTTACG |
| 6 | GAGGAGGGCAGCAAACGG | AA | CACACGATCAGCGGCCGAATAAGGA | AACACACCGTTGCCATATTTCAATT | TA | GAAGAGTCTTCCTTTACG |
| 7 | GAGGAGGGCAGCAAACGG | AA | GGCCAAGGGAATTGTTTGTATGGAA | CGAAGACGACGTGGGAGGATCCGTC | TA | GAAGAGTCTTCCTTTACG |
| 8 | GAGGAGGGCAGCAAACGG | AA | AGGACAGCCACGTCGTTCTCCATGT | GAAAAACTTAAAGGACTGCTCAGTT | TA | GAAGAGTCTTCCTTTACG |
| 9 | GAGGAGGGCAGCAAACGG | AA | ATGCTTCCACTCCCACCACAACTCC | GTTTATCAAATTGTGGGTGAATGAT | TA | GAAGAGTCTTCCTTTACG |
| 10 | GAGGAGGGCAGCAAACGG | AA | AATTGAGCGTTCACCTTCTTTAATA | GCTATCATGCAGCGAGGATCCTGCT | TA | GAAGAGTCTTCCTTTACG |
| 11 | GAGGAGGGCAGCAAACGG | AA | CTGTAGATGGATCCGCCGCAAAAAT | TGGGCAGCTGTTATAATTATGGTTT | TA | GAAGAGTCTTCCTTTACG |
| 12 | GAGGAGGGCAGCAAACGG | AA | ACGGAACATCAGTGATGTCCATGCT | CGCCGTAGTACTGTAAGGACACCTG | TA | GAAGAGTCTTCCTTTACG |
| 13 | GAGGAGGGCAGCAAACGG | AA | AAAGGTCAGGTACCACCGGAGAAAT | CGCCGATGATGCGTTCAGATGGCTC | TA | GAAGAGTCTTCCTTTACG |

### SLO2

| Pair | Initiator | Spacer | Hybridzation | Hybridzation | Spacer | Initiator |
|---|---|---|---|---|---|---|
| 1 | CCTCGTAAATCCTCATCA | AA | TGAGTTTATTGTGAGTTCTGCTCGG | GGCTCATATTCATTTCCAACTATAT | AA | ATCATCCAGTAAACCGCC |
| 2 | CCTCGTAAATCCTCATCA | AA | TATGTGGCCTATGTGGTTGCAGATG | TTTGTACATTGAGACATCGATTTGA | AA | ATCATCCAGTAAACCGCC |
| 3 | CCTCGTAAATCCTCATCA | AA | TTTGTGAGTGTGAATGTTTGTGTGG | ACTGACGTAAGTGCTTTAACTAACA | AA | ATCATCCAGTAAACCGCC |
| 4 | CCTCGTAAATCCTCATCA | AA | GAATTTGGTACAGTTTCCGATACTT | TGCGAACGCTGGTTCACATAGCTCA | AA | ATCATCCAGTAAACCGCC |
| 5 | CCTCGTAAATCCTCATCA | AA | TGTATCTATATGGGGTTATACGTGG | ACAAGACATTGATAGCACCTTCATA | AA | ATCATCCAGTAAACCGCC |
| 6 | CCTCGTAAATCCTCATCA | AA | CATTCGATATATACAGTTATACAGT | GTTTGCTACACCTGGCTTAGAACTA | AA | ATCATCCAGTAAACCGCC |
| 7 | CCTCGTAAATCCTCATCA | AA | AGCGATAAACTGATAAGCACGCGCC | GCAATATACAATATAAGCCTACTTC | AA | ATCATCCAGTAAACCGCC |
| 8 | CCTCGTAAATCCTCATCA | AA | CTCTACGGGTGTGTCTGTGTGTGCT | AATACTTTCGCTCTGCCTAGAACTT | AA | ATCATCCAGTAAACCGCC |
| 9 | CCTCGTAAATCCTCATCA | AA | CGCCTAACATACATGGCACACCTAA | ATGGGTTAATGCCACCTCCACCGCC | AA | ATCATCCAGTAAACCGCC |
| 10 | CCTCGTAAATCCTCATCA | AA | CTGCATGAGGGTAGTGAAGTGACTT | CTTAATGGGATTACTATGGTTCGTT | AA | ATCATCCAGTAAACCGCC |
| 11 | CCTCGTAAATCCTCATCA | AA | GACTGTCGGGTAGAGAAAGAGAATT | GTCCTCGCTGATTGCCAACCACCGT | AA | ATCATCCAGTAAACCGCC |
| 12 | CCTCGTAAATCCTCATCA | AA | GTTCGACTTGCGGCGCGAGTTGTGT | CAGGTTGATGTTCGAGCAGAACGAG | AA | ATCATCCAGTAAACCGCC |
| 13 | CCTCGTAAATCCTCATCA | AA | GGTTGATTATCACATAGGACAGGTG | CTTCCTCCAGTTTGAGATCACAGCT | AA | ATCATCCAGTAAACCGCC |
| 14 | CCTCGTAAATCCTCATCA | AA | TTCGAGGTGTCCTGCGTGCGGTAAA | GAGTTACTCACATGAGACGTGTCCG | AA | ATCATCCAGTAAACCGCC |
| 15 | CCTCGTAAATCCTCATCA | AA | CGCACAAAGGTAATCACATAGTCCT | GGCGCCTGGTCAATGCCAAGCAGCA | AA | ATCATCCAGTAAACCGCC |
| 16 | CCTCGTAAATCCTCATCA | AA | GCATACTTGTCGTGGGCCCGGAACT | TCCTTCTCCATTTTGCTCAGATGGA | AA | ATCATCCAGTAAACCGCC |
| 17 | CCTCGTAAATCCTCATCA | AA | TATGGTGTTGCAGTCTGAGAGGGAG | GAATTTGAACATATTCTGCACGGCC | AA | ATCATCCAGTAAACCGCC |
| 18 | CCTCGTAAATCCTCATCA | AA | CCCAGCATCCAGTAGACCAAAGGAA | AGCAGATCGTCCAGGCAGTCGATCG | AA | ATCATCCAGTAAACCGCC |
| 19 | CCTCGTAAATCCTCATCA | AA | TGAGCTCTCAGCGGGATGATGAAGT | GGATTCAGCGAGGTCTTCGATCGGA | AA | ATCATCCAGTAAACCGCC |
| 20 | CCTCGTAAATCCTCATCA | AA | AACAGCAGAGTGGCTTCTTCTCTTT | AGTGCTCGCAGACCTGGGCCAACTG | AA | ATCATCCAGTAAACCGCC |
| 21 | CCTCGTAAATCCTCATCA | AA | CATCTCGTCGTCGATCTCATCATCA | AATCTTCTCGGATGGCGAACGCCAG | AA | ATCATCCAGTAAACCGCC |
| 22 | CCTCGTAAATCCTCATCA | AA | TTGGGATTGTTGGCAAAGGGCACAT | TGGTTGAGCACATCTGGGCTGAGCA | AA | ATCATCCAGTAAACCGCC |
| 23 | CCTCGTAAATCCTCATCA | AA | TCCGATGTTTGATTTTGATTAACGA | CTCCCCTCCTTGGCAGCTGCCGTGG | AA | ATCATCCAGTAAACCGCC |
| 24 | CCTCGTAAATCCTCATCA | AA | TAGAATGGTTTCCTCGTAGAATTCC | CATAATGTGCCTGGGTCCGGGATTC | AA | ATCATCCAGTAAACCGCC |
| 25 | CCTCGTAAATCCTCATCA | AA | CCCAGGACGATGTGGTAGATCTCGT | TCGTACTCGCCAAAGAATCGACTGT | AA | ATCATCCAGTAAACCGCC |
| 26 | CCTCGTAAATCCTCATCA | AA | TCCTTGACGGCCCAGGAACGTAGAA | ACATACTGCGGTACATTCGGTGCGA | AA | ATCATCCAGTAAACCGCC |
| 27 | CCTCGTAAATCCTCATCA | AA | TCCTTTAGACATGATCCCTGAATGT | TTCATCCTGGCGCGCGCCAAGTCGC | AA | ATCATCCAGTAAACCGCC |
| 28 | CCTCGTAAATCCTCATCA | AA | TTTGGGCGCGATGTGAACTGTAACT | TGGAGCACACCACCACGTGCTTTTC | AA | ATCATCCAGTAAACCGCC |
| 29 | CCTCGTAAATCCTCATCA | AA | AATGTCCGGCACAAAGTCGCCGTAT | GATGACCATATAAAGCTGCGAGGGC | AA | ATCATCCAGTAAACCGCC |
| 30 | CCTCGTAAATCCTCATCA | AA | ACCACAAACGCTCGTGAAGACCAAA | ATGGCCAGCGCGCTGAAAGTGCTGG | AA | ATCATCCAGTAAACCGCC |
| 31 | CCTCGTAAATCCTCATCA | AA | CAGCGATCGCTTGGCCAGCCAGCAA | GCGATGGAGGTCATTAAACATGTTC | AA | ATCATCCAGTAAACCGCC |
| 32 | CCTCGTAAATCCTCATCA | AA | GTTGCCCTTATAGCCTAGATATGTG | GTGAAAGGAGAGTATCTGCTGCCAG | AA | ATCATCCAGTAAACCGCC |
| 33 | CCTCGTAAATCCTCATCA | AA | AACTCCTCCTCCGTCAGTTTGGCCG | TCCCAGTTGATGATTGGATTCTCCT | AA | ATCATCCAGTAAACCGCC |
| 34 | CCTCGTAAATCCTCATCA | AA | CGACAGTAACTTAAGGAATAAGTCG | TATCACACGTATTATGTAGAGAACA | AA | ATCATCCAGTAAACCGCC |
| 35 | CCTCGTAAATCCTCATCA | AA | ACTCCACACGAACTCTGGGGATCTG | CTTTGAATGTGTTTTCGTTCACATA | AA | ATCATCCAGTAAACCGCC |
| 36 | CCTCGTAAATCCTCATCA | AA | TGCAGTTGTGGTTATTATTGTTATC | TGTAGTTACGAATGCTAGTTAAACG | AA | ATCATCCAGTAAACCGCC |
| 37 | CCTCGTAAATCCTCATCA | AA | GCCAGGTTGGCTATATCGTGCTAGG | CGGAAATAGATGCACTCTCGCAGGA | AA | ATCATCCAGTAAACCGCC |
| 38 | CCTCGTAAATCCTCATCA | AA | AGGAAAGCTCCCAGAAAGGATTGTT | ATAGACAGATCTACATAAAGCACAG | AA | ATCATCCAGTAAACCGCC |
| 39 | CCTCGTAAATCCTCATCA | AA | TTTCTCGACACTAGTGTGTTGTATA | TCTGTAACTTGCCCGATCTAAATGC | AA | ATCATCCAGTAAACCGCC |
| 40 | CCTCGTAAATCCTCATCA | AA | TGGTTTTGCCTCTTTCCGCTCTTAT | ACACGAGAATTGACAGCAAATTCGA | AA | ATCATCCAGTAAACCGCC |

### SoxN

| Pair | Initiator | Spacer | Hybridzation | Hybridzation | Spacer | Initiator |
|---|---|---|---|---|---|---|
| 1 | CCTCGTAAATCCTCATCA | AA | TTGTTGGATTTGTTTGTTTTCGCTG | TATTATTTCCAGGTGCTTGGCAATG | AA | ATCATCCAGTAAACCGCC |
| 2 | CCTCGTAAATCCTCATCA | AA | ACGCTGCTTCTTGGTTTGTTTTTAC | TGCGCCATGTTTCACAATCACGTTT | AA | ATCATCCAGTAAACCGCC |
| 3 | CCTCGTAAATCCTCATCA | AA | TTTGTATTATGCAGTAGATGCTTGC | GGTTTACTAACATTTTCTCTGGTTG | AA | ATCATCCAGTAAACCGCC |
| 4 | CCTCGTAAATCCTCATCA | AA | TATTTTCATCGCCTCGCCACAACTG | TTGACATTTTTCTTTGCTTTTCTGA | AA | ATCATCCAGTAAACCGCC |
| 5 | CCTCGTAAATCCTCATCA | AA | CCGCGTCCCGTTGCTACCTTCAATT | TTACTCCTACTAAACTTGTGCTTGA | AA | ATCATCCAGTAAACCGCC |
| 6 | CCTCGTAAATCCTCATCA | AA | GTACAGCAAAAGGTGAGTGGAAATG | CGCACATTTAATGTCTGAGAATGTT | AA | ATCATCCAGTAAACCGCC |
| 7 | CCTCGTAAATCCTCATCA | AA | TGCCGTTGTCCCCGTAATTGGCAAC | TGCTAAACCAAACGAATTTGTTGTT | AA | ATCATCCAGTAAACCGCC |
| 8 | CCTCGTAAATCCTCATCA | AA | TATGCAACAAGCGGGAAATCAAGGG | TTATTTGTATCACACAATTGTTTTA | AA | ATCATCCAGTAAACCGCC |
| 9 | CCTCGTAAATCCTCATCA | AA | TGAGTTGTCATCTTACTATTGTGGT | TCGAAGCAGAACGGTATGGAATGCA | AA | ATCATCCAGTAAACCGCC |
| 10 | CCTCGTAAATCCTCATCA | AA | GTGTTACTCTGCCACTTCGAAATTC | CCTATTCTCATTTATTTATATGCGC | AA | ATCATCCAGTAAACCGCC |
| 11 | CCTCGTAAATCCTCATCA | AA | AATTGTGTCTGTGTGTGTGTGGCTG | TTAAACCTGCTTAAGAACTAAGGCT | AA | ATCATCCAGTAAACCGCC |
| 12 | CCTCGTAAATCCTCATCA | AA | GGCTTTAGTTAATATTTTCAATAAT | TTGTGGGCGTGGGTGTGCGTATCCT | AA | ATCATCCAGTAAACCGCC |
| 13 | CCTCGTAAATCCTCATCA | AA | TGTTGGTGCTGCTGCGAAAGGTGTT | TGCATGGTATGGTGTTGCTGCGGGA | AA | ATCATCCAGTAAACCGCC |
| 14 | CCTCGTAAATCCTCATCA | AA | TGCTCATCCGGCAGATGGTACATAT | TCAGTCTGATAGTGCAGCAGGTGGC | AA | ATCATCCAGTAAACCGCC |
| 15 | CCTCGTAAATCCTCATCA | AA | CCCGCTTCATGATGTGATTGTTGTT | CTGCTGCCGCCGCTGCAGAGCTGTA | AA | ATCATCCAGTAAACCGCC |
| 16 | CCTCGTAAATCCTCATCA | AA | ACATCGCATACCCGTTGTTGTACAT | ATGTCTGTCCGCCGGACACCGTCTG | AA | ATCATCCAGTAAACCGCC |
| 17 | CCTCGTAAATCCTCATCA | AA | GGCATGTATCCATTGGGCGCGTTCA | GGATCTGCGTGCATCATGTAGCCAT | AA | ATCATCCAGTAAACCGCC |
| 18 | CCTCGTAAATCCTCATCA | AA | CCTTGGTCTTGGTCAGCGTTTTAGT | GCATCAGGCCGCCCATTGGGTACTT | AA | ATCATCCAGTAAACCGCC |
| 19 | CCTCGTAAATCCTCATCA | AA | CCGCTTCACACGATCCGCGTGCTGT | TGACCAAACCATGAACGCATTCATA | AA | ATCATCCAGTAAACCGCC |
| 20 | CCTCGTAAATCCTCATCA | AA | TTGTTGTTGCTGGATCCTCCAGTTA | TTCTTGTTGGCCGTGGCGCTGTTGT | AA | ATCATCCAGTAAACCGCC |
| 21 | CCTCGTAAATCCTCATCA | AA | TCTGGGTGGGACTCAATTCGCTGGA | ATGTCATGTGATGGCTTCCGATGCT | AA | ATCATCCAGTAAACCGCC |
| 22 | CCTCGTAAATCCTCATCA | AA | TGGTTGTGATGGCTCAGCTGCGAGT | ATATGGGCGCTCATGTGGTGATGAT | AA | ATCATCCAGTAAACCGCC |
| 23 | CCTCGTAAATCCTCATCA | AA | TCATATCCGATTCCATGGTCAGCAT | GCATGGTGGCATGCAGCAGGCTGCC | AA | ATCATCCAGTAAACCGCC |
| 24 | CCTCGTAAATCCTCATCA | AA | GATAAAGCTGATTTGACTCAAAGTT | GCGGGGATTACTTCCAGTTTTCGGT | AA | ATCATCCAGTAAACCGCC |
| 25 | CCTCGTAAATCCTCATCA | AA | CAGTTACTCTTTAGCTTTAGTTTAC | TCTCCGTGGATACTCGCCTCTAAGA | AA | ATCATCCAGTAAACCGCC |
| 26 | CCTCGTAAATCCTCATCA | AA | TGTGGGAAACCATGTTGCACTTGTG | TTGCTTAGTTAGACATTTTGAACAT | AA | ATCATCCAGTAAACCGCC |
| 27 | CCTCGTAAATCCTCATCA | AA | TCTTTACGCGTATCAAAAATGTGGC | GCAGTTTGTATTTCGTAGTTTTTCG | AA | ATCATCCAGTAAACCGCC |
| 28 | CCTCGTAAATCCTCATCA | AA | GCTAGCAAACAAACAAGAGGCGGCG | AATATCAGATATACATATATGTTGT | AA | ATCATCCAGTAAACCGCC |
| 29 | CCTCGTAAATCCTCATCA | AA | GACGCGAAATTGCCTCAGAATATTG | AACACTTGAAAATTGTTGCTAAAAC | AA | ATCATCCAGTAAACCGCC |
| 30 | CCTCGTAAATCCTCATCA | AA | GAGCACCTTTCTGTTTCTCCGGACG | AAAAATACCTCCACACTGGGCGGAA | AA | ATCATCCAGTAAACCGCC |
| 31 | CCTCGTAAATCCTCATCA | AA | GCTGCTTTTTCCCTGGCTGTATGAT | GCGAAACCTGTTCGAGGTGCGTTCT | AA | ATCATCCAGTAAACCGCC |
| 32 | CCTCGTAAATCCTCATCA | AA | TTGGCTAGCTGATGACGAGGCACGA | TTTTTCTTTCTTTCGTATCACCGCT | AA | ATCATCCAGTAAACCGCC |
| 33 | CCTCGTAAATCCTCATCA | AA | ACGATACGGATACGATACGACACGA | AGAACCACCGATGCGATGCGATGCG | AA | ATCATCCAGTAAACCGCC |
| 34 | CCTCGTAAATCCTCATCA | AA | TGATGTTGCTGCGTGATGTTGCTGC | CGAATTCGAGACAGCGAACAACCGA | AA | ATCATCCAGTAAACCGCC |
