## Supplement for "Regional specialization, polyploidy, and seminal fluid transcripts in the Drosophila female reproductive tract"

### Supplementary Information: FACS methods

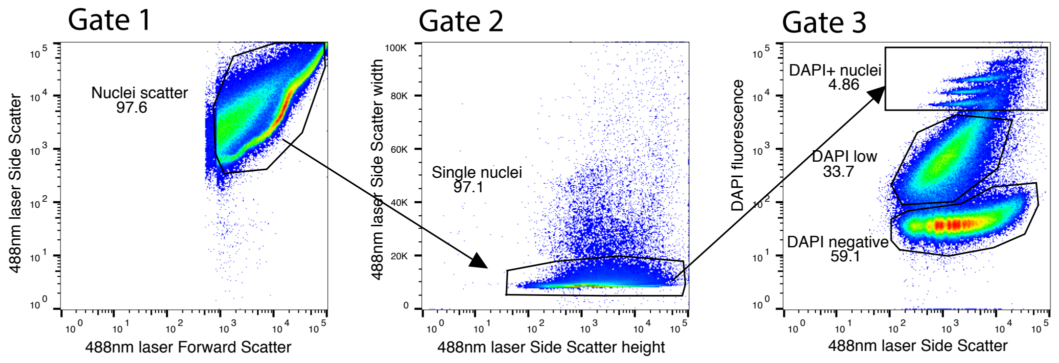


#### Back-gated positions of collected nuclei vs DAPI-low and DAPI-negative objects:

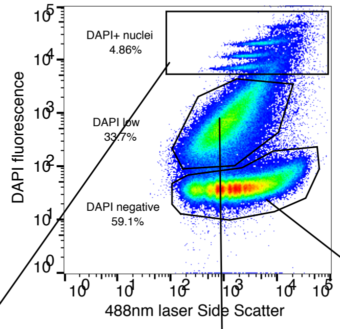


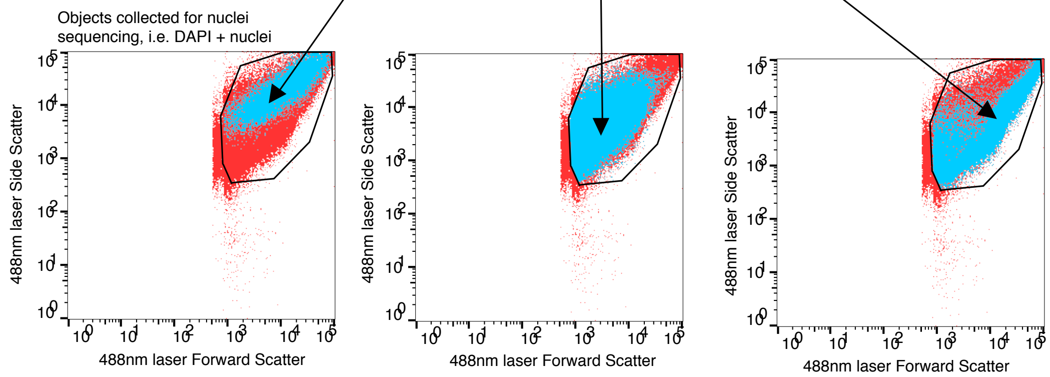


#### DAPI DNA-stain median fluorescent intensity (MFI) for nuclei subsets:

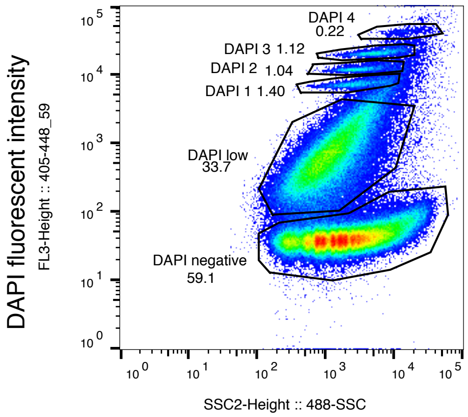


**Subset : MFI value , percent of all sorted objects belonging to this subset**

DAPI 1 : MFI : 7061, 1.4%
DAPI 2 : MFI : 11548, 1.04%
DAPI 3 : MFI : 19754, 1.12%
DAPI 4 : MFI : 40112, 0.22%
DAPI low : MFI : 571, 33.7%
DAPI negative : MFI : 39.0, 59.1%

Expression levels per cluster for genes targeted by in situ hybridization

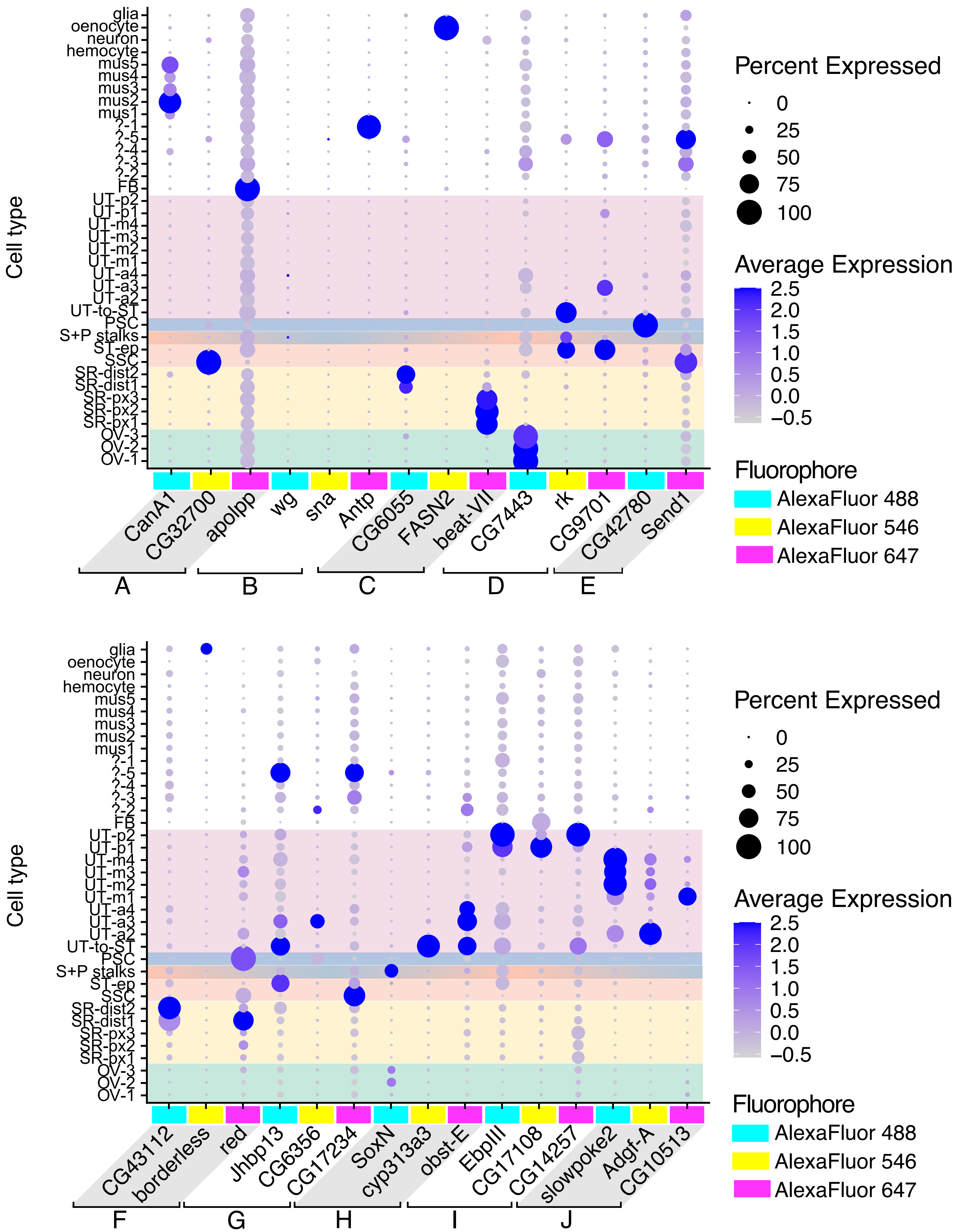

#### Pool A

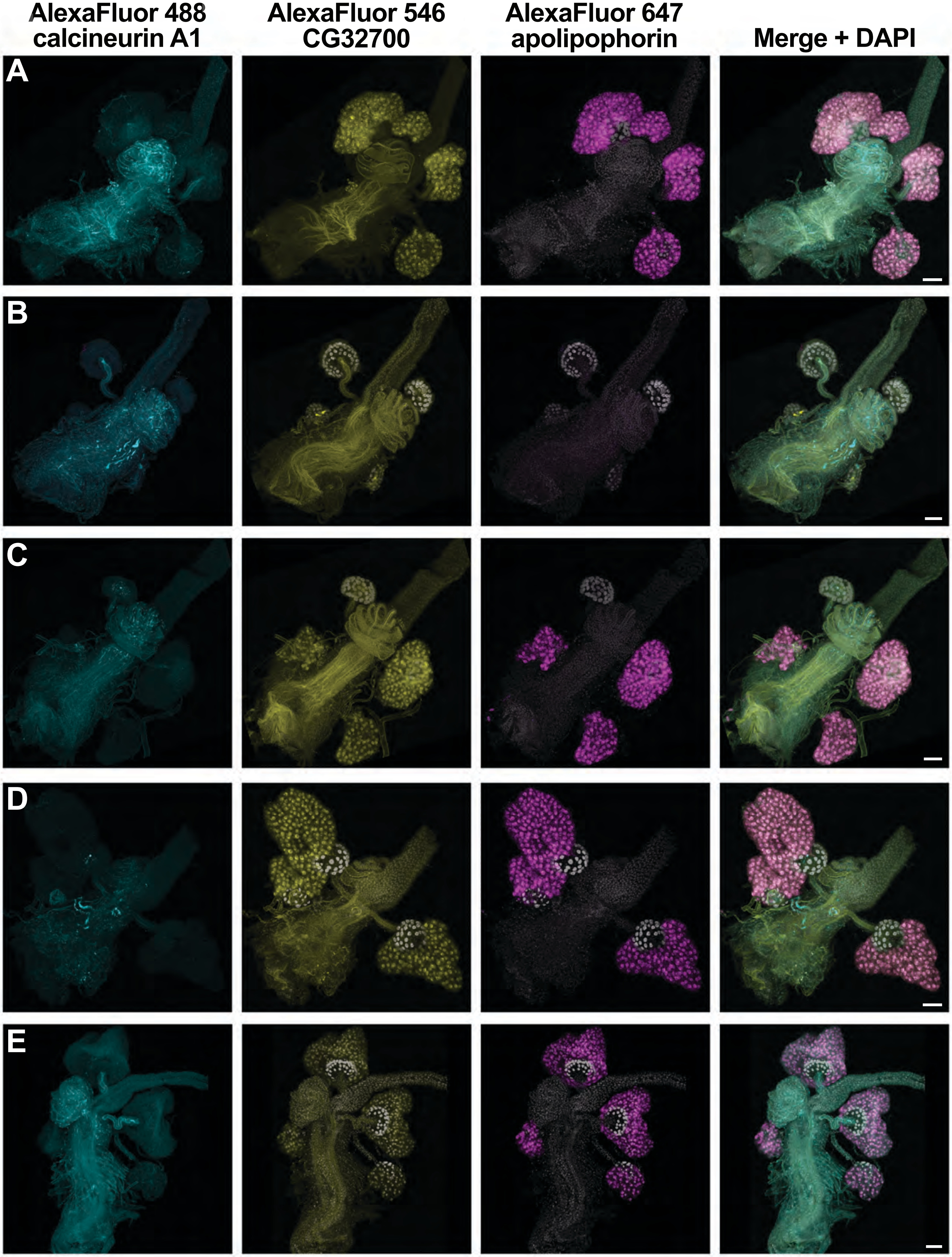
AlexaFluor 488
calcineurin A1
AlexaFluor 546
CG32700
AlexaFluor 647
apolipophorin
Merge + DAPI
A
B
C
D
E

### Pool B

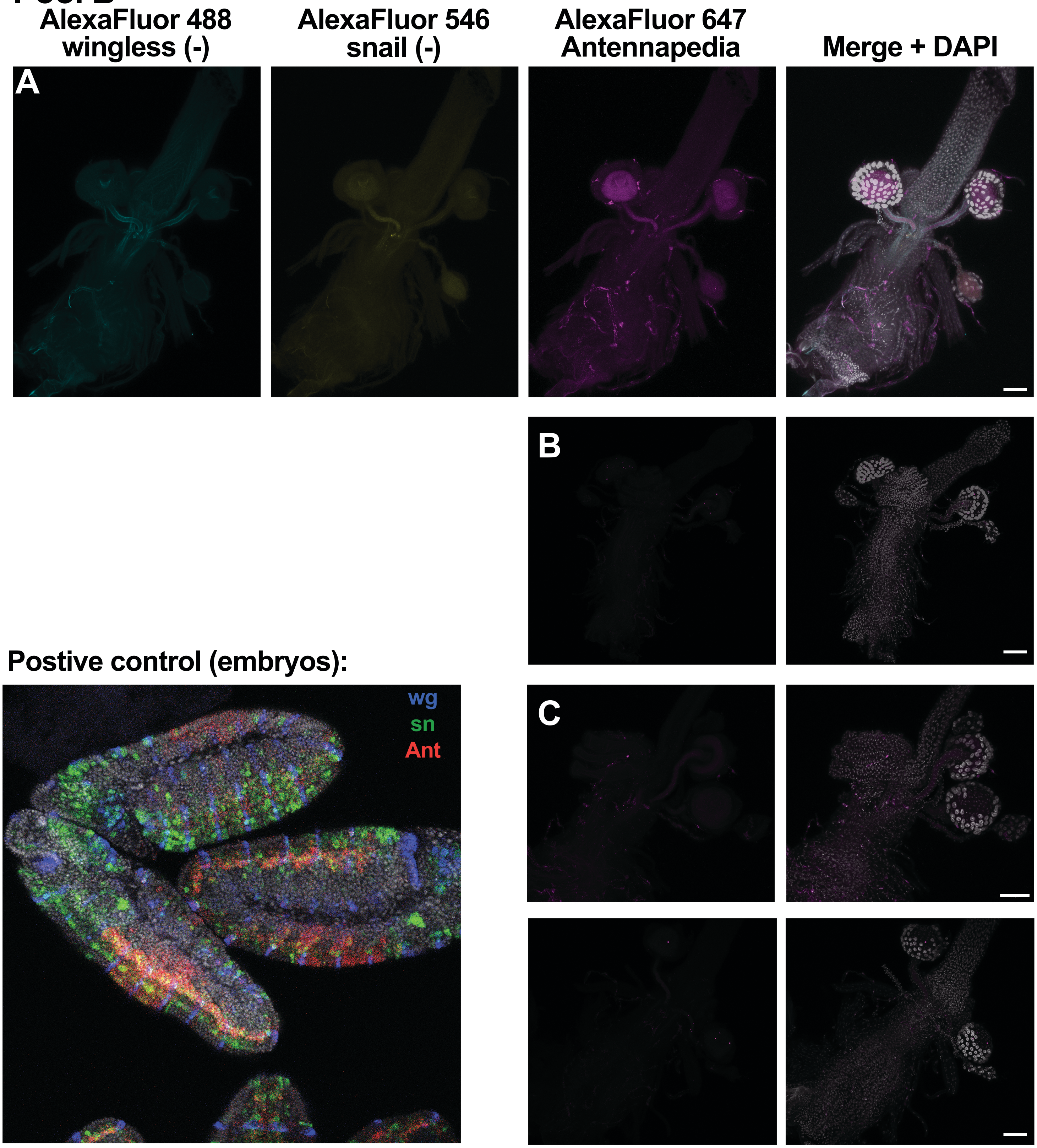
AlexaFluor 488
wingless (-)
AlexaFluor 546
snail (-)
AlexaFluor 647
Antennapedia
Merge + DAPI
A
B
C
Postive control (embryos):
wg
sn
Ant

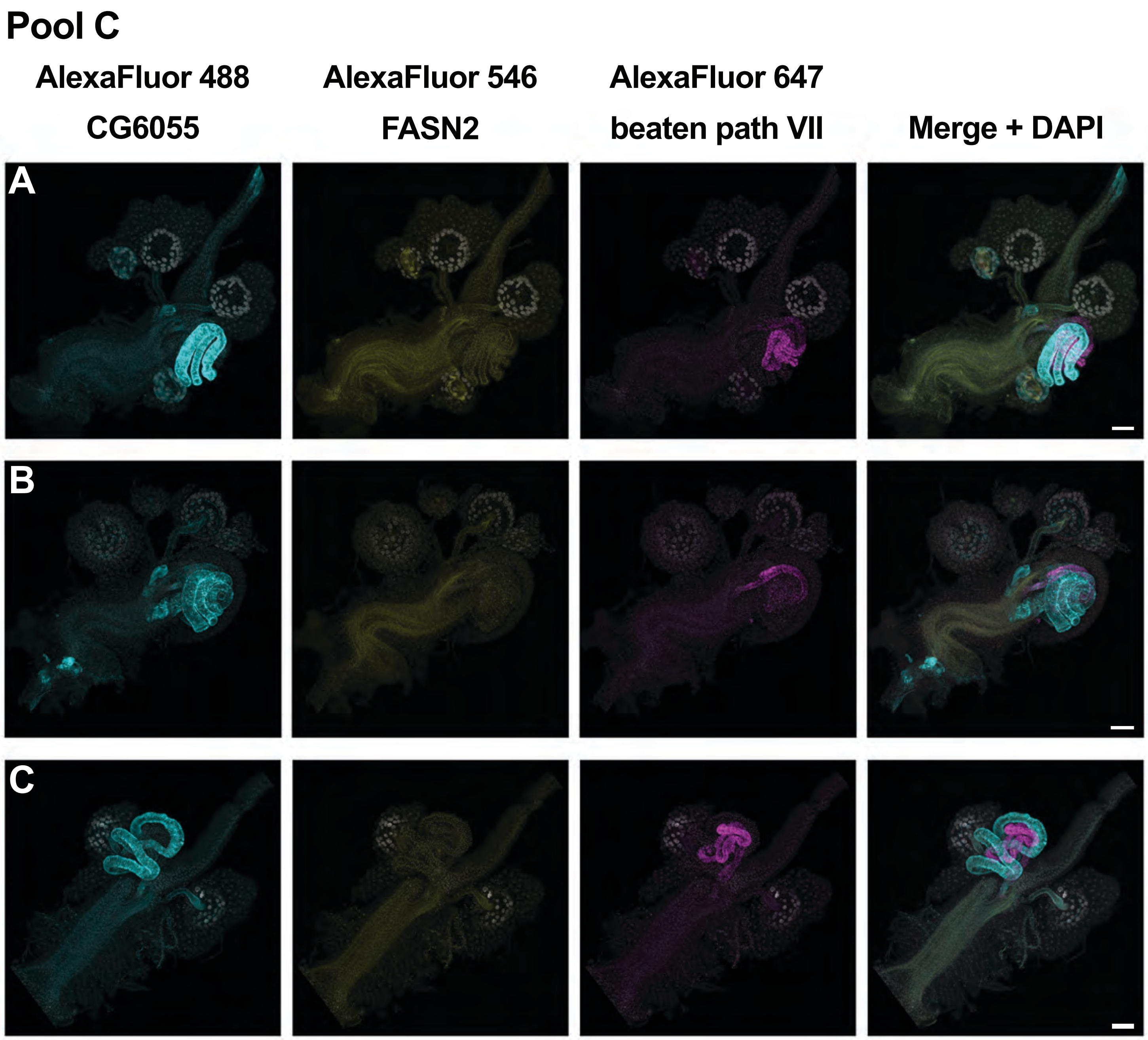
Pool C
AlexaFluor 488
CG6055
AlexaFluor 546
FASN2
AlexaFluor 647
beaten path VII
Merge + DAPI
A
B
C

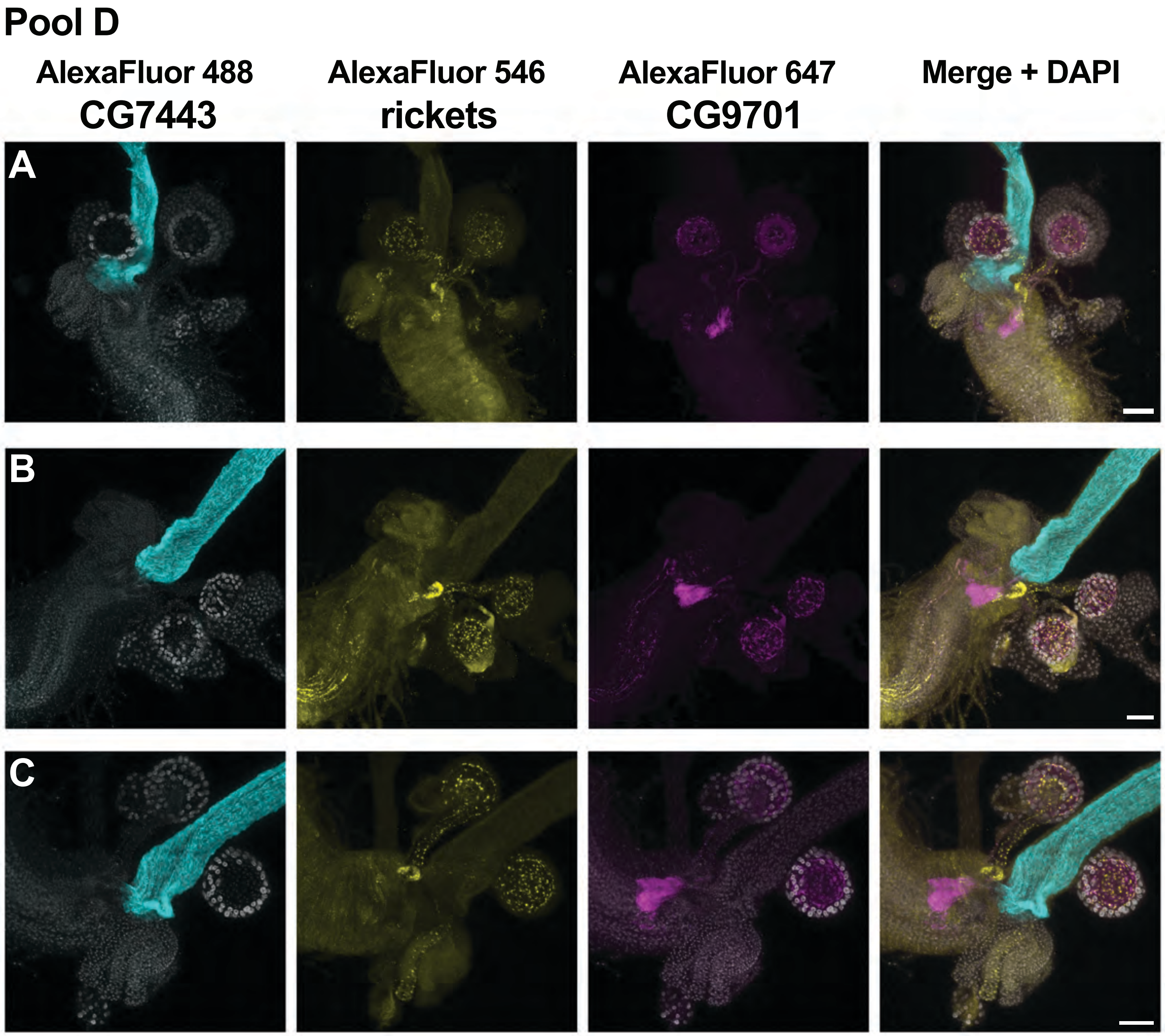
Pool D
AlexaFluor 488
CG7443
AlexaFluor 546
rickets
AlexaFluor 647
CG9701
Merge + DAPI
A
B
C

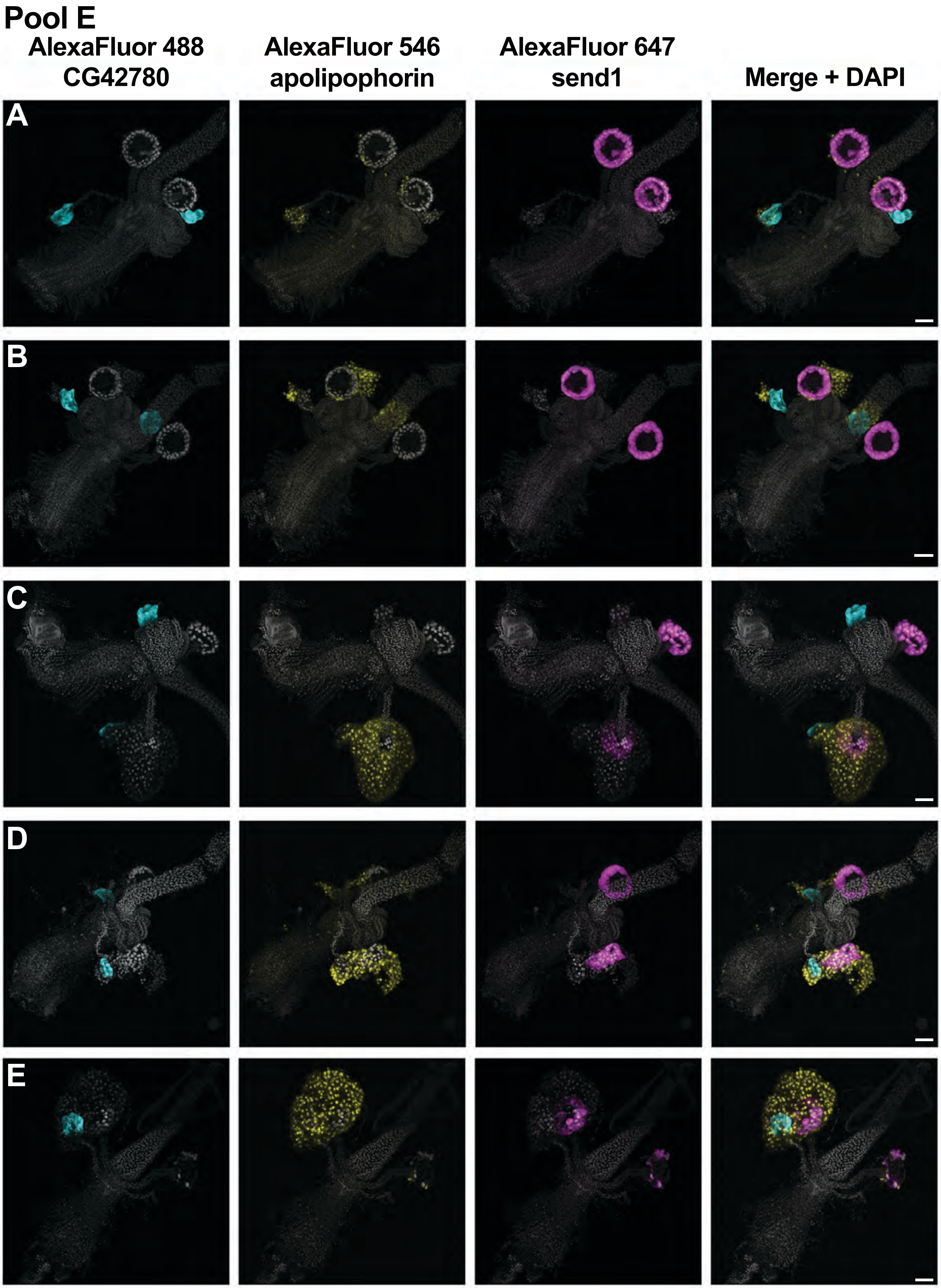
Pool E
AlexaFluor 488
CG42780
AlexaFluor 546
apolipophorin
AlexaFluor 647
send1
Merge + DAPI
A
B
C
D
E

### Pool F

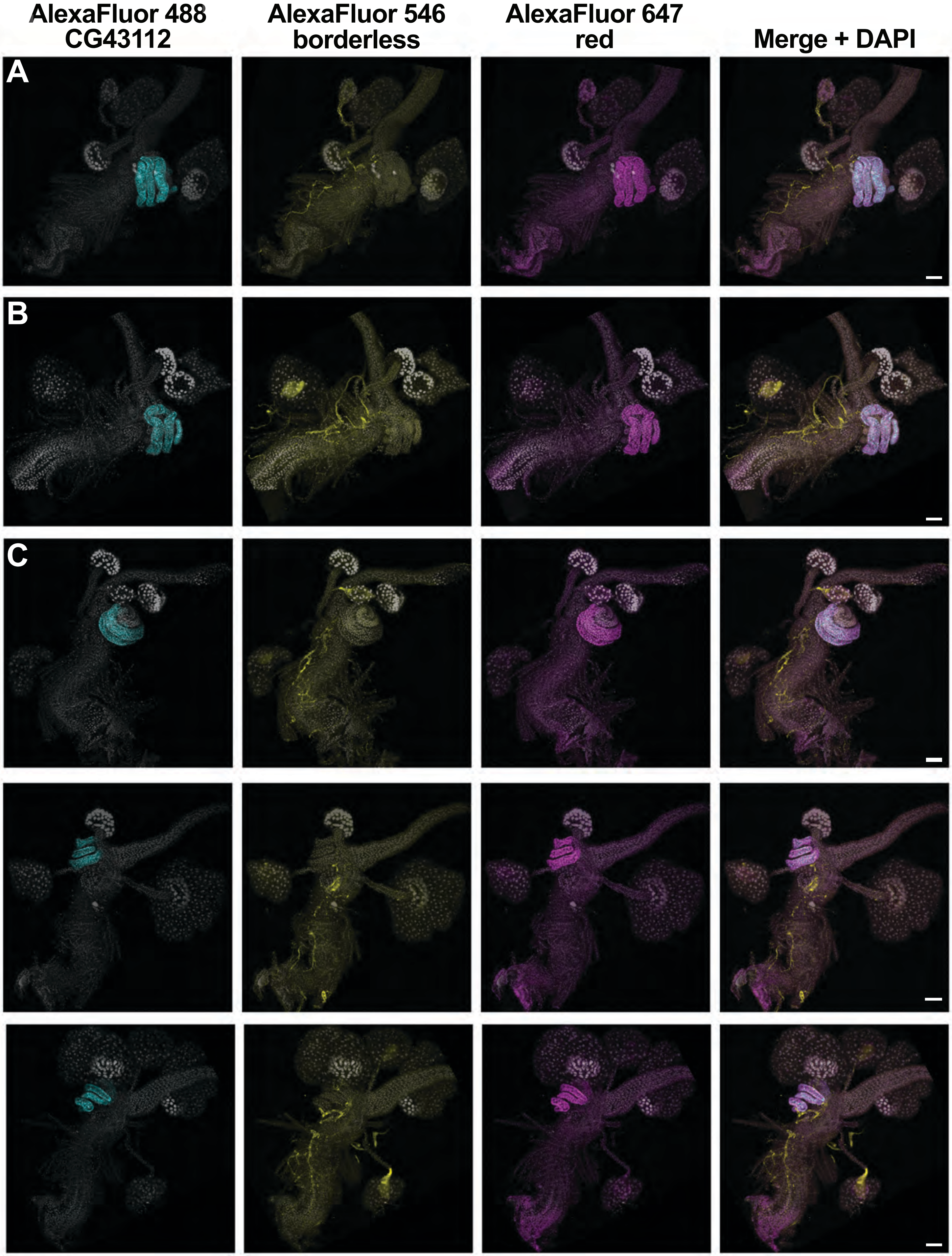

AlexaFluor 488
CG43112
AlexaFluor 546
borderless
AlexaFluor 647
red
Merge + DAPI
A
B
C

### Pool G

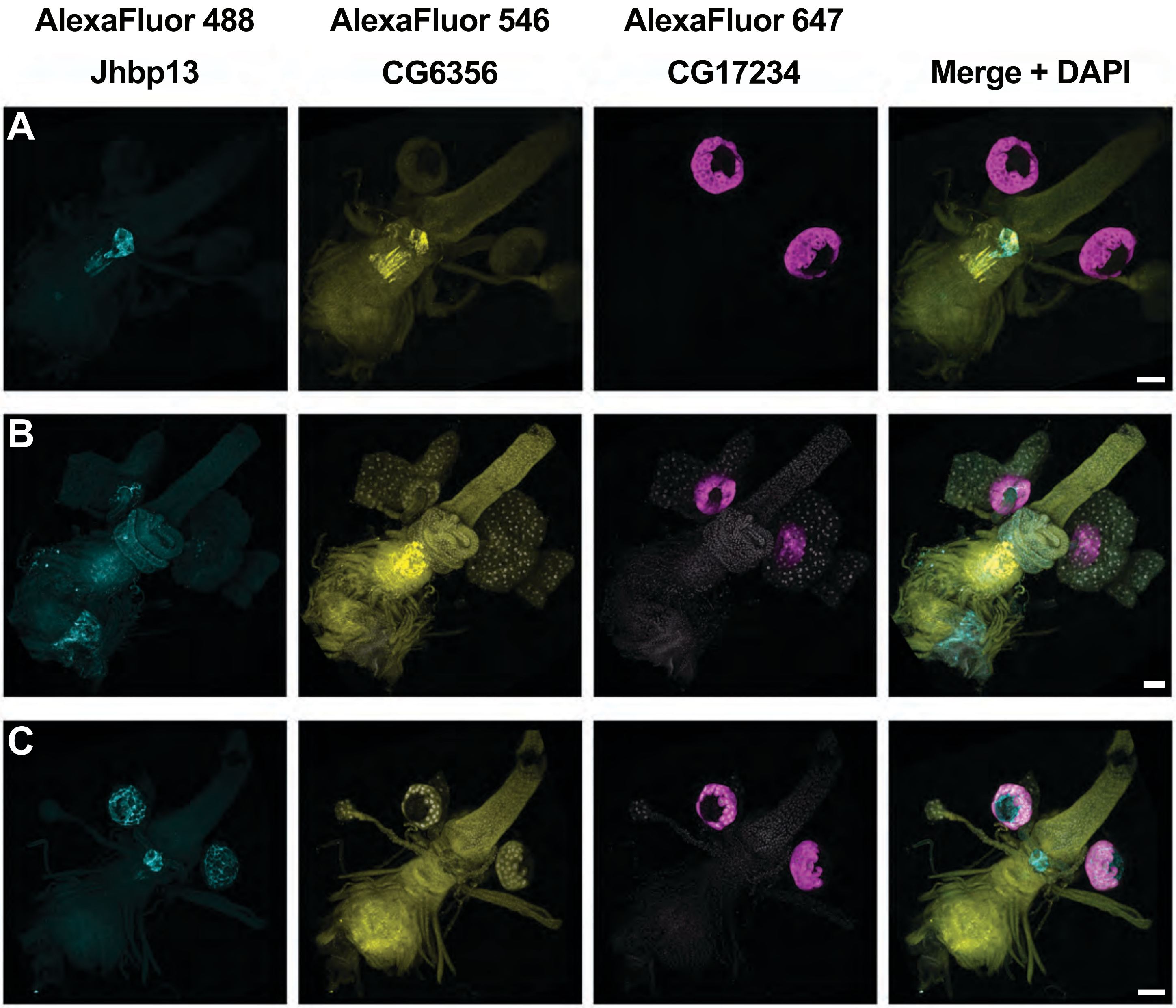

AlexaFluor 488
AlexaFluor 546
AlexaFluor 647
Jhbp13
CG6356
CG17234
Merge + DAPI
A
B
C

#### Pool H

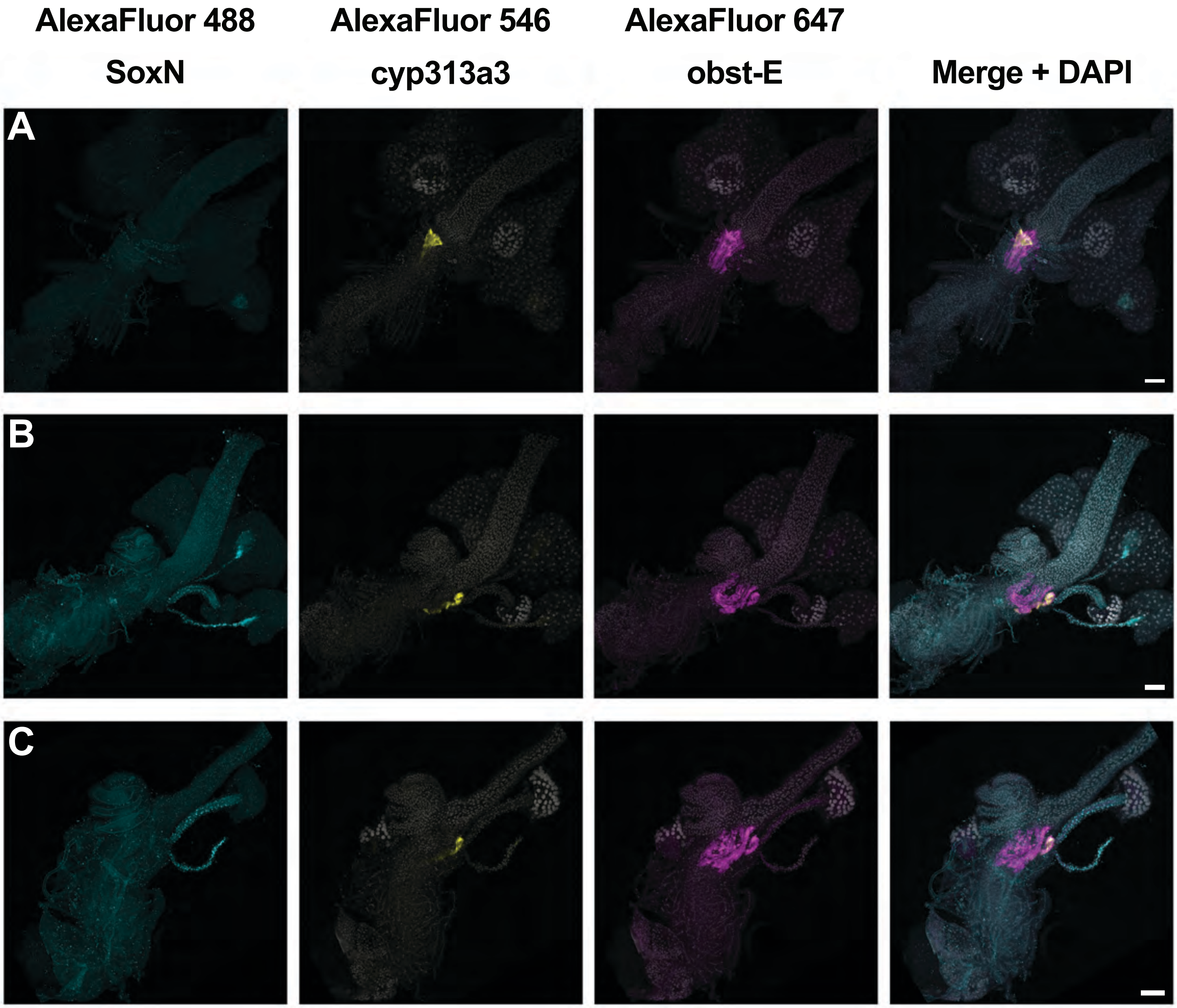
AlexaFluor 488
SoxN
AlexaFluor 546
cyp313a3
AlexaFluor 647
obst-E
Merge + DAPI
A
B
C

### Pool I

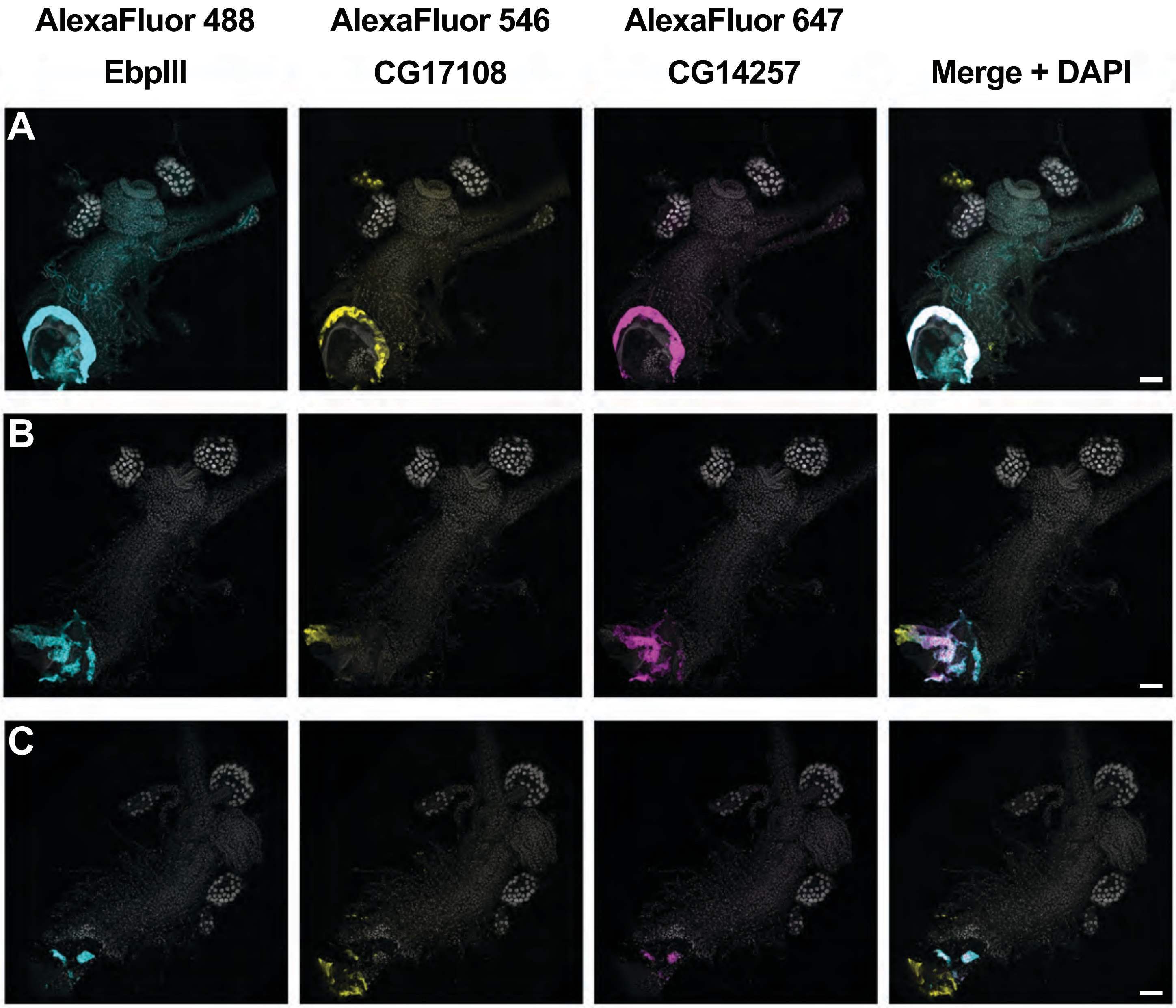

AlexaFluor 488
AlexaFluor 546
AlexaFluor 647
EbpIII
CG17108
CG14257
Merge + DAPI
A
B
C

### Pool J

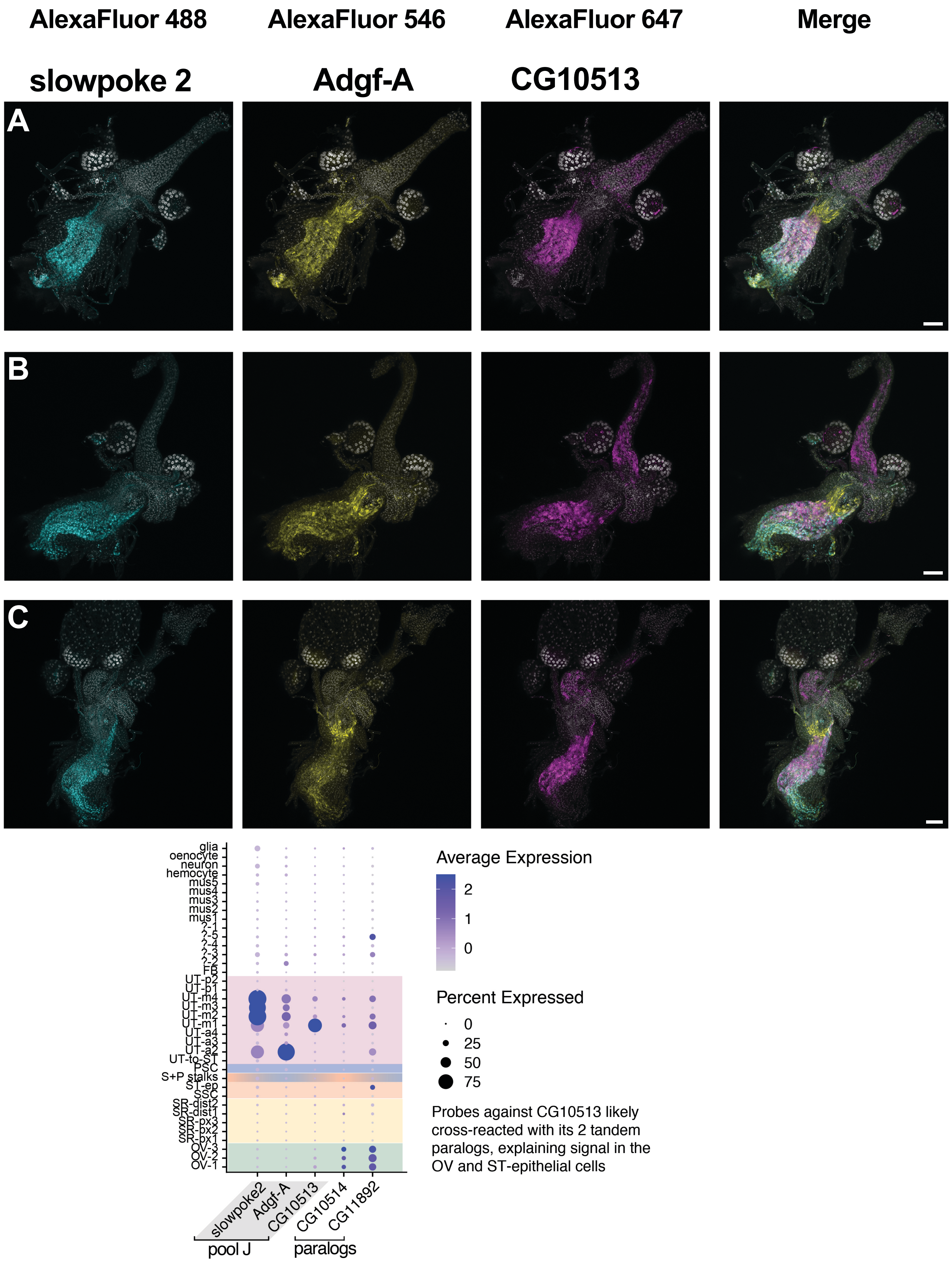
AlexaFluor 488
AlexaFluor 546
AlexaFluor 647
Merge
slowpoke 2
Adgf-A
CG10513
A
B
C
glia
oenocyte
neuron
hemocyte
mus5
mus4
mus3
mus2
mus1
?-1
?-5
?-4
?-3
?-2
FB
UT-p2
UT-p1
UT-m4
UT-m3
UT-m2
UT-m1
UT-a4
UT-a3
UT-a2
UT-to-ST
PSC
S+P stalks
ST-ep
SSC
SR-dist2
SR-dist1
SR-px3
SR-px2
SR-px1
OV-3
OV-2
OV-1
slowpoke2
Adgf-A
CG10513
CG10514
CG11892
pool J
paralogs
Average Expression
2
1
0
Percent Expressed
0
25
50
75
Probes against CG10513 likely cross-reacted with its 2 tandem paralogs, explaining signal in the OV and ST-epithelial cells

#### Negative controls: maximal laser power & gain

AlexaFluor 488 | AlexaFluor 546 | AlexaFluor 647 | Merge + DAPI

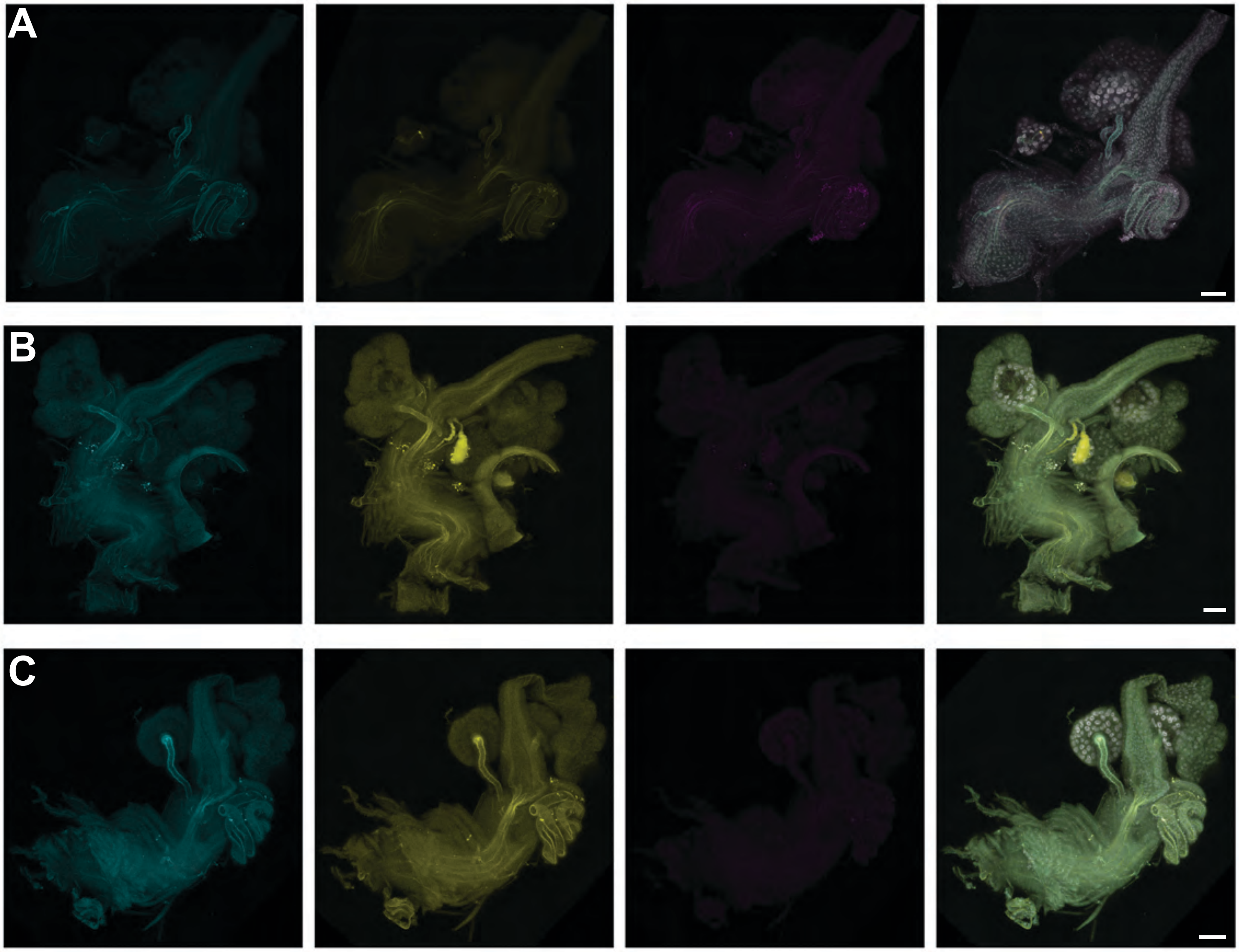


#### Negative controls: typical (mean) laser power & gain

AlexaFluor 488 | AlexaFluor 546 | AlexaFluor 647 | Merge + DAPI

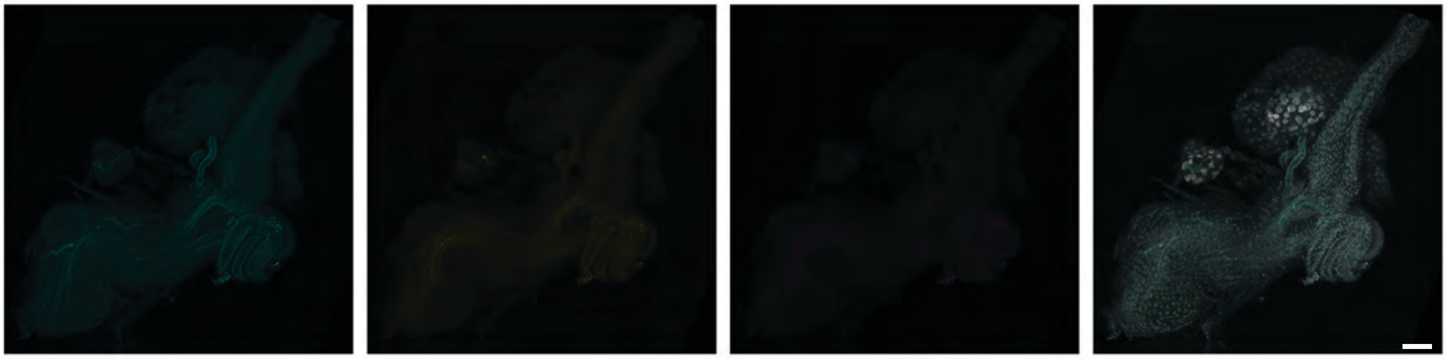

#### Negative controls: typical (mean) laser power & gain

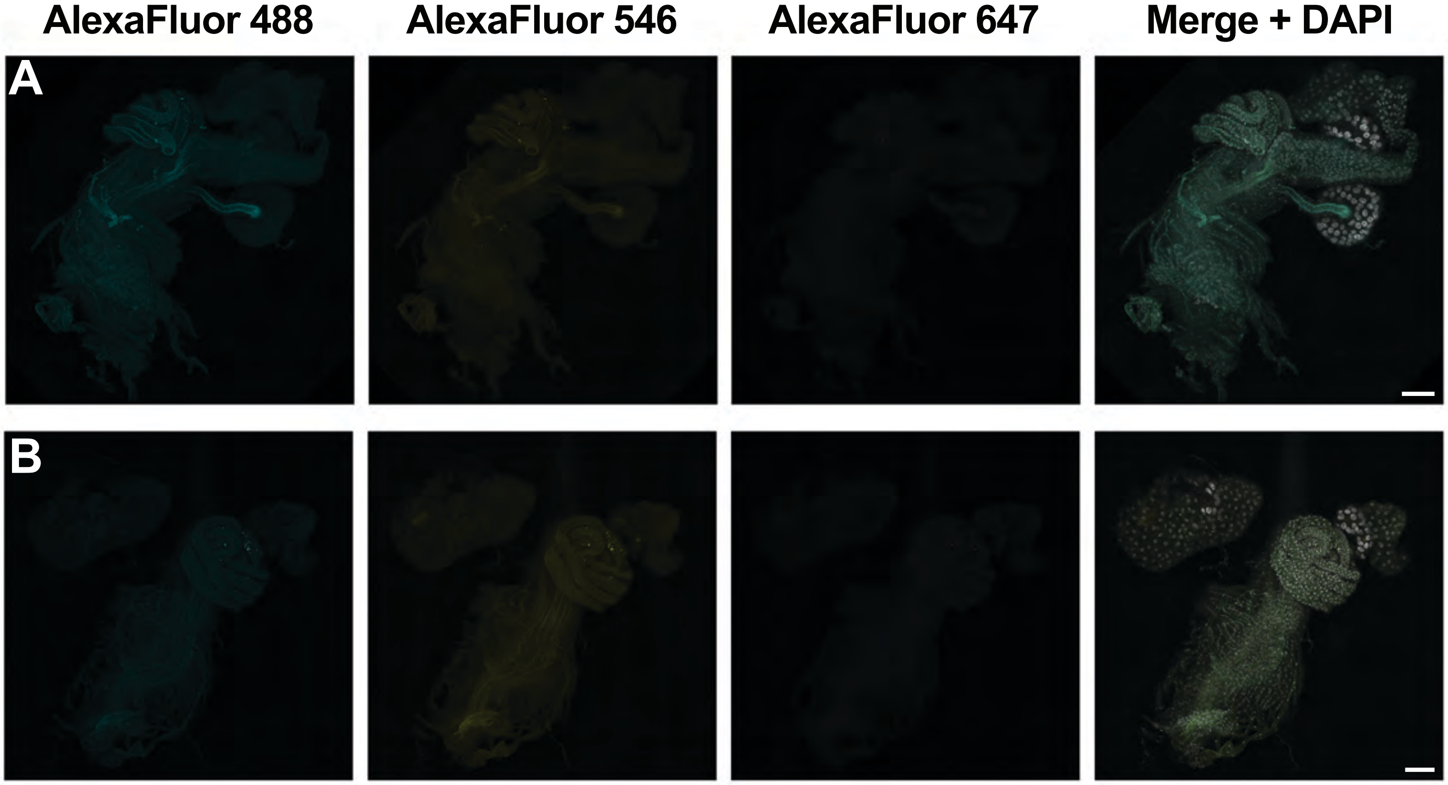


#### Negative controls: typical (median) laser power & gain

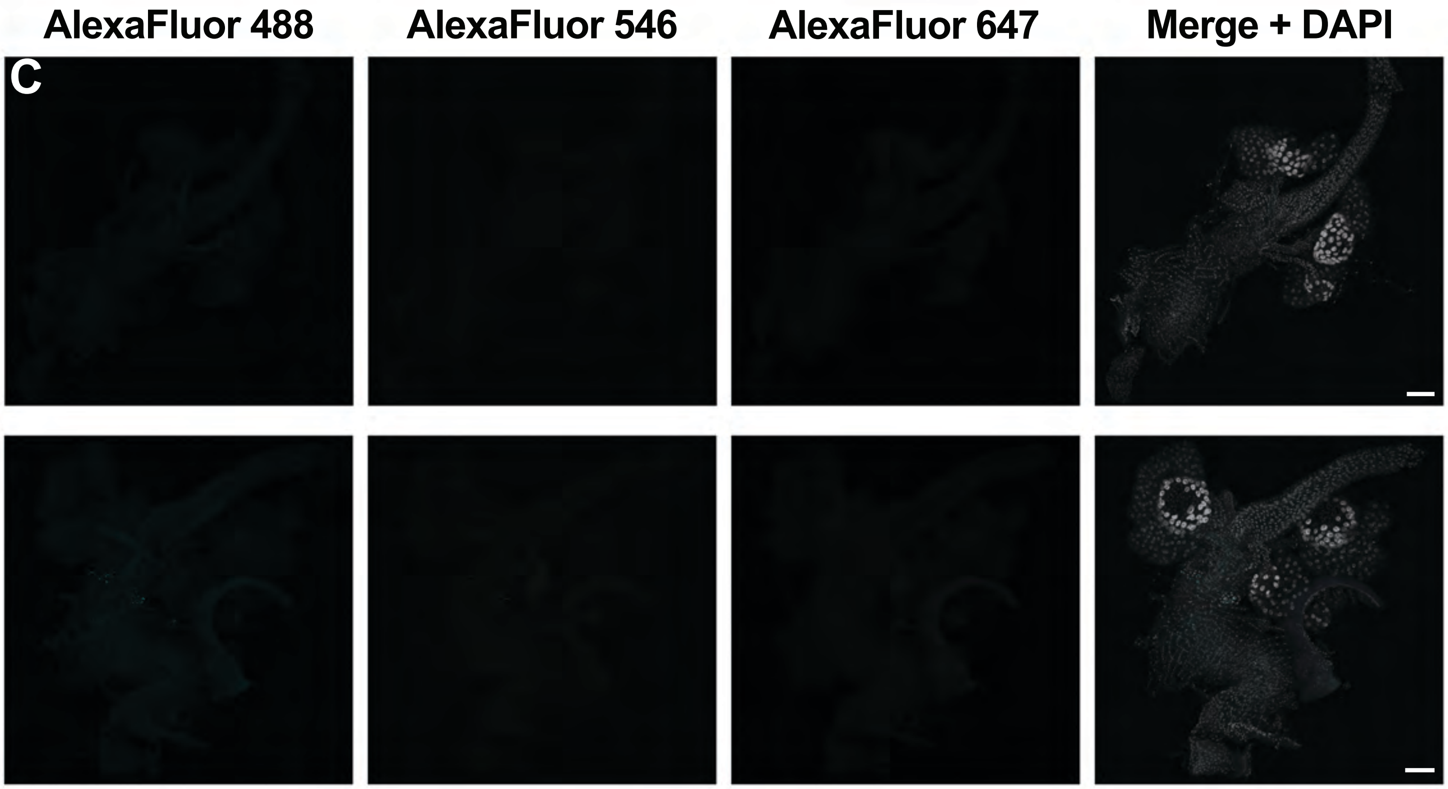
